# Impact of Yaq-001, a non-absorbable, engineered carbon bead of controlled porosity in rodent models of cirrhosis and acute on chronic liver failure

**DOI:** 10.1101/2023.08.15.553396

**Authors:** Jinxia Liu, Jane Macnaughtan, Yi Jin, Frederick Clasen, Abeba Habtesion, Alexandra Phillips, Francesco De Chiara, Ganesh Ingavle, Paul Cordero-Sanchez, Junpei Soeda, Jude A Oben, Jia Li, Haifeng Wu, Lindsey Ann Edwards, I. Jane Cox, Susan Sandeman, Nathan Davies, Rajeshwar Mookerjee, Gautam Mehta, Saeed Shoaie, Julian R. Marchesi, Fausto Andreola, Rajiv Jalan

**Author notes:** Both authors share first authorship (Jinxia Liu, Jane Macnaughtan). **Correspondence**: Rajiv Jalan, UCL Institute for Liver and Digestive Health, Upper third floor, Royal Free Campus, Rowland Hill Street, Hampstead, London, NW3 2PF. **Conflict of Interest**. Rajiv Jalan is the inventor of OPA, which has been patented by UCL and licensed to Mallinckrodt Pharma. He is also the founder of Yaqrit Discovery, Hepyx Limited (spin out companies from University College London), and Cyberliver. He has research collaborations with Yaqrit Discovery. Yaq-001 was licensed by Yaqrit Ltd. from UCL. JRM has received consultancy fees from EnteroBiotix and Cultech, and speaker fees from Falk Forum. **Financial support**. This study was performed with support from a grant from the EU H2020, Grant Agreement number: 634579 — CARBALIVE — H2020-PHC-2014-2015/H2020-PHC-2014 programme. JRM and the Division of Digestive Diseases at Imperial College London receives financial support from the NIHR Imperial Biomedical Research Centre. **Authors’ contributions**. RJ, FA, JL, JM, ND - contributed to the conception and design of the study. SS and GI contributed to the conception and design of the *in vitro* studies. RJ, ND, FA - provided administrative, study supervision, obtained funding, material support. JL, JM, LE, YJ, FC, AH, AP, FD, GI, PC, JS, JO, JL, HW, JC, SS, RM - performed experiments and substantially contributed to the acquisition of data and its analysis. All authors were involved in the interpretation of data. JL and JM drafted the manuscript. All authors revised the manuscript critically for important intellectual content.

## Abstract

**Objective:** Translocation of gut bacterial lipopolysaccharide (LPS) is associated with complications of cirrhosis. Current strategies to target bacterial translocation are limited to antibiotics with risk of resistance. This study aims to explore therapeutic potential of a non-absorbable, engineered carbon bead, Yaq-001 in cirrhosis and acute-on-chronic liver failure (ACLF) models.

**Design:** The performance of Yaq-001 was evaluated in *in vitro* studies. Two-rodent models of cirrhosis (4-week, bile duct ligation (BDL): Sham (n=36); Sham-Yaq-001 (n=30); BDL (n=37); BDL-Yaq-001 (n=44)) and ACLF (BDL-LPS: Sham-LPS (n=9); Sham-LPS-Yaq-001 (n=10); BDL-LPS (n=16); BDL-LPS-Yaq-001(n=12)). The treated-groups received Yaq-001 for 2-weeks. Samples were collected for assessment of organ and immune function, transcriptomics, microbiome composition and metabolomics.

**Results:** *In vitro*, Yaq-001 exhibited rapid adsorption kinetics for endotoxin and bile acids without exerting an antibiotic effect. *In vivo*, Yaq-001 produced significant improvement in ALT, ammonia, liver cell death, portal pressure, markers of systemic inflammation and renal function in BDL animals. Yaq-001-treated ACLF animals had significantly better survival, ALT, portal pressure, brain water and creatinine. *Ex-vivo* LPS-induced reactive oxygen species production in portal venous monocytes and Kupffer cell populations was diminished with Yaq-001 treatment. Transcriptome analysis demonstrated a significant modulation of inflammation, cell death and senescence pathways in the liver, kidneys, brain and colon of Yaq-001-treated BDL rats. Yaq-001 impacted positively on the microbiome composition with significant modulation of *Family Porphyromonadaceae* and *Genus Barnesiella*. Urinary ^1^HNMR analysis suggested a shift in metabolomic signature in Yaq-001-treated BDL rats.

**Conclusions:** This study provides strong pre-clinical rationale for developing Yaq-001 for treatment of patients with cirrhosis.

**Significance of this study:** *What is already known on this topic?:* Current strategies to target bacterial translocation in cirrhosis are limited to antibiotics with risk of resistance. Yaq-001 is an insoluble, non-absorbable, non-antibiotic, engineered carbon bead of tailored porosities, which works as an adsorbent in the gut and is completely excreted after oral administration.

*What this study adds?:*

1. Yaq-001 rapidly adsorbs endotoxin, ammonia and bile acids without influencing bacterial growth kinetics *in vitro*.
2. Yaq-001 reduces mortality of ACLF animals and impacts positively on markers of gut permeability, liver injury, portal pressure, brain and kidneys in rodent models of cirrhosis and ACLF.
3. Yaq-001 administration was associated with positive impact on the composition of the gut microbiota, reduction in severity of endotoxemia and ammonia, which significantly reduced the severity of inflammation, cell death, signaling pathways and LPS sensitivity.

*How this study might affect research, practice or policy?:* The data provide the pre-clinical rationale to proceed to clinical trials in patients with cirrhosis aiming to prevent the occurrence of complications.

## INTRODUCTION

Gut-derived bacterial ligands, in particular endotoxin, drive a dysregulated inflammatory response, which has been implicated in the complications of cirrhosis such as sepsis, spontaneous bacterial peritonitis, renal dysfunction and hepatic encephalopathy^1–3^. This dysregulated inflammatory response is central in the development of acute-on-chronic liver failure (ACLF)^4^. Markers of bacterial translocation such as endotoxin and bacterial DNA have been shown to be associated with complications of cirrhosis and diminished survival highlighting their pathogenic importance^5–7^. The microbiome in cirrhosis is characterized by reduced diversity and abundance of autochthonous bacteria^1^. Whilst antibiotics have been shown to impact positively on complications of cirrhosis, their use is associated with bacterial superinfection and antibiotic resistance^8, 9^. Furthermore, antibiotics reduce bacterial diversity further rendering the microbiome less resilient.

One of the consequences of bacterial translocation in cirrhosis is that the endotoxin-sensing pathways in different organs are known to be primed resulting in heightened susceptibility to organ injury^3, 10^. Adsorption of free endotoxin without exerting direct effects on bacterial growth kinetics, therefore has the potential to attenuate susceptibility to organ injury without producing the deleterious effects on the microbiome. Considering this, we developed a synthetic non-absorbable, non-antibiotic, endotoxin sequestrant and generated the hypothesis that this may be a novel therapeutic strategy to restore the microbiome, prevent bacterial translocation, systemic inflammation and cirrhosis complications. Yaq-001 is a non-absorbable, engineered, activated carbon of bimodal porosity tailored to the micro (<2nm) and meso-macroporous (30-200 nm) range and high surface area^11–13^. These properties confer a high adsorptive capacity for larger biologically relevant molecules such as bacterial toxins in addition to smaller intraluminal targets. The most closely associated experimental oral adsorbent is AST-120, a microporous carbon bead, which has not been shown to be efficacious in patients with hepatic encephalopathy^14^.

In this study, we sought to determine the adsorptive capacity of Yaq-001 and its effect on bacterial growth kinetics in *in vitro* studies. We then evaluated the *in vivo* biological effects of Yaq-001 in two animal models representing characteristics of cirrhosis and ACLF respectively. We studied the effects of Yaq-001 on measures of multiorgan function, systemic and portal hemodynamics, immune function, multiorgan transcriptomics and microbiome composition.

## METHODS

## STUDIES *IN VITRO*

Methodological details are described in **Supplementary section**. Adsorption of biomolecules of varying molecular weights (albumin, myoglobin, and caffeine) was evaluated. Then, the effect of Yaq-001 on the kinetics of bacterial growth was studied for *Staphylococcus aureus (S. aureus)* and *Escherichia coli (E. coli)*. Scanning electron microscopy was performed to characterise the beads and pore size distribution was assessed using mercury porosimetry.

## STUDIES *IN VIVO*

The methodological details are described in **Supplementary section**.

### Study design

This study aimed to characterize the therapeutic potential of Yaq-001 in models of cirrhosis and ACLF. The different experiments performed are shown in **Fig.S1**. To evaluate the effect of Yaq-001 in cirrhosis and for prevention of ACLF, studies in two animal models were performed.

#### Cirrhosis

Sham (n=36); Sham-Yaq-001(n=30); BDL (n=37); BDL-Yaq-001 (n=44); Sham-LPS (n=9).

#### Prevention of ACLF

Sham-LPS-Yaq-001 (n=10); BDL-LPS (n=16); BDL-LPS-Yaq-001(n=12).

Yaq-001 (0.4 g/100 g body weight per day) was administered for 2-weeks prior to sacrifice. At the time of sacrifice, mean arterial pressure (MAP) and portal pressure were measured. Blood and tissues were then samples for later analysis.

### Analysis of biosamples

Plasma levels of alanine transaminase (ALT), alkaline phosphatase (ALP), total bilirubin (TBIL), albumin, bile acids, creatinine, urea, ammonia, endotoxin, bacterial DNA, D-lactate and brain water were measured.

Peripheral blood cells and Kupffer cell reactive oxidant species (ROS) were measured. Hematoxylin-Eosin (H&E), Picrosirius Red (PSR) staining and TUNEL stains were performed in liver tissues. The mRNA in different organs was analyzed by using nSolver4.0 software (NanoString Technologies). To define effect on the microbiome, 16s microbiome study was performed. To determine the effect of Yaq-001 on modulating metabolism, urinary ^1^H-NMR analysis was performed.

## STATISTICAL ANALYSIS

Based on the *in vitro* studies, we anticipated a 50% decrease in circulating endotoxin in the treatment groups with an alpha error of 0.05 and power of 80%, resulting in a minimum sample size of 5 animals/ group. As this study included several pathophysiological end points, multiple experimental groups were included over a 5-year period. All the data accrued from these studies are described in this paper. All 194 rats in eight groups from three independent batches were included in the analysis as shown in **Fig.S1** and **Table S1**.

Group comparisons for continuous variables were performed using Man-Whitney U test (no-normal distribution) or unpaired t-test (normal distribution) and for categorical variables by using Chi-squared test. The data were analysed using R package (R version 4.4.4). 16s microbiome study and circos correlation were analyzed by using Wilcoxon rank sum test and spearman correlation. Software used included Graphpad Prism 9.0 (GraphPad software, Inc., San Diego, CA).

## RESULTS

## STUDIES *IN VITRO*

### Yaq-001 rapidly adsorbs endotoxin without influencing bacterial growth kinetics

Yaq-001 beads exhibited a consistent macroporous structure with a bead diameter within the 250-500μm range and an internal porosity in the nanoporous range (**Fig.1A**). Mercury porosimetry showed that Yaq-001 had a consistent pore size distribution plot in the meso-macroporous range from 30-200 nm (**Fig.S2**). Yaq-001 rapidly adsorbed albumin (66.5kDa), myoglobin (16.7kDa) and caffeine (0.194kDa) representing different sized biomolecules (**Fig.1B**). Yaq-001 adsorbed LPS (18kDa) reducing the concentrations from 2.5 to 1.5 EU mL^-1^ (60%) within 30 minutes. No endotoxin was detected in the control solution (0 EU mL^-1^) (**Fig.1B**). Yaq-001 also adsorbed a range of bile acids (**Fig.1C**). Direct co-incubation of Yaq-001 with bacterial suspensions of either *E. coli* or *S. aureus* indicated that Yaq-001 did not affect bacterial growth kinetics for either species following direct contact in comparison to the antibiotic controls (**Fig.1D**).

**Fig. 1.**
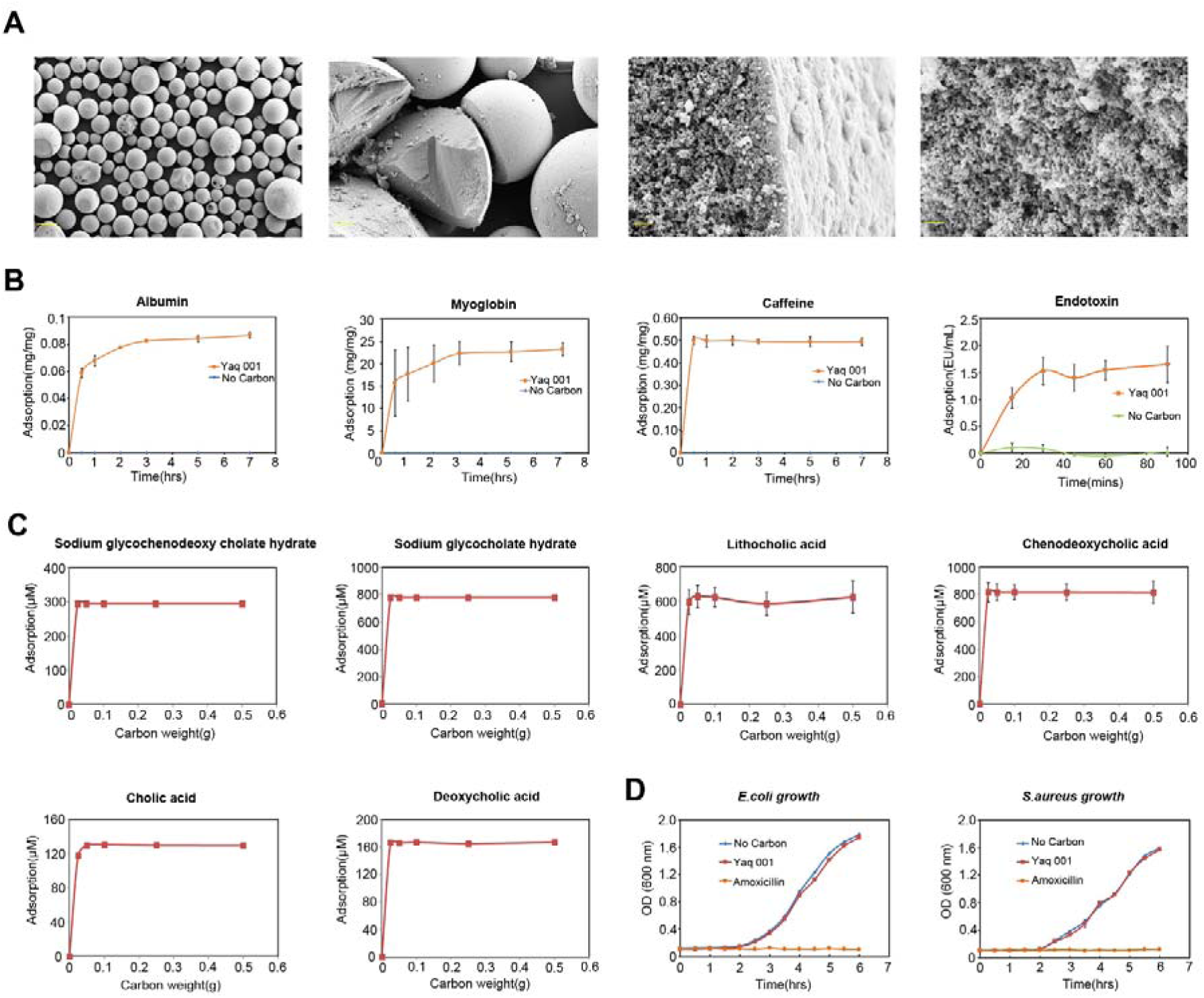
Physical characteristics of Yaq-001, adsorptive capacity and effect on bacterial kinetics. (A) Physical characteristics of Yaq-001 using scanning electron microscopy demonstrating meso-macroporous domains (Scale bars: from left to right: 200 μm, 40 μm, 400 nm and 400 nm) (n=3). (B) Adsorption kinetics of albumin, myoglobin, caffeine and endotoxin by Yaq-001 (n=3). (C) Adsorption of a range of bile acids including sodium glycochenodeoxy cholate hydrate, sodium glycocholate hydrate, lithocholic acid, chenodeoxycholic acid, cholic acid and deoxycholic acid by Yaq-001(n=3). (D) Growth curves of *E. coli* (n=3) and *S. aureus* (n=3) in the presence of Yaq-001, amoxicillin and vehicle.

## STUDIES *IN VIVO*

### Studies in BDL rats

#### Effect of Yaq-001 on liver injury and portal pressure

BDL rat model was used to assess the effect of Yaq-001 in cirrhosis (**Fig.2A**). Significant reduction in 4-week body weight was observed in BDL rats (p<0.0001), which was prevented by administration of Yaq-001 (p=0.045) (**Fig.2B**). Yaq-001 was associated with a significantly lower plasma ALT (p=0.007) (**Fig.2C**). Alkaline phosphatase, bilirubin and albumin were not impacted by Yaq-001 (**Fig.S3A, B, C**). Total bile acid concentrations were not different between the BDL and Sham groups and there was no significant impact of Yaq-001 (**Fig.S3E**). MAP was lower in BDL animals and no effect of Yaq-001 was observed (**Fig.S3F**). Yaq-001 resulted in a significant reduction in portal pressure compared to untreated BDL rats [(median (IQR) 11.1 mm Hg (10.3-11.7) *vs* 12.4 mm Hg (10.8-13.3), (p=0.025)] (**Fig.2C**). TUNEL assay showed significantly more intense staining in the liver tissue of BDL compared to Sham rats (**Fig.2D**) (p<0.0001), which was significantly reduced in Yaq-001-treated BDL rats compared to untreated-BDL rats (p=0.025). Collagen proportionate area (CPA) was significantly higher values in BDL rats (p=0.0007), which was unchanged with Yaq-001 (p=0.122) (**Fig.S3D**).

**Fig. 2.**
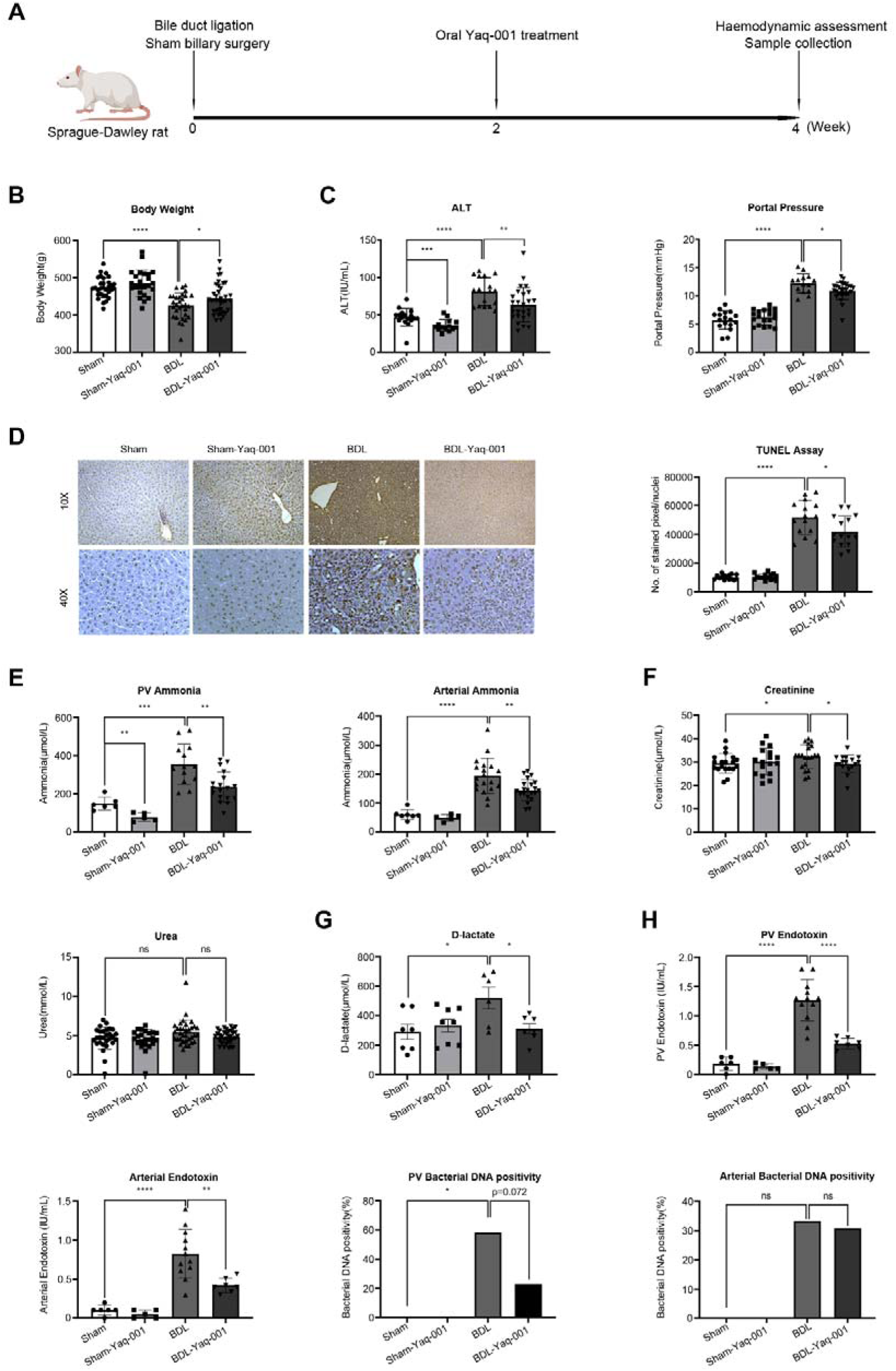
Effect of Yaq-001 on organ dysfunction, endotoxemia and bacterial translocation in BDL rats. (A) Rats underwent bile duct ligation for 4-weeks as a model of cirrhosis (n=23-37/group) in Experiment 1 and the treatment groups received Yaq-001 for 2 weeks before sacrifice. (B) 4-week body weight in four groups: Sham (n=31), Sham-Yaq-001 (n=24), BDL (n=31) and BDL-Yaq-001 (n=38) in Experiments 1 and 2. Significantly lower final body weights were observed in BDL compared to Sham controls (p<0.001). Yaq-001-treated BDL rats had a significantly higher body weights compared to untreated-BDL rats (p<0.05). (C) Plasma alanine transaminase (ALT) concentrations in Sham (n=17), Sham-Yaq-001 (n=14), BDL (n=17) and BDL-Yaq-001 (n=26) groups. Portal pressure (PP) measurements in Sham (n=17), Sham-Yaq-001 (n=19), BDL (n=14) and BDL-Yaq-001 (n=26) groups. Significantly higher ALT and PP were observed in BDL compared to Sham controls (p<0.0001). Yaq-001-treated BDL rats had a significantly lower ALT and PP compared to untreated-BDL rats (p<0.01, p<0.05). (D) TUNEL assay of liver tissue with quantification of staining by digital image analysis. Significantly higher TUNEL staining was observed in BDL compared to Sham controls (p<0.0001). Yaq-001-treated BDL rats had a significantly lower TUNEL staining compared to untreated-BDL rats (p<0.05) indicative of a reduction in liver cell death with Yaq-001 treatment. (E) Arterial ammonia concentrations in Sham (n=7), Sham-Yaq-001 (n=5), BDL (n=19), BDL-Yaq-001(n=21) groups. Portal venous ammonia concentrations in Sham (n=6), Sham-Yaq-001 (n=5), BDL (n=13), BDL-Yaq-001(n=18) groups. Significantly increased arterial ammonia concentrations and portal venous ammonia concentrations were observed in BDL compared to Sham controls (p<0.0001, p=0.0001). Yaq-001 significantly decreased arterial and portal venous ammonia concentrations in BDL rats (p<0.01 for both). (F) Serum creatinine in Sham (n=19), Sham-Yaq-001 (n=17), BDL (n=20), BDL-Yaq-001 (n=17) and urea in Sham (n=28), Sham-Yaq-001 (n=23), BDL (n=30), BDL-Yaq-001 (n=34) groups. Yaq-001 markedly decreased serum creatinine levels in BDL rats (p<0.05). (G) Plasma D-lactate in Sham (n=7), Sham-Yaq-001 (n=8), BDL (n=6), BDL-Yaq-001 (n=7). Plasma D-lactate was significantly increased in the BDL group compared with Sham animals (p<0.05). Yaq-001 resulted in a significant reduction in plasma D-lactate in BDL rats (p<0.05). (H) Portal venous [Sham (n=6), Sham-Yaq-001 (n=5), BDL (n=12) and BDL-Yaq-001 (n=7)] and arterial endotoxin concentrations [Sham (n=6), Sham-Yaq-001 (n=5), BDL (n=12) and BDL-Yaq-001 (n=7)]. Portal venous [Sham (n=6), Sham-Yaq-001 (n=5), BDL (n=12) and BDL-Yaq-001 (n=13)] and arterial plasma bacterial DNA positivity [Sham (n=6), Sham-Yaq-001 (n=6), BDL (n=12) and BDL-Yaq-001 (n=7)]. Significantly higher portal venous endotoxin and arterial endotoxin were observed in BDL rats compared to Sham rats (p<0.0001). Significantly higher portal venous plasma bacterial DNA positivity was observed in BDL rats compared to Sham rats (p<0.05). Yaq-001 administration was associated with a significant reduction of portal venous and arterial endotoxin compared to untreated-BDL rats (p<0.0001, p<0.01). Yaq-001 administration reduced bacterial DNA positivity, which was not statistically different (p>0.05).

#### Effect of Yaq-001 on ammonia, organ dysfunction, endotoxemia and bacterial translocation

##### Ammonia

Arterial and portal venous ammonia concentrations were significantly increased in BDL rats (p<0.0001), which was significantly reduced by Yaq-001 [(p=0.003) and (p=0.004) respectively] (**Fig.2E**). None of the animals showed signs of hepatic encephalopathy.

##### Kidneys

BDL animals had significantly higher plasma creatinine (p=0.049), which was significantly reduced with Yaq-001 (p=0.025) (**Fig.2F**). Urea was higher in BDL group (p=0.092), which was reduced with Yaq-001 treatment (p=0.095) (**Fig.2F**).

##### Gut permeability, Endotoxemia, Bacterial DNA and Cytokines

The microbial metabolite, D-lactate, a marker of gut-specific intestinal barrier damage and translocation^15^ was significantly increased in BDL rats (p=0.032) and was significantly reduced by Yaq-001 (p=0.035) (**Fig.2G**). BDL rats exhibited marked endotoxemia in the portal vein and the artery (p<0.0001 for each), which was significantly reduced with Yaq-001 [(p<0.0001) (p=0.003) respectively] (**Fig.2H**). Portal venous bacterial DNA was detectable in significantly higher number of BDL rats (p<0.05), which was markedly reduced in Yaq-001 administered BDL rats (p=0.08) (**Fig.2H**). Plasma IL-β concentration were higher in the BDL rats but no significant differences were observed in TNF-a, IL-6 and IL-10. No significant changes were seen with Yaq-001 (**Table S2**).

### Studies in the ACLF model

This experiment was performed to determine whether Yaq-001 treatment for 2-weeks prevents the occurrence of ACLF when BDL animals are administered LPS (**Fig.S1**, **Fig.3A**).

**Fig. 3.**
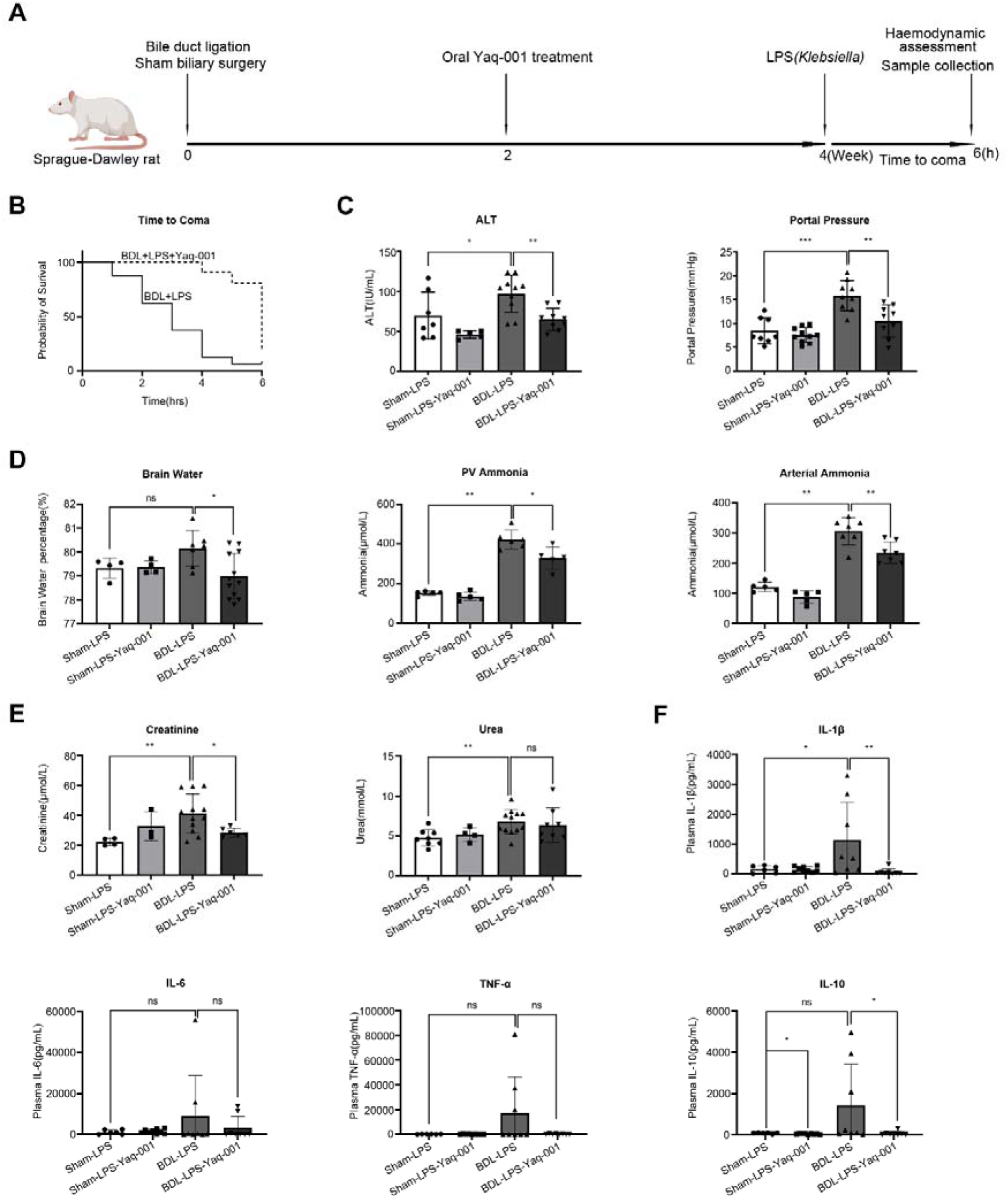
Effect of Yaq-001 on multiorgan function in ACLF. (A) Rats underwent sham biliary surgery or bile duct ligation (BDL) for 4-weeks. The treated group received Yaq-001 for two weeks prior to LPS injection. Animals were sacrificed either at coma stages or 6 hours after LPS injection (n=9-16/group). (B) Kaplan-Meier analysis of BDL-LPS rats with (n=16) or without (n=12) Yaq-001 treatment. Yaq-001 treatment significantly improved the survival of BDL-LPS rats compared to untreated-BDL-LPS rats (log rank test, p=0.003). (C) Plasma ALT concentrations in Sham-LPS (n=7), Sham-LPS-Yaq-001 (n=5), BDL-LPS (n=10) and BDL-LPS-Yaq-001 (n=9) groups. PP measurements in Sham-LPS (n=8), Sham-LPS-Yaq-001 (n=10), BDL-LPS (n=9) and BDL-LPS-Yaq-001 (n=9) groups. Yaq-001-treated BDL-LPS rats had a significantly lower ALT and PP compared to untreated-BDL-LPS rats (p<0.005). (D) Brain water percentage in Sham-LPS (n=4), Sham-LPS-Yaq-001 (n=4), BDL-LPS (n=7), BDL-LPS-Yaq-001 (n=13) groups. Arterial ammonia concentrations in Sham-LPS (n=5), Sham-LPS-Yaq-001 (n=5), BDL-LPS (n=7), BDL-LPS-Yaq-001 (n=7) groups. Portal venous ammonia concentrations in Sham-LPS (n=5), Sham-LPS-Yaq-001 (n=5), BDL-LPS (n=6), BDL-LPS-Yaq-001 (n=5) groups. Yaq-001 decreased brain water percentage and arterial/portal venous ammonia concentrations in BDL-LPS rats compared to untreated rats (p<0.05, p<0.01, p<0.05). (E) Serum creatinine in Sham-LPS (n=4), Sham-LPS-Yaq-001 (n=3), BDL-LPS (n=12) and BDL-LPS-Yaq-001 (n=6) groups. Serum urea in Sham-LPS (n=8), Sham-LPS-Yaq-001 (n=4), BDL-LPS (n=12) and BDL-LPS-Yaq-001 (n=8) groups. Yaq-001 significantly decreased creatinine levels in BDL-LPS rats (p<0.05). (F) Plasma cytokines in Sham-LPS (n=6), Sham-LPS-Yaq-001 (n=9), BDL-LPS (n=8) and BDL-LPS-Yaq-001 (n=8) groups. Yaq-001 significantly decreased plasma IL-1β and IL-10 concentrations in BDL-LPS groups (p<0.01, p<0.05).

#### Survival

Animals were sacrificed either at coma stages (considered as a surrogate for mortality) or at 6-hours post LPS. Yaq-001 significantly reduced time to coma of BDL-LPS rats compared to untreated controls (p<0.01) (**Fig.3B**). All animals in the two Sham groups were alive at 6-hours following LPS (data not shown).

*Liver*: Yaq-001 was associated with significantly lower ALT in BDL-LPS rats compared to untreated rats (p=0.004) (**Fig.3C**). No significant effect of Yaq-001 was observed on alkaline phosphatase, bilirubin and albumin (**Fig.S4 A, B, C**). The severity of fibrosis measured using CPA and the body weight were unchanged (**Fig.S4D, E**).

#### Systemic and Portal hemodynamics

No significant difference in MAP was observed with Yaq-001 (**Fig.S4F**) but Yaq-001 produced a significant reduction in portal pressure compared to untreated controls (p=0.003), (**Fig.3C**).

#### Brain

Yaq-001 significantly reduced brain water compared with untreated-rats (p=0.017) (**Fig.3D**). Arterial and portal venous ammonia concentrations were significantly increased in BDL-LPS rats, which was significantly reduced in Yaq-001-treated animals [(p=0.007) and (p=0.017) respectively] (**Fig.3D**).

#### Kidneys

Creatinine concentrations were significantly higher in BDL-LPS animals (p=0.004), which was significantly reduced by Yaq-001 (p=0.03) (**Fig.3E**).

#### Cytokines

BDL-LPS group had a significantly higher plasma IL-1β, which was significantly reduced with Yaq-001 (p=0.003) (**Fig.3F**). Plasma IL-10 was higher in BDL-LPS and was significantly reduced with Yaq-001 (p=0.028) (**Fig.3F**). No significant differences were observed in IL-6 or TNF-α concentrations between any of the groups (**Table S2**).

### Effect of Yaq-001 on peripheral blood cells and Kupffer cells

Significant increase in total leucocyte, neutrophil and monocyte counts in the artery and portal vein were observed with BDL rats (**Fig.4A**, B) (p=0.008 and p=0.016 respectively), which was significantly reduced with Yaq-001 in the arterial blood and insignificantly reduced in the portal vein (**Fig. 4B**). To determine whether Yaq-001 impacts on the response of peripheral inflammatory cells and Kupffer cells to generate reactive oxygen species (ROS) to LPS *ex vivo,* studies using isolated cells incubated with LPS, were performed. Yaq-001 was associated with significantly lower LPS-induced ROS production in CD163^-^ Kupffer cells in BDL rats (p=0.036) and portal venous CD43^hi^ monocyte populations of BDL rats (p=0.029) (**Fig.4C**).

**Fig. 4.**
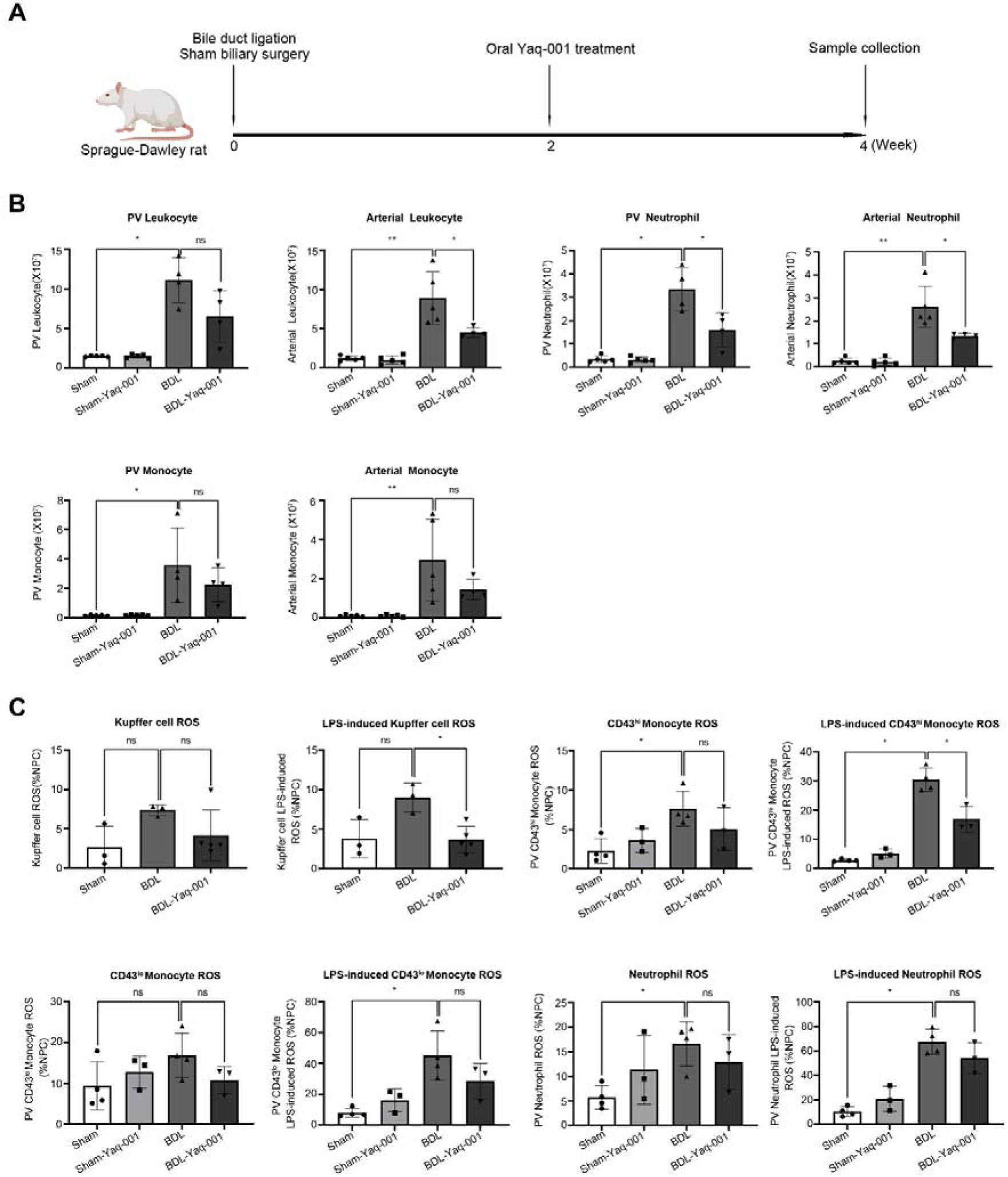
Effect of Yaq-001 on peripheral and Kupffer cell populations. (A) Rats underwent bile duct ligation for 4-weeks as a model of cirrhosis (n=4-5/group). Yaq-001 was administered to treated rats 2-weeks before sacrifice. Blood was sampled from the artery or the portal vein and the liver was perfused to isolate Kupffer cells. (B) Absolute portal venous total leukocyte, neutrophil and monocyte populations in Sham (n=5), Sham-Yaq-001 (n=5), BDL (n=4) and BDL-Yaq-001 (n=4) groups. Absolute arterial total leukocyte, neutrophil and monocyte populations in Sham (n=5), Sham-Yaq-001 (n=5), BDL (n=5) and BDL-Yaq-001 (n=4) groups. BDL was associated with a significant increase in leukocyte, neutrophil and monocyte levels in portal vein and artery compared to Sham controls (p<0.05, p<0.01) respectively. Yaq-001 significantly decreased total leukocyte level in artery and neutrophil levels in both portal vein and artery (p<0.05). (C) Constitutive and LPS-induced ROS production in CD163-gated liver non-parenchymal cell fraction (n=3-5/group), portal venous monocytes (n=3-4/group) and portal venous neutrophil populations (n=3-4/group) (expressed as a percentage of the parent population) in Sham, Sham-Yaq-001, BDL and BDL-Yaq-001. BDL resulted in a significant increase in LPS-induced monocyte and neutrophil ROS production (p<0.05). Yaq-001 significantly attenuated LPS-induced ROS production both in portal venous monocytes and Kupffer cell populations (p<0.05).

### Transcriptomic analysis of gene expression profiles from the Liver, Colon, Brain and Kidneys

Multiorgan transcriptomic analysis was performed to determine the possible molecular mechanisms underlying the clinical effects of Yaq-001. The four groups studied were as follows: Sham (n=3), Sham-Yaq-001 (n=3), BDL (n=3) and BDL-Yaq-001 (n=4) (**Fig.5A**, **Fig.6A**). All differentially expressed genes (DEGs) and related pathways in the liver, colon, kidney and brain are listed in **Table S3**. The top 20 and significant DEGs are listed in **Table S4**.

**Fig. 5.**
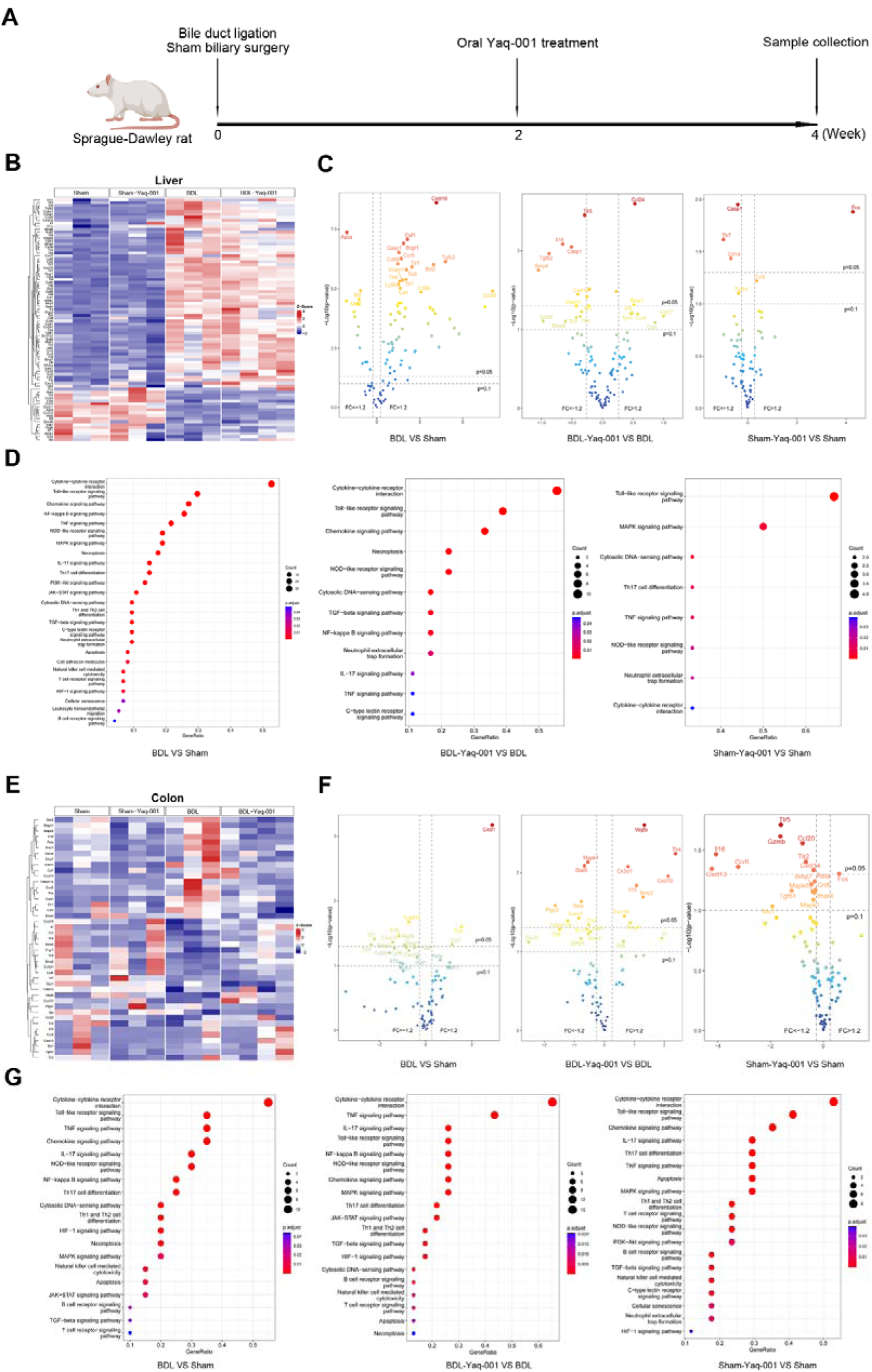
Effect of Yaq-001 on gene expression profiles in the liver and gut in BDL rats. (A) Rats underwent bile duct ligation for 4-weeks as a model of cirrhosis (n=3-4/group) and the treatment groups received Yaq-001 for 2-weeks before sacrifice. Liver and colon were collected for transcriptomic analysis. (B) Heatmap of differentially expressed genes (DEGs) in liver tissue between Sham (n=3), Sham-Yaq-001 (n=3), BDL (n=3) and BDL-Yaq-001 (n=4) groups. DEGs were identified at 1.2-fold change and p=0.1 threshold in three pairwise groups (BDL versus Sham, BDL-Yaq-001 versus BDL, Sham-Yaq-001 versus Sham). (C) Volcano plot of pairwise DEGs in liver among Sham (n=3), Sham-Yaq-001 (n=3), BDL (n=3) and BDL-Yaq-001 (n=4) groups. The vertical dashed lines indicated the threshold for 1.2-fold change. The horizontal dashed line indicated the adjusted p=0.05 and p=0.1 threshold. The right part indicates up-regulation of gene expression, and the left part indicates down-regulation of gene expression. The top 20 genes are indicated by gene names. (D) Functional enrichment analysis of liver genes pairwise based on the Kyoto Encyclopedia of Genes and Genomes (KEGG) database. The significantly changed pathways are shown in panels including inflammation, TLR signaling, cell death, cell senescence and intracellular signaling. (E) Heatmap of DEGs in colonic tissue between Sham (n=3), Sham-Yaq-001(n=3), BDL (n=3) and BDL-Yaq-001 (n=4) groups. DEGs were identified at 1.2-fold change and p=0.1 threshold in three pairwise groups (BDL versus Sham, BDL-Yaq-001 versus BDL, Sham-Yaq-001 versus Sham). (F) Volcano plot of pairwise DEGs in colon among Sham (n=3), Sham-Yaq-001 (n=3), BDL (n=3) and BDL-Yaq-001 (n=4) groups. The vertical dashed lines indicated the threshold for 1.2-fold change. The horizontal dashed line indicates adjusted p=0.05 and p=0.1 threshold. The right part indicates up-regulation of gene expression, and the left part indicates down-regulation of gene expression. The top 20 genes are indicated by gene names. (G) Functional enrichment analysis of colon genes in pairwise three groups based on KEGG database. The significantly changed pathways are shown in panels including inflammation, TLR signaling, cell death, cell senescence and intracellular signaling.

**Fig. 6.**
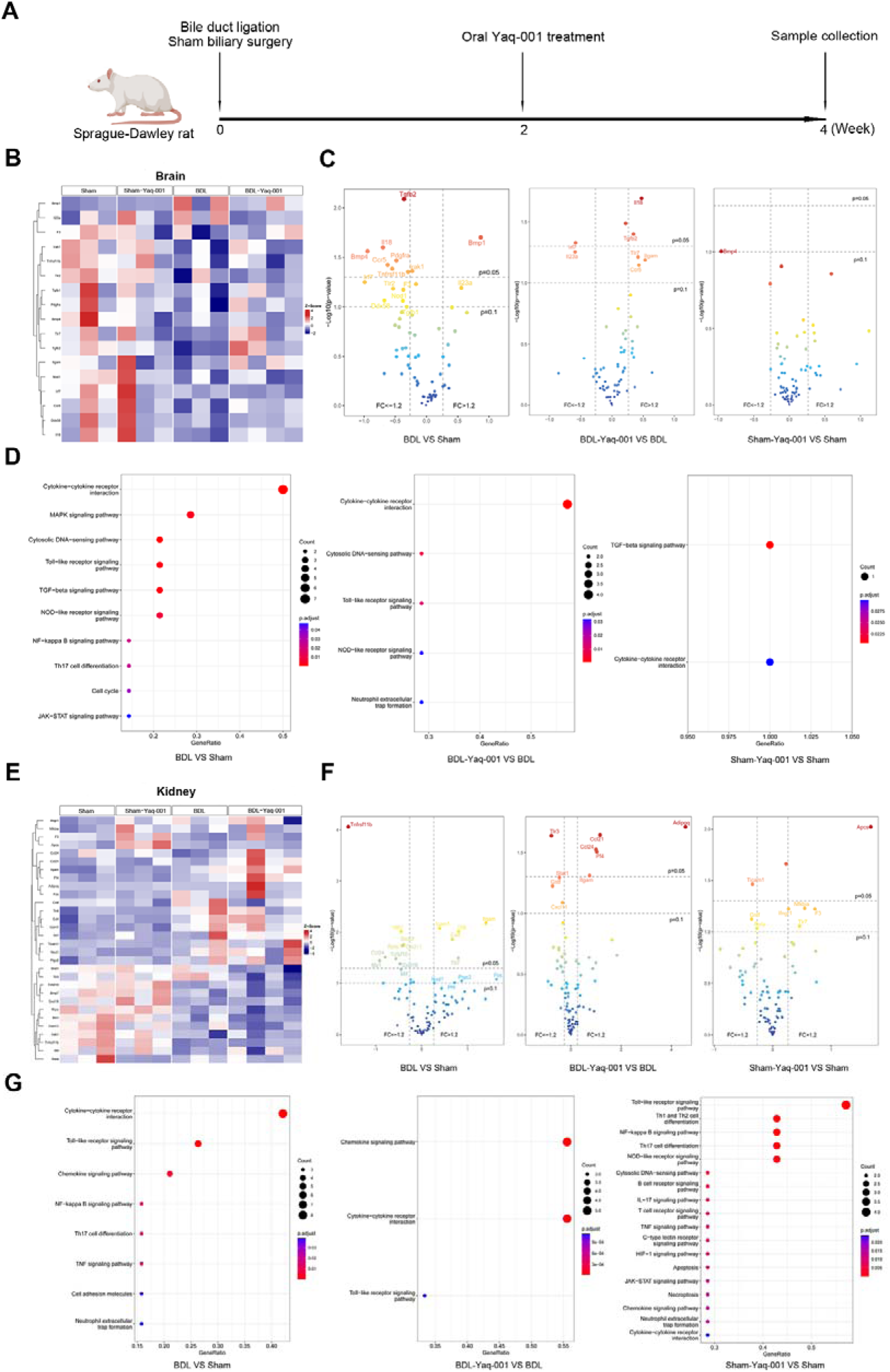
Effect of Yaq-001 on gene expression profiles in the brain and kidneys in BDL rats. (A) Rats underwent bile duct ligation for 4-weeks as a model of cirrhosis (n=3-4/group) and the treatment groups received Yaq-001 for 2-weeks before sacrifice. Brain and kidneys were collected for transcriptomic analysis. (B) Heatmap of DEGs in brain tissue between Sham (n=3), Sham-Yaq-001 (n=3), BDL (n=3) and BDL-Yaq-001 (n=4) groups. DEGs were identified at 1.2-fold abundance difference and p=0.1 threshold in three pairwise groups (BDL versus Sham, BDL-Yaq-001 versus BDL, Sham-Yaq-001 versus Sham). (C) Volcano plot demonstrates pairwise DEGs in the brain among Sham (n=3), Sham-Yaq-001 (n=3), BDL (n=3) and BDL-Yaq-001 (n=4) groups. The vertical dashed lines indicates the threshold for 1.2-fold abundance difference. The horizontal dashed line indicates adjusted p=0.05 and p=0.1 threshold. The right part indicates up-regulation of gene expression, and the left part indicates down-regulation of gene expression. The top 20 genes are indicated by the gene names. (D) Functional enrichment analysis of brain is shown pairwise for three groups based on the KEGG database. The significantly changed pathways were shown in panels including inflammation, TLR signaling, cell senescence and intracellular signaling. (E) Heatmap of DEGs in kidney tissue between Sham (n=3), Sham-Yaq-001(n=3), BDL (n=3) and BDL-Yaq-001 (n=4) groups. DEGs were identified at 1.2-fold abundance difference and p=0.1 threshold in three pairwise groups (BDL versus Sham, BDL-Yaq-001 versus BDL, Sham-Yaq-001 versus Sham). (F) Volcano plot demonstrated pairwise DEGs in the kidney among Sham (n=3), Sham-Yaq-001 (n=3), BDL (n=3) and BDL-Yaq-001 (n=4) groups. The vertical dashed lines indicates the threshold for 1.2-fold abundance difference. The horizontal dashed line indicates adjusted p=0.05 and p=0.1 threshold. The right part indicates up-regulation of gene expression, and the left part indicates down-regulation of gene expression. The Top 20 genes are indicated by the gene names. (G) Functional enrichment analysis of kidney shown pairwise for three groups based on the KEGG database. The significantly changed pathways are shown in panels including inflammation and TLR signaling.

#### Effect of Yaq-001 on gene expression profiles in the liver and gut in BDL rats

*Liver*: Analysis of liver tissue showed 82 DEGs at the threshold of 1.2-fold change and p=0.1 in the four groups (**Fig.5B**). Compared with the Sham group, expression of 62-genes was upregulated, and 15-genes were downregulated in BDL. These significantly changed genes were associated with inflammation, cell death and senescence. Compared to the untreated BDL group, the expression of 7-genes was upregulated and 12-genes were downregulated in the Yaq-001-treated BDL group, indicating the potential role of Yaq-001 in reducing inflammation, cell death and cell senescence. Furthermore, 2-genes were upregulated, and 4-genes downregulated in Sham-Yaq-001 group in comparison to Sham group (**Fig.5C**). Functional analysis demonstrated that BDL rats had enriched pathways related to inflammation, cell senescence, cell death, TLR signaling and other related signaling pathways in comparison with Sham (**Fig.5D**). Yaq-001 treatment targeted the altered pathways compared with untreated BDL group. Additionally, Yaq-001 treatment also changed the pathways in the liver when compared to Sham group, demonstrating its effect in rats even without cirrhosis (**Fig.5D**).

##### Colon

43 DEGs were identified from the colonic tissue (**Fig.5E**). 5-genes that correlated with inflammation and cell death were upregulated and 15-genes were downregulated in BDL compared with the Sham group. Moreover, the expression of 10-genes was upregulated, and 13-genes were downregulated with Yaq-001 treatment. Only 1-gene was upregulated in the Sham-Yaq-001 group, and 16-genes were downregulated with Yaq-001 compared with the untreated Sham group (**Fig.5F**). Functional analysis indicated that inflammation, cell senescence, cell death, TLR signaling and intracellular signaling were associated with BDL in comparison with the Sham group (**Fig.5G**). Yaq-001 targeted the altered pathways, indicating the potential mechanisms in the prevention of gut dysfunction and permeability (**Fig.5G**).

#### Effect of Yaq-001 on gene expression profiles in the brain and kidney in BDL rats

##### Brain

17 DEGs were identified from the brain tissue (**Fig.6B**). Compared with Sham group, expression of 2-genes was upregulated and 13-genes were downregulated in BDL animals. These significantly changed genes were associated with inflammation, cell death, and cell senescence. Compared to the untreated-BDL group, the expression of 5-genes was upregulated and 2-genes were downregulated in the Yaq-001-treated BDL group (**Fig.6C**). Functional analysis demonstrated that BDL rats had enriched pathways related to inflammation, cell senescence, cell death, TLR signaling and intracellular signaling (**Fig.6D**). Yaq-001 targeted cytokine-cytokine receptor interaction, cytosolic DNA-sensing pathway, TLR signaling pathway, NOD-like receptor signaling pathway, neutrophil extracellular trap formation, TGF-beta signaling pathway and cytokine-cytokine receptor interaction pathways compared to untreated-BDL group (**Fig.6D**).

##### Kidneys

30 DEGs were identified from kidney tissue (**Fig.6E**). 9-genes that correlated with inflammation were downregulated in BDL. The expression of 5-genes was upregulated and 4-genes were downregulated with Yaq-001 treatment compared to untreated-BDL group. 5-genes were upregulated in Sham-Yaq-001 group, and 3-genes were downregulated with Yaq-001 compared with untreated-Sham group (**Fig.6F**). Functional analysis indicated that inflammation and TLR signaling were associated with BDL in comparison with Sham (**Fig.6G**). Compared with the untreated-BDL group, Yaq-001 targeted the altered pathways, indicating the potential mechanisms in the prevention of renal dysfunction (**Fig.6G**).

#### Effect of Yaq-001 on the gut microbiome profile

The effects of Yaq-001 on the microbiome bacterial composition was assessed by metataxonomics. At the family level, an abundance of six bacteria were significantly changed at the threshold of 2-fold change and *Porphyomonadaceae* was significantly changed (p<0.05) comparing BDL with Sham **(Fig.7A)**. At genus level, 19 bacteria including were significantly changed in abundance. *Barnesiella* was significantly changed (p<0.05) comparing BDL with Sham group **(Fig.7B)**. These changes were reversed with Yaq-001 treatment compared to untreated-BDL rats (**Fig.S5A, B**, **Table S5 and Fig.S5C, D).** For between groups sample diversity, PERMANOVA analysis revealed a significant difference in beta diversity between groups (R2 = 0.32, p = 0.001). Yaq-001 appeared to moderately restore the beta diversity in the BDL group especially in PCoA2 axis (**Fig.S5E, F)**.

**Fig. 7.**
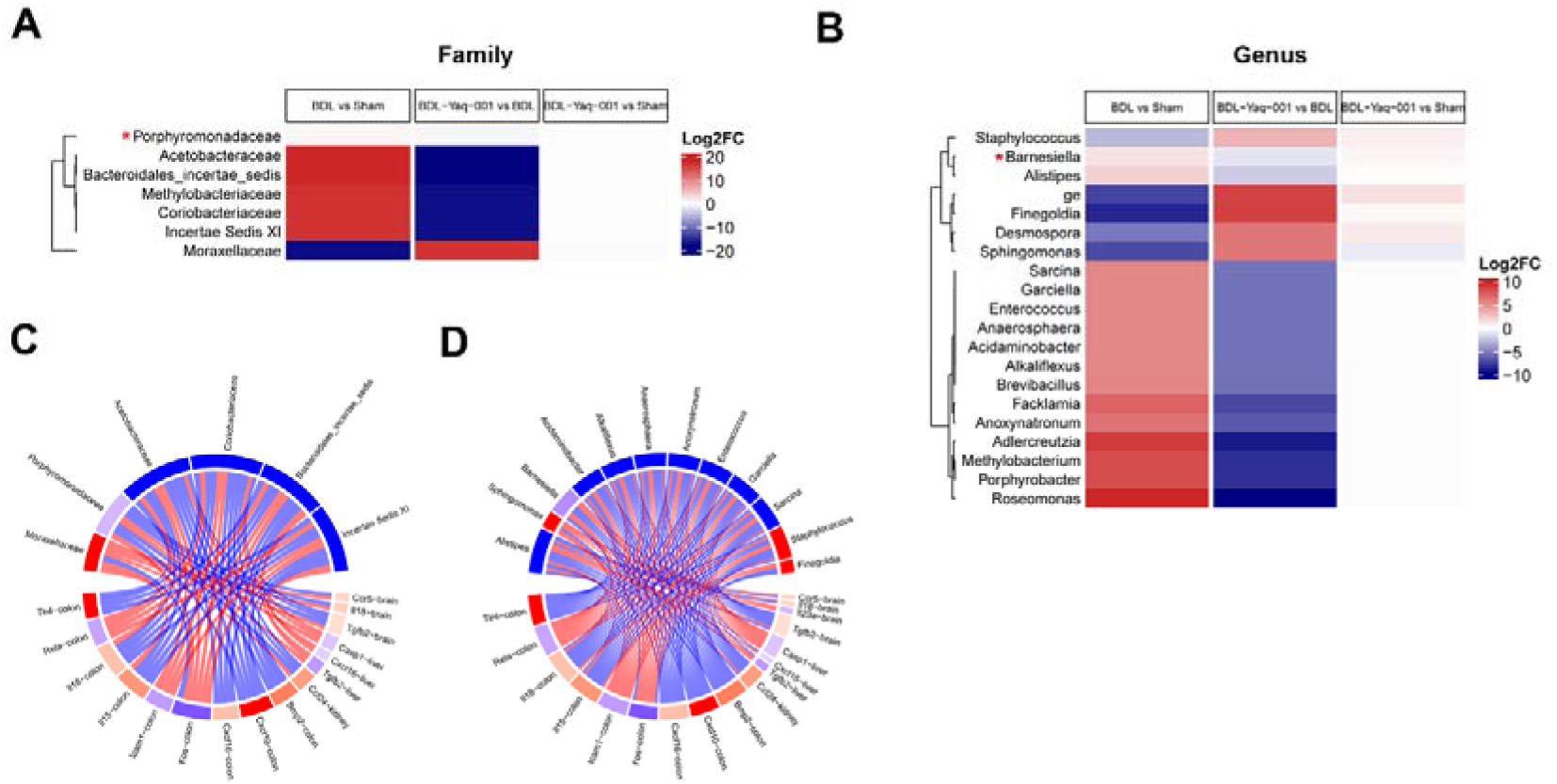
Effect of Yaq-001 treatment on the microbiome composition. (A) Heatmap of gut microbiome associated with the effect of Yaq-001 as determined by 16S PCR at the family level. The Family *Porphyromonadaveae* with asterisk was statistically differently abundant between BDL (n=7) vs Sham (n=6), and between BDL-Yaq-001 (n=7) vs BDL groups (n=7) (Wilcoxon rank sum test, p<0.05). The abundance of this family was statistically higher in BDL group than in Sham group, and its abundance statistically decreased in the BDL-Yaq-001 group than in the BDL group. The other six families in the heatmap were with marked fold changes between BDL vs Sham, and between BDL-Yaq-001 vs BDL groups (|log2FC|>2). Of these, five were more abundant in the BDL group than in the Sham group. The abundance largely decreased in the Yaq-001-treated group. In addition, of these, one family was less abundant in the BDL group than in the Sham group. The abundance increased in the Yaq-001-treated group. (B) Heatmap of gut microbiome at the Genus level. The Genus *Barnesiella* with asterisk was statistically differently abundant between BDL vs Sham, and between BDL-Yaq-001 vs BDL groups (Wilcoxon rank sum test, p<0.05). The abundance of this genus was statistically higher in BDL group than in the Sham group, and its abundance statistically decreased in the BDL-Yaq-001 group. The other 19 genera in the heatmap represent those with significant fold change values between BDL vs Sham, and between BDL-Yaq-001 vs BDL groups (|log2FC|>2). Of these, 14 were more abundant in the BDL group compared with the Sham group. The abundance decreased in the Yaq-001-reated BDL group. In addition, 5 genera were less abundant in the BDL group than in the Sham group. Their abundance increased in the Yaq-001-treated BDL animals. (C, D) Correlation plots between markedly changed genes and gut microbiome at family/genus. The genes were from amongst the top 20 changed genes in BDL animals with Yaq-001 treatment. Nodes represent either genes (lower semi-circular part) or bacteria (upper semi-circular part) at the family and genus level. The nodes are colored based on the log-fold change for the differential gene expression and differences in the bacterial abundance. The red nodes indicate an increase and blue nodes indicate a decrease. Edges represent the correlation coefficient calculated between genes and microbial genus or family with red indicating a positive correlation and blue a negative correlation. Correlation coefficients greater or equal to 0.4 were plotted in plot C (Spearman’s coefficient >= 0.4), and D shows all correlations.

To further investigate the potential importance of the changes in the microbiome induced by Yaq-001, we correlated these with all significantly changed DEGs and the top 20 DEGs in the four organs. Circos plots indicated a significant correlation between them (**Fig.7C, D and Fig.S6A, B**, **Fig.S6C)**. *Porphyromonadaceae*, was observed to positively correlate with three DEGs -TGFB2 and CASP1 in liver tissue, and FOS in colonic tissue. Also, it correlated negatively with five DEGs-TGFB2, IL-18 and CCR5 in brain tissue, CXCL10 in colon tissue and CCL24 in kidney tissue.

### Effect of Yaq-001 on metabolomic profile

Significant difference of acetate/creatinine, glycine/creatinine, lactate/creatinine, betaine/breatinine, trimethylamine oxide/creatinine and bile acid/creatinine ratio were observed in BDL compared to Sham. Treatment of BDL rats with Yaq-001 resulted in significant resolution of acetate/creatinine, glycine/creatinine and lactate/creatinine compared to the untreated BDL animals (**Fig.S7**).

## Discussion

This study explored the role of Yaq-001 in models of cirrhosis and ACLF. The study showed that Yaq-001 reduced the mortality of ACLF animals and impacted positively on markers of gut permeability, liver injury, portal pressure, brain and kidneys in two BDL models. These pleiotropic effects of Yaq-001 were associated with restoration of the composition of the microbiome bacterial community, reduction in the severity of endotoxemia and ammonia, severity of inflammation, cell death, signaling pathways and LPS sensitivity. The data provide the experimental rationale to proceed to further evaluation in clinical trials.

Translocation of bacteria, its products and metabolites are critically important in the pathogenesis of complications of cirrhosis^16–18^. Bacterial LPS plays a key role in driving systemic inflammation and the resultant organ failure in cirrhosis^1, 19^. Indeed, selective gut decontamination using norfloxacin or rifaximin are the current standard of care for secondary prophylaxis of patients with spontaneous bacterial peritonitis and hepatic encephalopathy respectively^20–21^. However, the use of these antibiotic strategies induces the risk of antibiotic resistance.^22^ The data presented here provide an alternative gut-restricted, non-antibiotic strategy, Yaq-001, which has the potential to diminish translocation and improve organ injury. The *in vitro* studies demonstrate that Yaq-001 has the optimal pore size distribution to bind intraluminal factors such as free endotoxin without significant effect on bacterial growth kinetics.

Endotoxemia has also been implicated in immune dysfunction resulting in a dysregulated systemic inflammatory response syndrome, which is strongly associated with the progression of cirrhosis and ACLF ^23^. Yaq-001 reduced the severity of endotoxemia and bacterial DNA positivity, which was associated with attenuated systemic inflammation. Significant improvements in LPS-induced ROS production were observed in trafficking portal venous monocytes suggesting that Yaq-001 had attenuated the primed state of monocyte/macrophage populations within the gut-liver axis. This observed reduction in LPS-induced ROS production may be important in explaining the reduction in plasma IL-1β in LPS-treated BDL rats.

Gut microbiota are important in modulating intestinal health, permeability, bacterial translocation, systemic inflammation and complications of cirrhosis^24–26^. BDL was associated with marked changes in the abundance of microbiota, which were reversed by Yaq-001. In particular, the abundance of *Porphyromonadaceae* and *Barnesiella* were significantly elevated in BDL rats and significantly decreased with Yaq-001. This change is potentially important as *Porphyromonadaceae* is a pro-inflammatory bacterium that has been positively correlated with hepatic encephalopathy^27^ and, *Barnesiella* and *Porphyromonadaceae* have been associated with liver cancer ^28.29^. Urinary NMR analysis reflects the combined metabolic status of both host and microbiota. Yaq-001 was associated with a distinct shift of acetate, glycine and lactate in metabolomic profile in BDL rats. They are products generated by mixed acid fermentation (MAF) typically by bacteria such as Enterobacter. MAF is not the preferred metabolic pathway for facultative anaerobes and may be indicative that Enterobacter populations are under conditions of metabolic stress in Yaq-001 treated BDL animals. As these species are often pathogenic in cirrhosis, this may represent a beneficial change.

Plasma D-lactate, a marker of increased gut permeability was reduced by Yaq-001^30^. Elevated plasma D-lactate levels in cirrhosis is associated with decompensation^21^. Transcriptomic analysis of colonic tissue demonstrated upregulation of genes associated with necroptosis, apoptosis and inflammation in BDL animals. Functional analyses pointed to modulation of colonic inflammation by Yaq-001, IL-17 signaling, which is known to have diverse biological functions, promoting protective immunity against many pathogens, neutrophil recruitment, antimicrobial peptide production and enhanced barrier function^31, 32^.

Yaq-001 significantly reduced the severity of liver injury and portal hypertension in both models of cirrhosis and ACLF. The lack of significant differences in CPA between untreated and Yaq-001-treated BDL groups suggests that the reduction in portal pressure is possibly due to modulation of the dynamic component of portal hypertension, in which inflammation is known to play a role ^33, 34^ and proposes Yaq-001 as a potential treatment for portal hypertension. Reduction in ALT levels and histology confirmed a reduction in liver injury in the Yaq-001 treated animals. The reduction in liver injury in the LPS treated BDL animals suggests that Yaq-001 has a particular effect on endotoxin sensitivity *in vivo.* This hypothesis was tested in isolated Kupffer cells, which confirmed that LPS-induced ROS production was significantly impacted by Yaq-001 treatment.

Transcriptomic analysis of liver tissue demonstrated that the upregulated genes, CXCL16, CASP1 and TGFB2 in BDL rats was prevented by Yaq-001 administration. Silencing of CXCL16 alleviates hepatic ischemia reperfusion injury and CXCL16 variant is also associated with Hepatitis B virus related acute liver failure^35^. CASP1 mediates pro-inflammatory cytokine release and pyroptotic cell death in cirrhosis and its inhibition has been shown to prevent ACLF ^36^. TGFB2 is an important mediator of cellular senescence^37, 38^. Of note, Yaq-001 also modified necroptosis and cytosolic DNA-sensing pathways representing cell death. Both pyroptosis and necroptosis are known to be activated by LPS and are immunogenic forms of cell death that can trigger further cell death and lead to systemic inflammation^39^. These effects of Yaq-001 potentially explains the effect of Yaq-001 in reducing liver injury ^40, 41^.

Yaq-001 administration had a significant impact on time to coma of ACLF rats, which is considered as a surrogate for mortality compared to untreated controls. Yaq-001 also significantly lowered portal venous and arterial ammonia levels, which was associated with reduced brain water. Transcriptomic analysis of brain tissue showed that IL-18, TGFB2, CCR5 and IL-23A were dysregulated in BDL rats and these were corrected by Yaq-001. IL-18 is released during pyroptosis by activation of the inflammasome complex in neuroinflammatory and neurodegenerative diseases^42^. The effect of Yaq-001 on TGFB2 may mean that it has an impact on senescence, which is known to be associated with hepatic encephalopathy. CCR5 has been implicated in neuroprotection and is novel therapeutic target in stroke^43^. The impact of Yaq-001 on IL-23A indicates possible reduction in neuroinflammation.

In both cirrhosis and ACLF models, Yaq-001 reduced renal dysfunction. Transcriptomic analysis of kidney tissue showed that CCL24 was downregulated in BDL rats, which was prevented in the Yaq-001-treated animals. CCL24 protects renal function in the development of early diabetic nephropathy by exerting an anti-inflammatory effect^44^. Yaq-001 impacted, in particular on the cytokine-cytokine receptor interactions and chemokine and toll-like signaling pathways, which were abnormal in the BDL rats.

BDL animals become sarcopenic and lose weight^45^, which was significantly abrogated by Yaq-001. The possible mechanisms underlying this effect are likely multifactorial^46^. Yaq-001 reduced ammonia significantly, which has been shown to induce sarcopenia. Weight loss in cirrhosis is also attributed to an increased catabolic state in the context of systemic inflammatory response and thus the observed improvement in body weight may reflect the diminished catabolic state with reduced inflammation^46^. These data further emphasize the lack of deleterious effects on nutritional status, but it is not possible to comment on any effect on micronutrients and vitamins, which will need to be explored in future studies.

The data correlating the changes in the microbiota induced by administration of Yaq-001 with changes in the gene expression of multiple relevant pathways is particularly important as it achieves the beneficial effects in the distant organs such as the liver, brain and kidneys with demonstrable changes without leaving the gut. The exact mechanisms by which this occurs cannot be directly inferred from the data derived from this study. One possibility is that alongside LPS adsorption and modulation of other unmeasured toxins, the *milieu* of the gut is changed allowing proliferation of more autochthonous bacteria^47^, which impacts on gut inflammation that reduces gut permeability leading to a reduction in endotoxemia, systemic and organ inflammation, organ priming, improvement of organ function and LPS-sensitivity. In this study, most of these changes have been described, but whether this is happening in sequence has not been studied.

Limitations of this study include potential underpowering of the study for 16S rRNA gene studies. The rodent microbiome is not directly analogous to the human and further clinical studies will be required to verify the effects on the gut microbiome’s bacterial composition.

Although Yaq-001 was effective in adsorbing a variety of bile acids *in vitro* and reduced bile acids significantly in Sham animals, no impact on bile acids was seen in BDL animals. This possibly reflects the effect of the BDL model, where no increase in bile acids was observed. Studies in other models will be needed to determine the role of Yaq-001 in modulating bile acid metabolism. Although, Yaq-001 was observed to impact positively on the gene expression profiles of multiple pathways, their exact relevance at the protein or cellular level has not been explored.

In conclusion, the data provides compelling evidence for the potential of Yaq-001 as a novel therapy targeting the gut microbiome, bacterial translocation and gut permeability that impacts on systemic inflammation and organ function in models of cirrhosis and improves survival in ACLF. Translation to clinical studies is warranted to further assess its safety and efficacy.

## Abbreviations

ACLF: acute-on-chronic liver failure
LPS: lipopolysaccharides
BDL: bile duct ligation
ALT: alanine aminotransferase
ALP: alkaline phosphatase
TBIL: total bilirubin
MAP: mean arterial pressure
CPA: collagen proportionate area
PSR: picrosirus red
PP: portal pressure
ROS: reactive oxidant species
DEGs: differential expressed genes
KEGG: Kyoto Encyclopedia of Genes and Genomes
TLR: toll-like receptor
TNF-a: tumor necrosis factor-a.

## Acknowledgements

We would like to thank Fraser Simpson (Department of Genetics, Evolution & Environment, University College London) for his support to perform the Nanostring. The urinary NMR studies were facilitated by a Medical Research Council research grant under the High-throughput "omic" Science and Imaging funding scheme [MC_PC_13045].

