## Supplementary Data for "Impact of Yaq-001, a non-absorbable, engineered carbon bead of controlled porosity in rodent models of cirrhosis and acute on chronic liver failure"

**Table of contents**

Supplementary materials and methods----------------------------------------------2

Fig.S1----------------------------------------------------------------------------------17

Fig.S2----------------------------------------------------------------------------------19

Fig.S3----------------------------------------------------------------------------------20

Fig.S4----------------------------------------------------------------------------------22

Fig.S5----------------------------------------------------------------------------------24

Fig.S6----------------------------------------------------------------------------------27

Fig.S7----------------------------------------------------------------------------------29

Table S1-------------------------------------------------------------------------------30

Table S2-------------------------------------------------------------------------------66

Table S3-------------------------------------------------------------------------------67

Table S4-------------------------------------------------------------------------------78

Table S5-------------------------------------------------------------------------------86

Supplementary references-----------------------------------------------------------91

**Supplementary materials and methods**

**STUDIES *IN VITRO***

***Adsorption studies***

Adsorption of biomolecules of varying molecular weight (albumin, myoglobin and caffeine).For each of the molecular weight size markers, the kinetics of adsorption was studied compared to that of no carbon control. For endotoxin adsorption, the carbon samples were sterilised at 200ºC for 3 hours. Endotoxin solution was added to each carbon sample and incubated on a shaking plate. In separate studies, different bile acids were spiked. The adsorption kinetics of endotoxin and bile acids were measured using the chromogenic limulus amebecyte lysate assay and uv/vis spectroscopy respectively.

*Materials for the adsorption studies*

Yaq-001 was obtained from Yaqrit Ltd, UK. Bovine serum albumin (Cat. No. A6003), myoglobin from equine skeletal muscle (Cat. No. M0630), caffeine (Cat. No. C0750), bicinchoninic acid solution (Cat. No. B9643), copper (II) sulfate pentahydrate 4% solution (Cat. No. C2284), and phosphate buffer solution, pH= 7.4 (Cat. No. P3619) were procured from Sigma-Aldrich (St Louis, MO, USA).

Cholic acid (Cat. No. C1129), deoxycholic acid (Cat. No. D2510), lithocholic acid (Cat. No. L6250), chenodeoxycholic acid (Cat. No. C9377), sodium glycocholate (Cat. No. G7132) and sodium glycochenodeoxycholate (Cat. No. 50534)] were ordered from Merck (Darmstadt, Germany).

Simulated intestinal fluid solution (SIF) was prepared by dissolving 6.8 g of monobasic potassium phosphate in 250 mL of water and then adding 77 mL of 0.2 N sodium hydroxide and 500 mL of water. The resultant solution was adjusted to a pH of 6.8 ± 0.1 using either 0.2 N sodium hydroxide or 0.2 N hydrochloric acid, then diluted to 1000 mL with water.

*Albumin (66.7 kDa) adsorption kinetics*

For the kinetics study, 0.1g of Carbon adsorbent was pre-wetted into microcentrifuge tubes (15 mL capacity) with PBS under vacuum for 3 hours at -1000 mbar pressure. Pre-wetting solutions were removed by micropipette and 5 mL of bovine albumin solution (2 mg/mL) was added to each tube. Samples were incubated at 37°C on orbital shaker for 24h at 120 rpm. The albumin concentration was measured by collecting supernatants following centrifugation for 2 minutes at 1600 g at 0, 0.5, 1, 2, 3, 5 and 7 hours. Albumin concentration was measured using a BCA method according to manufacturer's instructions. The BCA working reagent was prepared by mixing 50 parts of reagent A to 1 part reagent B to give a light green colour. Albumin standards concentrations of 0, 0.1, 0.2, 0.4, 0.6, 0.8, 1, 1.5 and 2 mg/mL in PBS were used to prepare a standard curve to calculate unknown albumin concentration. 2 mL of BCA working reagent was mixed with 0.05 mL of albumin sample, standard or blank. The solutions were Incubated at 37 ºC for 30 minutes, transferred into a cuvette and absorbance was measured using a Jenway 6705 UV/Vis spectrophotometer (Bibby Scientific Ltd, UK) at 562 nm by UV spectrophotometry.

*Myoglobin (16.7 kDa) adsorption kinetics*

For the kinetics study, 0.1g of carbon adsorbent samples were pre-wetted in triplicate into 15 mL microcentrifuge tubes with PBS under vacuum for 3 hours at -1000 mbar pressure. Pre-wetting solution was removed, 5 mL of myoglobin solution (500 ug/mL) was added and samples were incubated at 37°C on orbital shaker for 24h at 120 rpm. Supernatant were collected at 0, 0.5, 1, 2, 3, 5 and 7 hours, centrifuged for 2 minutes at 1600 g and transferred to a sterile universal tube for measurement of myoglobin concentration. 3 mL of supernatant sample from controls and experimental samples was placed in a cuvette and absorbance was measured using a Jenway 6705 UV/Vis spectrophotometer at a wavelength of 409 nm. A standard curve was prepared using myoglobin standard ranging from 0-500 µg/mL.

*Caffeine (0.194kDa) adsorption kinetics*

For the kinetic study, 0.1g of carbon adsorbent samples were pre-wetted in triplicate into 15 mL microcentrifuge tubes with PBS under vacuum for 3 hours at -1000 mbar pressure. 5 mL of a 10 mg/mL caffeine solution was added into each tube and incubated at 37°C on an orbital shaker for 24h at 120 rpm. 3 mL of supernatant was collected into clean microcentrifuge tubes at 0, 0.5, 1, 2, 3, 5 and 7 hours following centrifugation for 2 minutes at 1600 g for measurement of residual concentration. The caffeine concentration was measured using a Jenway 6705 UV/Vis spectrophotometer (Bibby Scientific Ltd, UK) at a wavelength of 273 nm. A standard curve was prepared using a concentration range of 0 to 10 mg/mL. Standards and samples were diluted in methanol (1:100) and 3 mL of supernatant was added to a quartz cuvette for analysis.

*Endotoxin adsorption*

All carbon samples were sterilised at 200ºC for 3 hours. 0.1 g sample was weighed into pyrogen free glass vials in triplicate. 4 mL spiked endotoxin solution (10 EU/mL) in SIF was added to each carbon sample and incubated on a shaking plate (Stuart Equipment, UK) at 37ºC for 90 minutes. 450 µL of sample was collected at 0, 15, 30, 45, 60 and 90 minutes for the kinetic study. The standards (50, 5, 0.5, 0.05, 0.005 EU/mL) were made according to manufacturer's instructions in endotoxin free glass tubes using a 5mL volume for each dilution and vortexing well between each dilution. 0.1 mL of standard, sample or control in duplicate was added to the wells of a 96 well plate. 0.1 mL LAL reagent was prepared and immediately added to each well. The plate was placed in an incubating microplate reader (Biotek Instruments, USA) that measures absorbance at 405 nm and monitored over time using Gen5 2.0 software.

*Bile acids*

For the adsorption kinetics study, series of carbon adsorbent samples (0, 0.025, 0.05, 0.1, 0.25 & 0.5 g) were pre-wetted in triplicate into 15 mL microcentrifuge tubes with PBS under vacuum for 3 hours at -1000 mbar pressure. Pre-wetting solution was removed, 5 mL of bile acid spiked simulated intestinal fluid (SIF) solution [Cholic acid (150 μM), Deoxycholic acid (200 μM), Lithocholic acid (800 μM), Chenodeoxycholic acid (800 μM), Sodium glycholate (800 μM) and Sodium glycochenodeoxycholate (354 μM)] was added and samples were incubated at 37°C on orbital incubator shaker S150 (Stuart Equipment, UK) for 24h at 120 rpm. Supernatants were collected after 24 hours, centrifuged for 2 minutes at 1600 g and transferred to a sterile universal tube for measurement of bile acid concentration. 3 mL of supernatant sample from controls and experimental samples was placed in a cuvette and absorbance was measured using a Jenway 6705 UV/Vis spectrophotometer (Bibby Scientific Ltd, UK) at a wavelength of 405 nm. A standard curve was prepared using respective bile acid standards ranging from 0-1000 μMol/L.

**Bacterial growth in the presence of Yaq-001**

The effect of activated carbons on the kinetics of bacterial growth was studied for *S. aureus* and *E. coli.* 5 mL of overnight culture was prepared aerobically by standard technique at 37°C, which was then transferred to a 15 mL falcon tube and centrifuged at 4000 g for 10 minutes at 4°C.The bacterial pellet was re-suspended in 5 mL of phosphate buffered saline (PBS). 0.1 mL was added to 100 mL of Tryptone soya broth (TSB) to achieve a final concentration of $\sim 10^6$ CFU/mL. The bacterial cultures supplemented with 10 mg/mL of activated carbon samples were incubated for 6 hours in triplicate at 37°C whilst shaking at 150rpm. 1 mL aliquots were withdrawn from the culture medium and the optical density at 600 nm was measured every 30 minutes in triplicate. Comparison of the growth curves obtained in the absence of carbon samples and amoxicillin antibiotic (control) were evaluated. The carbon beads did not alter the bacterial growth kinetics of *E. coli* and *S. aureus* cultures over 6 hours.

**Scanning electron microscopy analysis**

Activated carbon samples were mounted on aluminium stubs using carbon adhesive tape and coated with a 2nm thick layer of platinum using a Quorum Q150TES coater (Quorum Technologies, UK). The surface as well as internal morphology was examined using a Zeiss Sigma field emission field emission scanning electron microscope (FESEM) (Carl Zeiss Microscopy, Germany) at an accelerating voltage of 5 kV at 100, 500 and 50000x magnifications.

**Pore size distribution by mercury porosimetry**

Mercury porosimetry analysis was performed using a mercury porosimeter PoreMaster (Quantachome Instruments, USA). Prior to the analysis, all materials were dried in a vacuum oven at 110°C under 800 bar vacuum for 3 hours. The meso- (2-50nm) and macro- (diameter ≥50 nm) pore size distributions of the AC beads were determined by mechanical intrusion of mercury. Data was analysed using PoreWin 6.0 software (Quantachrome Instruments, USA).

**Bile duct ligation model**

All animal experiments were conducted according to Home Office guidelines under the UK Animals in Scientific Procedures Act 1986. Male Sprague-Dawley rats (body weight 280-300 g, age 8-10 weeks) were used (Charles River Laboratories UK Ltd.). All rats were housed in the unit and given free access to standard powdered rodent chow and water, with a light/dark cycle of 12 hours, at a temperature of 19°C to 23°C and humidity of approximately 50%.

Under halothane anesthesia 194 rats underwent bile duct-ligation (BDL) or Sham biliary surgery by randomisation. The rat model used to mimic ACLF in this study was published and described by Harry D *et al*[1].Rats were pair-fed powered chow -/- pre-hydrated Yaq-001 carbon (250-500 μm) at a dose of 0.4 g/100 g body weight per day from 2 weeks after bile duct ligation until completion of the experiment at 4 weeks from initial surgery. Intraperitoneal *Klebsiella* lipopolysaccharide (LPS) (0.33 mg/kg) was administered to 4 subgroups 6 hours prior to completion of study. The following groups were studied: Sham (n=36), Sham-Yaq-001 (n=30), Sham-LPS (n=9), Sham-LPS-Yaq-001 (n=10), BDL (n=37), BDL-Yaq-001 (n=44), BDL-LPS (n=16), BDL-LPS-Yaq-001 (n=12). All rats had biliary cirrhosis. None had ascites as at 4 weeks after BDL, ascites is not observed.

**Haemodynamic measurements**

Under halothane anaesthesia (5 mL/min induction 2 mL/min maintenance) an internal carotid catheter (0.96 outer diameter Portex fine-bore polythene tubing, Scientific Laboratory Supplies Ltd., Nottingham, UK) was inserted as previously described[2]. The catheter was held in place for the duration of the study by both proximal and distal holding sutures. The catheter was transduced and mean arterial pressure determined. A laparotomy was performed under sterile conditions and a catheter placed in the portal vein. Arterial and portal venous catheters were transduced. Measurements were transduced to a Powerlab (4SP) linked to a computer with Chart v5.0.1 software.

***Blood and tissue sampling***

Blood samples were taken from abdominal aorta or right heart. EDTA plasma was centrifuged at 1000 g for 10 minutes and stored at −80°C. All tissues (liver, brain, kidneys and colon) were snap-frozen in liquid nitrogen and stored at −80°C. Organs were harvested in formalin (48 h) for histological assessment.

**Biochemical analysis**

Biochemical profile was determined using standard techniques (COBAS).

**D-lactate assay**

Plasma was collected in EDTA tubes from rats in different groups and analyzed using the D-Lactate Assay Kit (Colorimetric, ab83429) according to the manufacturer's protocols.

**Brain water percentage**

The entire brain was weighed immediately after sacrifice using an electronic balance to determine the wet weight. The brain was dried in an oven at 100°C for 24 hours to obtain the dry weight. The BW content was calculated according to the formula: BW content (%) = (Wet weight - Dry weight) / (Wet weight) × 100.

**Ammonia levels**

Standard operating procedures for ammonia measurement that involved collection of the sample in cooled EDTA tubes, rapid sample transport to the laboratory on ice and spectrophotometric assays. Plasma arterial and portal venous ammonia was detected by using Fujifilm Dri-Chem NX500 (Fujifilm Corporation, Japan) instrument and related cartridges.

**Bacterial DNA isolation from plasma**

An aliquot (100 μL) of fresh or thawed plasma was added to 600 μL of chilled Nuclei Lysis Solution and homogenized for 10 seconds. This was followed by a 15-30 minutes incubation step at 65°C. 3 μL of RNase solution was added to the lysate, mixed and incubated for 15-30 minutes at 37°C. The solution was cooled to room temperature, 200 μL of Protein Precipitation solution added and subsequently chilled on ice for 5 minutes. This solution was centrifuged at 13000-16000 g for 4 minutes. The supernatant was transferred to a fresh tube containing 600 μL of room temperature isopropanol and mixed gently by inversion. The samples were centrifuged at 13000-16000 g for 1 minute, supernatant removed and 600 µL of room temperature 70% ethanol added. Following mixing and a further centrifugation step, the ethanol was aspirated and pellet allowed to air dry for 15 minutes. The DNA was re-suspended in 100 µL of DNA Rehydration solution overnight at 4°C.

**DNA amplification and sequencing (plasma samples)**

Two µL of DNA template was added to a reaction mix containing: 10 mmol/L Tris buffer (pH 8.3), 50 mmol/L KCl, 1.5mmol/L $MgCl_2$, 200 mol/L of each deoxynucleoside triphosphate, 50 pmol of primers 5_-AGAGTTTGATCATGGCTCAG-3_ and 5_ACCGCGACTGCTGCTGGCAC-3_, 1.25 U BioTaq (Bioline, London, England) to complete a final volume of 50 µL. A 35-cycle PCR was run in GeneAmp 9700 (Applied Biosystems, Foster City, CA) using the following profile: 94°C for 30 seconds, 55°C for 30 seconds, 72°C for 60 seconds. The total PCR reaction volume was filtered with QIAquick Spin Columns (QIAquick PCR Purification Kit; QIAGEN, West Sussex, UK) to remove rests of primers. Five µL of purified product was analyzed by 2% agarose gel electrophoresis and UV visualization. A band of about 540 base pairs was obtained from different bacterial cultures corresponding to the specific amplification of the prokaryotic 16S rRNA gene.

**Histological analysis**

Liver tissue was processed in accordance with standard protocol and Haematoxylin and Eosin together with Sirius Red staining was performed. Sirius red staining was quantified using computer assisted digital image analysis. Collagen proportionate area was determined using Zeiss KS300 image analysis software.

**TUNEL assay**

Enzymatic in situ labeling of cell death was assessed by In Situ Cell Death Detection Kit, POD (11684817910, Roche, Basel, Switzerland). Briefly, PFFE liver slides were dewaxed (3x xylene, 5 minutes) and rehydrated (Ethanol, 95%, 90%, 80%, 70% and dd$H_2O$, 3 minutes each). After a wash in PBS, the tissues were surrounded by permanent histology pen and incubated with 150 µL Proteinase K (03115887001, Roche, Basel, Switzerland) solution 20 µg/mL in 10 mM Tris/HCl, pH 7.4-8 for 30 minutes. Following, the slides were incubated with 0.1% Triton X–100, 0.1% sodium citrate (T8787, 1613859, Sigma-Aldrich, Saint Louis, MO, USA), freshly prepared for 8 minutes. The slides were rinsed twice with PBS and incubated with 100 µL TUNEL reaction mixture for 60 minutes at -37°C in a humidified atmosphere in the dark. The slides were washed again in PBS twice and incubated with 100 µL Converter-POD in a humidified chamber for 30 minutes at 37°C. The samples were washed with PBS and 100 µL of POD substrate was added for 10 minutes. Nuclei were counterstained with hematoxylin (HHS16, Sigma-Aldrich, Saint Louis, MO, USA). The slides were dehydrated and mounted.

**Isolation of non-parenchymal cell fraction from rodent liver tissue**

Perfused liver tissue was dissected with a scalpel and homogenized in Hanks balanced salt solution (with calcium and magnesium - collagenase 0.01% and DNAse I (0.01%). The homogenate was transferred to a 50 mL Falcon tube and incubated at 37°C prior to filtration through a cell strainer (100 mm for rat tissue). This was centrifuged at 150 g for 5 minutes at 4°C and the supernatant subsequently centrifuged at 800 g for 10 minutes at 4°C. The supernatant was discarded and the pellet re-suspended in PF4 (HBSS with no calcium or magnesium, DNAse I 0.01%, bovine serum albumin (0.25%)) and centrifuged at 800 g for 10 minutes at 4°C. The pellet was re-suspended in 3.9 mL of RPMI 1640 and mixed gently with 2.1 mL (RPMI and Optiprep 22%). RPMI was layered on top followed by a 25 minute centrifugation step at 850 g without brake at 4°C. The non-parenchymal cells were isolated from the interface, re-suspended in an equivalent volume of PF4 and centrifuged at 800 g at 4°C for 10 minutes.

The pellet was re-suspended in 5 mL of Red Cell Lysis (RBC) buffer (BioLegend) and incubated for 5 minutes at 4°C with occasional shaking. The reaction was stopped by addition of 10 mL of PBS. The pellet was centrifuged at 800 g for 10 minutes at 4ºC, the supernatant discarded and the cells re-suspended in 3 mL of culture media. The cells were counted and adjusted to a concentration of $10^7$ cells/mL. $1x10^6$ cells were used in all subsequent assays.

**Whole blood preparation**

Two mL of blood was collected from portal vein and arterial blood. 40mL of RBC lysis buffer was added to each 2 mL sample, vortexed and incubated for 15 minutes at room temperature. Following centrifugation at 450 g for 5 minutes the supernatant was discarded and the pellet re-suspended with 1 mL of complete culture media (RPMI (Gibco), 10% FBS, 200 mM L-Glutamine). The cell number using the nucleoCounter method and adjusted to $1x10^7$ cell/mL.

**Kupffer cell population studies**

Two uL of Fc blocker (anti-CD32 antibody) was added to $10^6$ cells (non-parenchymal cell fraction) and incubated for 5 minutes at 4°C. The cells were co-incubated with anti-CD163 antibody for 30 minutes at 4°C in the dark**.** The cells were washed with 1 mL of FACS buffer, centrifuged at 800 g and re-suspended in 100 mL FACS buffer solution and analyzed immediately on a Becton Dickinson LSR II flow cytometer.

**Cell population assay**

Two µL of Fc blocker (anti-CD32 antibody) was added to $10^6$ cells and incubated for 10 minutes at 4°C. The cells were co-incubated with the following primary antibodies for 20 minutes at 4°C: PE-CD11b (0.2 mg/mL), Alexa647-CD43 (0.5 mg/mL), FITC-HIS48 (200 µg/mL). After incubation, 2 mL of FACS buffer was added to each tube, centrifuged at 450 g for 5 minutes at 4°C and supernatant discarded. This last step was repeated with 1 mL FACS solution and the pellet re-suspended the pellet in 100 µL of FACS buffer. 5 µL of DAPI solution was added into the each tube prior to analysis.

**Kupffer cell and portal venous ROS studies**

20 µg/mL of *E. coli* endotoxin was added to $1x10^6$ non-parenchymal cells sample and incubated for 30 minutes at 37°C. ROS inducer at a final concentration of 200-500 µM was used as a positive control. The samples were centrifuged at 500 g for 5 minutes and the supernatant discarded. The cells were re-suspended in 5 mL of wash buffer, centrifuged at 500 g for 5 minutes and the supernatant removed. The cells were re-suspended in 500 µL of ROS detection solution and incubated for 30 minutes at 37°C in the dark. Following centrifugation, the cells were re-suspended in 100 µL of FACS buffer, Fc blocker added (1:25) and incubated for 10 minutes at 4°C. Anti-CD163 antibody was added and the cells incubated for 30 minutes at 4°C in the dark (Kupffer cells). The cells were washed with 1 mL of FACS buffer, centrifuged at 800 g and re-suspended in 100 mL FACS buffer solution. All samples were kept at 4°C and analyzed immediately using a FACS LSR II machine. Data was analyzed using FlowJo software.

**Cytokine analysis**

Plasma cytokine levels were measured in EDTA anti-coagulated plasma using a Bio-Plex Pro rat cytokine assay kit (R & D systems, USA) and a Bio-Plex Magpix instrument (Bio-Rad Laboratories Ltd., Watford, UK) according to the manufacturer's instructions. The cytokines measured were IL-1β, IL-6, IL-10 and TNF-α.

**Endotoxin measurement**

The chromogenic limulus amoebocyte lysate kinetic assay (Charles River Laboratories) was used for the detection of endotoxin. Portal venous plasma

(100 µL) was diluted 1:10 with endotoxin-free water and incubated at 75°C for 30 minutes. 100 µL of sample and 100 mcl of LAL reagent were mixed in a 96-well plate and analyzed at 405 nm with spectrophotometer using the Endoscan V software. Results are expressed as EU/mL.

**Sequence-based microbiota composition determination**

Analysis of faecal microbiota was performed as previously described[3]. In brief, DNA was extracted from 200 mg of faecal pellets, one pellet per animal, and DNA was combined within treatment groups. Total DNA was extracted using an initial bead-beating step and the QIAamp DNA stool mini kit (Qiagen, West Sussex, UK). Universal 16S rRNA gene primers, designed to amplify from highly conserved regions corresponding to those flanking the V4 region, i.e. the forward primer F1 (5′-AYTGGGYDTAAAGNG) and a combination of four reverse primers R1 (5′-TACCRGGGTHTCTAATCC), R2 (5′-TACCAGAGTATCTAATTC), R3 (5′-CTACDSRGGTMTCTAATC) and R4 (5′-TACNVGGGTATCTAATC) (RDP's Pyrosequencing Pipeline: http://pyro.cme.msu.edu/pyro/help.jsp) were used for Taq-based PCR amplification. Sequencing was performed on a Roche 454 GS-FLX using Titanium chemistry by the Teagasc 454 Sequencing Platform (Teagasc, Fermoy, Ireland). Resulting reads were quality trimmed, clustered, aligned and checked for chimeras using the Qiime suite of tools. The reads for the major phyla were averaged for each group and expressed as a percentage of the total number of reads for that particular group.

We performed differential abundant analysis based on the bacterial abundance between groups, the bacteria with the p value of pairwise test lower than 0.5(Wilcoxon Rank Sum Test, $p<0.5$, Sham vs BDL and BDL vs BDL-Yaq-001) or the bacteria with greater than four-fold variation between groups ($\log2FC>2$, Sham vs BDL and BDL vs BDL-Yaq-001) were selected into visualization at both genus and family level.

### RNA isolation

Liver, colon, brain and kidneys from three rats in Sham, Sham-Yaq-001, BDL groups and four in BDL-Yaq-001 group were randomly selected to perform transcriptome analysis. Total RNA was isolated and cleaned up by using QIAzol Lysis Reagent (79306 Qiagen, CA, USA), and RNeasy Mini Kit (74104 Qiagen), respectively, according to the manufacturer's protocol. RNA integrity was analyzed with the Agilent RNA 6000 Nano chip on the Agilent 2100 Bioanalyzer and only samples with R.I.N. above 8 were kept.

### NanoString gene expression analysis

For NanoString gene expression analysis, we used 100ng RNA for inflammation profiling with a Customized NanoString nCounter CDR-RatCirrh-21590 panel including 167 genes (NanoString Technologies, Table S7). Each 5μL RNA sample was hybridized with 8 μL nCounter Reporter probe in hybridization buffer, and 2 μL nCounter Capture probes at 65°C for 16–30 h. Excess probes were removed through a twostep magnetic bead-based purification procedure using the nCounter Prep Station (NanoString Technologies). Specific target molecule abundance was quantified using the nCounter Digital Analyzer to count individual fluorescent barcodes, whereby the corresponding target molecules were assessed. Each assay involved a high-density scan encompassing 280 visual fields. Images of the immobilized fluorescent reporters in the sample cartridge were acquired using a CCD camera, and data were collected using the nCounter Digital Analyzer. The mRNA data analysis was performed and normalized against housekeeping genes using nSolver4.0 software (NanoString Technologies). Differentially expressed genes (DEGs) were downloaded from nSolver4.0 software following the criteria: $|FC| > 1.2$ and p-value $< 0.1$. The R packages "ComplexHeatmap 2.14.0" and "ggplot2 3.4.1" were used to present the heatmap and volcano plot of DEGs. Kyoto Encyclopedia of Genes and Genomes (KEGG) pathway analysis was performed by using the R package “ClusterProfiler 4.6.0” to reveal the key pathways of inflammation associated with DEGs (criteria: p-value < 0.05, q-value <0.05, significantly enriched).

**Gut microbiome studies**

The calculation of alpha diversity measures used phyloseq package (Version 1.42.0). Kruskal Wallis test was used for the comparison of alpha diversity measurements of four groups. PERMANOVA was used for beta diversity comparison test of groups using anova2 function of vegan package in R (Version 2.6.4). Beta diversity was measured by bray distance, and the distance was visualized by Principal Coordinate Analysis.

**Correlation analysis**

To assess whether microbiome compositional changes correlate with tissue gene expression we calculated the Spearman’s rank correlation coefficient between these datasets. For this purpose, raw metagenomics abundance data was correlated with raw gene expression data across samples using the spearmanr function as part of the scipy package in python3.10.

**$^{1}$H-NMR urinary metabolic profiling**

Rat urine was prepared for $^{1}$H-NMR analysis using a standard protocol[4]. Briefly 400 μL of urine was buffered to pH 7.4 using 200μL of 0.2M phosphate buffer, also containing D2O for an NMR field frequency lock and 3-(trimethylsilyl) propionic-2,2,3,3-d4 acid sodium salt (TSP) for a chemical shift reference and 550 μL of the buffered urine was transferred to a 5mm NMR tube. Urinary NMR spectra were acquired at 298K from using a JEOL ECP 500 MHz NMR spectrometer (JEOL Ltd, Tokyo, Japan). A standard pulse-collect sequence with water presaturation was used to acquire the NMR data. The spectral width was 15 ppm, pulse angle 90°, acquisition time 4.36 s and relaxation delay 3 s. 32K data points were acquired per collect and 64 transients were summated.

The receiver gain was constant for all samples. The resulting free induction decay was zero filled and multiplied by an exponential function corresponding to 0.3 Hz line broadening prior to Fourier Transformation. The NMR spectra were manually phased using the JEOL Delta.

Prior to statistical analysis, all NMR spectra were baseline corrected to a 4th degree polynomial, zero filled by a factor of 2 and referenced with the TSP peak set to 0.00 ppm using KIA version 9.0 (Bio-Rad, Philadelphia, USA). NMR spectral resonances were assigned according to the literature[5].

The resonances attributable to residual water and urea (δ 4.6– 6.4 ppm) were excluded from further analysis. NMR spectra were normalised to the total spectral integral in the range δ =0.2–10 ppm (excluding 4.6– 6.4 ppm). NMR spectra were bucketed (total buckets 603) using the Intelligent Bucketing algorithm and mean centred. Principal Components Analysis (PCA) was used as an unsupervised method for data visualisation and outlier identification.

**Study approval**

All animal experiments were in accordance with UK Home Office Animals (Scientific Procedures) Act 1986 (updated 2012) and a project license (No. 14378) provided by the UK Home Office.

### Supplementary figures

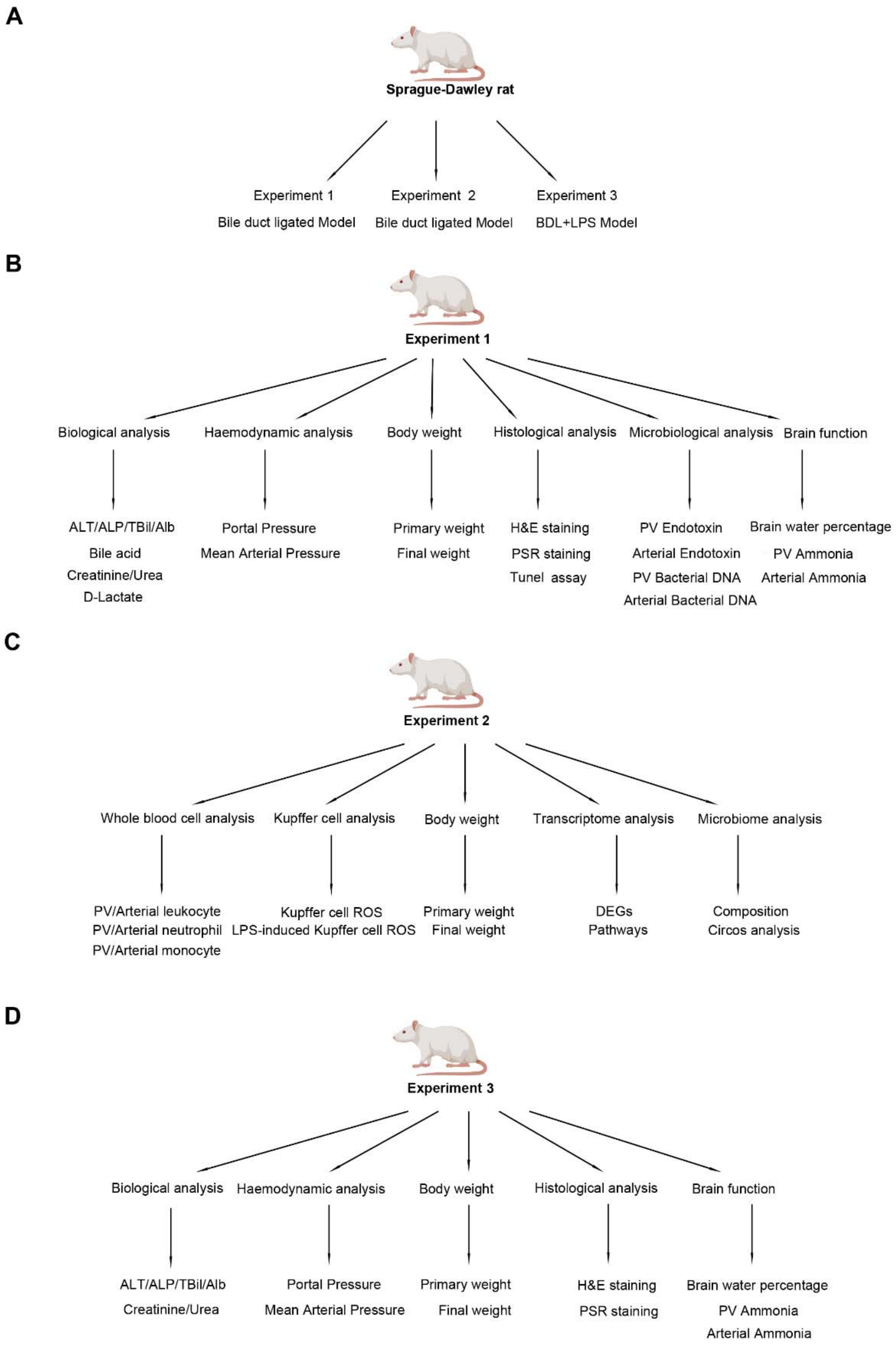


***Fig.S1. Study outline for investigating the efficacy of Yaq-001 in cirrhosis and ACLF.***

*(A) Our study consisted of three independent experiments. (B, C) Design of Experiments 1 and 2 in the four-week bile duct ligated (BDL) rat model, which was used as a model of cirrhosis. (D) Design of Experiment 3 in the four-week BDL rat model with a single intraperitoneal injection of Klebsiella pneumoniae lipopolysaccharide, 6 hours prior to final surgery which was used as a model of ACLF.*

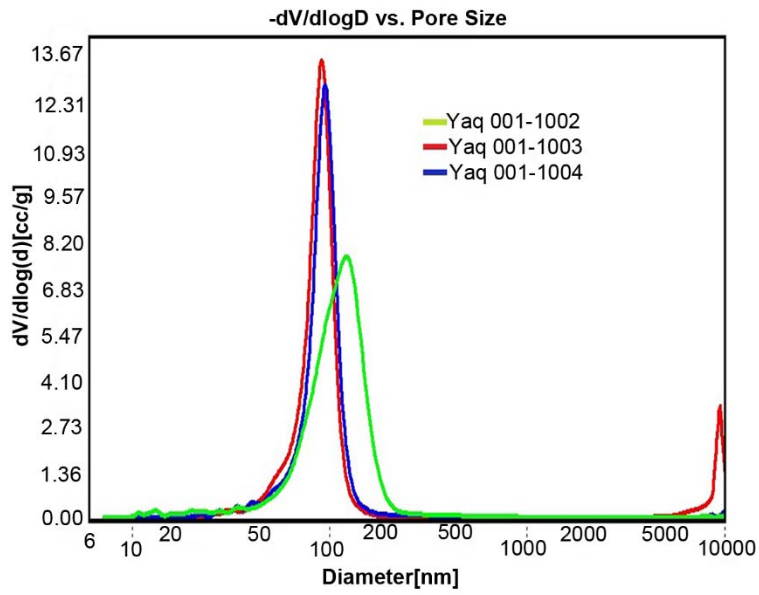


***Fig.S2. Mercury porosimetry analysis of Yaq-001.***

*Yaq-001 1002, Yaq-001 1003, and Yaq-001 1004 indicated different batches of Yaq-001. The different batches of Yaq-001 (1002, 1003 & 1004) produced a consistent pore size distribution plot in the meso-macroporous range from 30-200nm.*

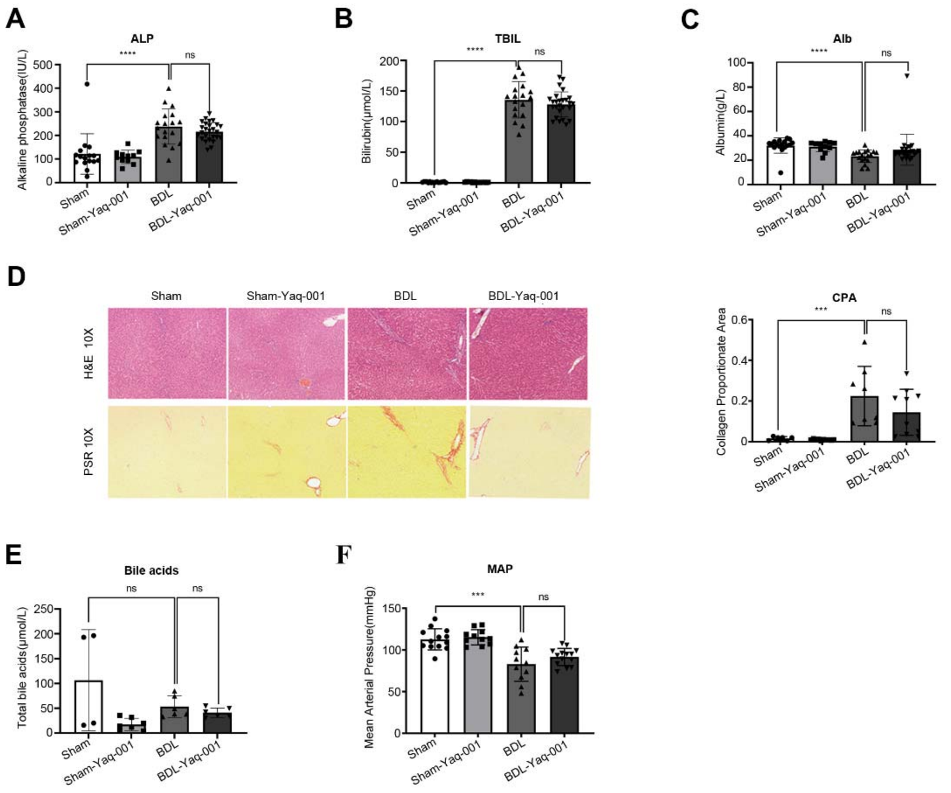


***Fig.S3. Biochemical profiles of Yaq-001 treatment in cirrhotic rats.***

*(A) Plasma alkaline phosphatase (ALP) concentrations in Sham (n=16), Sham-Yaq-001 (n=11), BDL (n=18) and BDL-Yaq-001 (n=26) groups. Significantly higher ALP concentrations were observed in BDL compared to Sham controls (p=0.0002). (B) Plasma total bilirubin (TBIL) concentrations in Sham (n=17), Sham-Yaq-001 (n=13), BDL (n=18) and BDL-Yaq-001 (n=26) groups. Significantly higher bilirubin concentrations were observed in BDL compared to Sham controls (p<0.0001). (C) Plasma albumin levels in Sham (n=24), Sham-Yaq-001 (n=22), BDL (n=18) and BDL-Yaq-001 (n=26) groups. Significantly lower albumin levels were observed in BDL compared to Sham controls (p<0.0001). (D) Haematoxylin & Eosin and PicroSirius Red staining of liver tissue with Collagen Proportionate area (CPA) in cirrhotic rats. BDL was associated with a significant increase in CPA compared to Sham controls (p<0.0001). Yaq-001 had no effect on CPA in either group. (E) Plasma total bile acids concentrations in Sham (n=4), Sham-Yaq-001 (n=7), BDL (n=6) and BDL-*

*Yaq-001 (n=6) groups. No significant difference was observed in BDL compared to Sham controls. Yaq-001 treated Sham rats had significantly lower bile acid concentration compared to untreated Sham rats ($p<0.05$). (F) Mean arterial pressure (MAP) measurements in Sham (n=13), Sham-Yaq-001 (n=11), BDL (n=11), BDL-Yaq-001 (n=14). Significantly lower MAPs were observed in BDL compared to Sham controls ($p<0.001$). Yaq-001 treatment had no effect on MAP.*

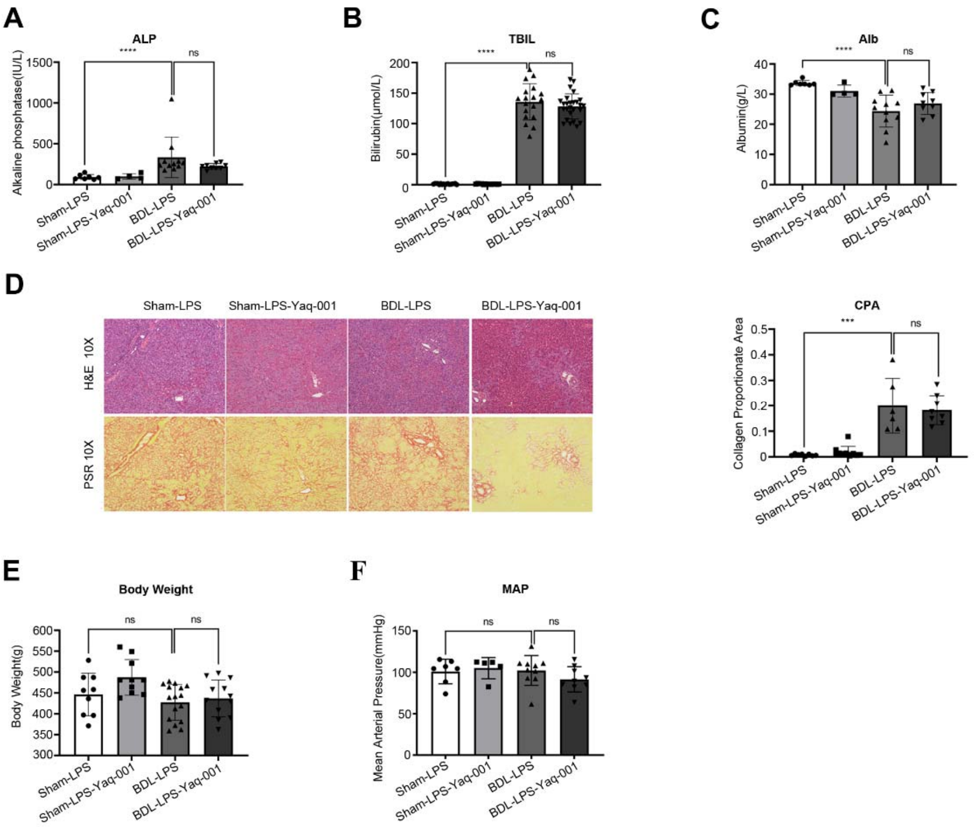


***Fig.S4. Biochemical profiles of Yaq-001 treatment in ACLF rats.***

*(A) Plasma ALP concentrations in Sham-LPS (n=7), Sham-LPS-Yaq-001 (n=4), BDL-LPS (n=11) and BDL-LPS-Yaq-001 (n=9) groups. Significantly higher ALP concentrations were observed in BDL-LPS compared to Sham-LPS controls (p=0.023). (B) Plasma total bilirubin concentrations in Sham-LPS (n=7), Sham-LPS-Yaq-001 (n=4), BDL-LPS (n=11) and BDL-LPS-Yaq-001 (n=9) groups. Significantly higher bilirubin concentrations were observed in BDL-LPS compared to Sham-LPS controls (p<0.0001). (C) Plasma albumin levels in Sham-LPS (n=7), Sham-LPS-Yaq-001 (n=4), BDL-LPS (n=11) and BDL-LPS-Yaq-001 (n=6) groups. Significantly lower albumin levels were observed in BDL-LPS compared to Sham-LPS controls (p=0.0004). Yaq-001 treatment had no effect on albumin. (D) Haematoxylin & Eosin and PSR staining of liver tissue in ACLF rats. BDL-LPS was associated with a significant increase in CPA compared to Sham-LPS controls (p=0.0002). Yaq-001 had no effect on CPA in*

*BDL-LPS rats. (E) 4-week body weights in Sham-LPS (n=9), Sham-LPS-Yaq-001 (n=10), BDL-LPS (n=16) and BDL-LPS-Yaq-001 (n=12) groups. Yaq-001-treated Sham-LPS and BDL-LPS rats had a slightly higher body weights compared to untreated rats. (F) MAP measurements in Sham-LPS (n=7), Sham-LPS-Yaq-001 (n=5), BDL-LPS (n=10), BDL-LPS-Yaq-001 (n=9). LPS and Yaq-001 treatment had no effect on MAP.*

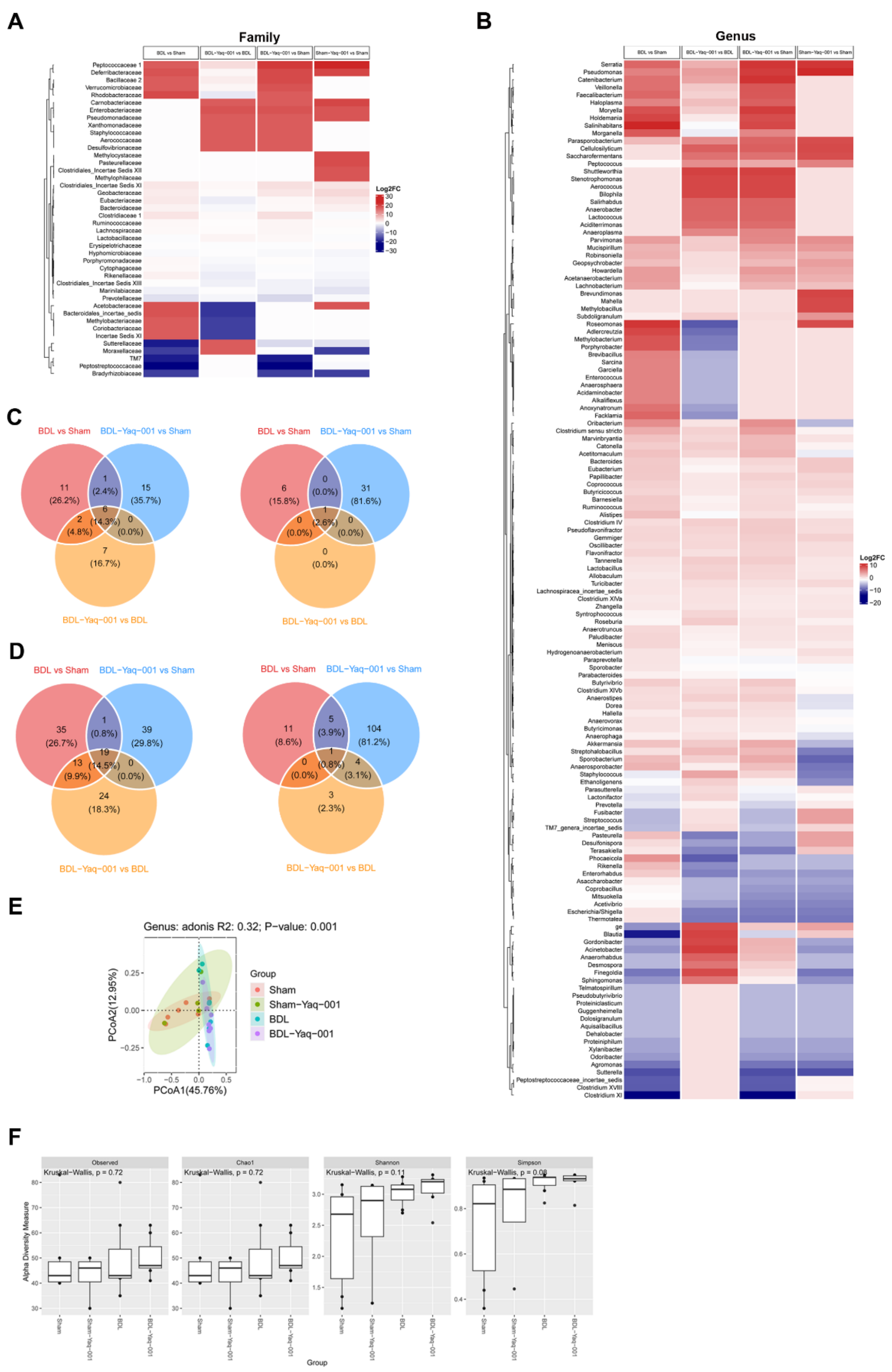


***Fig.S5. Yaq-001 treatment in cirrhotic rats is associated with a distinct microbiome signature.***

*(A, B) Heatmap of all gut microbiome as determined by 16S PCR for Sham*

*(n=6), Sham-Yaq-001 (n=4), BDL (n=7), BDL-Yaq-001 (n=7) groups at Family and Genus level. (C) The Venn plot of the bacteria changed with the effectiveness of the Yaq-001 at Family level. Left: The red circle indicates 21 families with significant fold change values between BDL and Sham (|log2FC|>2). The orange circle indicates 15 families with significant fold change values between BDL-Yaq-001 and BDL (|log2FC|>2). The blue circle indicates 22 families without significant fold change values between BDL-Yaq-001 and Sham (|log2FC|<=2). The overlapping part indicates 6 families with abundance changes with the treatment outcomes. Right: The red circle indicates 7 families with abundance being significantly different between BDL and Sham (Wilcoxon rank sum test, p<0.05). The orange circle indicates 1 family with abundance being significantly different between BDL-Yaq-001 and BDL (Wilcoxon rank sum test, p<0.05). The blue circle indicates 32 families without abundance being significantly different between BDL-Yaq-001 and Sham (Wilcoxon rank sum test, p>0.05). The overlapping part indicates 1 family with abundance differed with the treatment outcomes. (D) The Venn plot of the bacteria changes with the effect of the Yaq-001 at Genus level. Left: The red circle indicates 68 genera with significant fold change values between BDL and Sham (|log2FC|>2). The orange circle indicates 56 genera with significant fold change values between BDL-Yaq-001 and BDL (|log2FC|>2). The blue circle indicates 59 genera without significant fold change value between BDL-Yaq-001 and Sham (|log2FC|<=2). The overlapping part indicates 19 genera with abundance changed with the treatment outcomes. Right: The red circle indicates 17 genera with abundance being significantly different between BDL and Sham (Wilcoxon rank sum test, p<0.05). The orange circle indicates 8 genera with abundance being significantly different between BDL-Yaq-001 and BDL (Wilcoxon rank sum test, p<0.05). The blue circle indicates 114 genera without abundance being significantly different between BDL-Yaq-001 and Sham (Wilcoxon rank sum test, p>0.05). The overlapping part indicates 1 genus with abundance differed with the treatment outcomes. (E) Beta diversity*

*analysis of 16S microbiome data between four groups. The PCoA plot shows the clustering of samples from each group based on Bray-Curtis distance of 16S microbiome data. The percentage of variation explained by PCoA1 is 45.76%, and 12.95% for PCoA2. Together, PCoA1 and PCoA2 capture a total of 58.71% of the variation in the microbiome data. PERMANOVA analysis revealed a significant difference in beta diversity between groups (R2 = 0.32, p = 0.001). The BDL group exhibited the most distinct microbial community composition, while the Sham group had the most similar microbial community composition to the Sham-Yaq-001 group. Treatment with Yaq-001 appeared to moderately restore the beta diversity in the BDL group especially in PCoA2 axis. (F) Alpha diversity measures of 16S microbiome data of four groups. The box plot shows the distribution of alpha diversity values (Observed, Chao1, Shannon and Simpson indexes) for each group: BDL, BDL-Yaq-001, Sham, and Sham-Yaq-001. The middle black line in each box shows the median value of measures, the bottom and top lines of the box indicate the first and third quartiles, and the box whiskers represent the range of alpha diversity measures. The Simpson values were significantly different between groups (p<0.05, KW-test), with the highest Simpson value observed in the BDL-Yaq-001 group and the lowest diversity observed in the Sham group. Treatment with Yaq-001 appeared to increase alpha diversity in both BDL and Sham groups across four alpha diversity measures. Overall, these results suggest that bile duct ligation is associated with a reduction in microbial diversity in Observed and Chao1 indexes though the p values are not significant as the limited number of rats in this study, which can be partially restored via Yaq-001 treatment.*

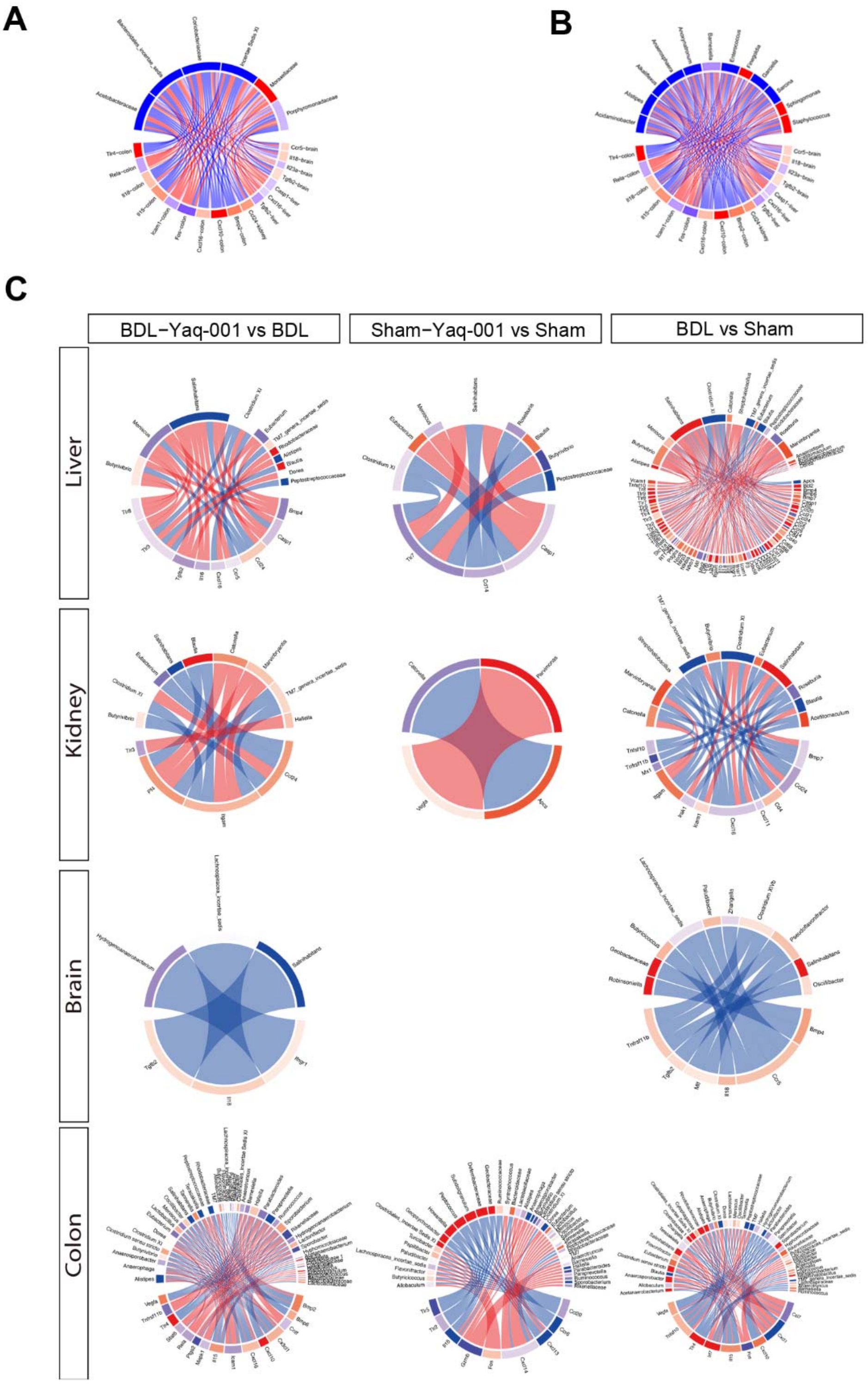


***Fig.S6. Circos plots illustrating the correlation between gene expression profiles of different tissues and microbiome abundance data.***

*(A, B) The correlations between all significantly changed DEGs and gut microbiome at family/genus. Nodes represent either genes (lower semi-circular*

*part) or bacteria (upper semi-circular part) at the family and genus level. The nodes are colored based on the log-fold change for the differential gene expression and bacteria abundance differences. The red nodes indicate an increase and blue nodes indicate a decrease. Edges represent the correlation coefficients calculated between genes and microbial genus or family with red indicating a positive correlation and blue a negative correlation. Correlation coefficients greater or equal to 0.4 were plotted in plot A (Spearman's coefficient >= 0.4), and B shows all correlations. (C) Spearman's rank correlation coefficients were first calculated between the microbiome abundance data and the respective tissue gene expression datasets and plotted for each differential gene expression experiment to show the changes for microbes and genes for each comparison. Nodes represent either genes or microbial genus/family and are coloured based on the log-fold change for the differential gene expression experiment with red indicating an increase and blue a decrease. Edges represent the correlation coefficient calculated between genes and microbial genus or family with red indicating a positive correlation and blue a negative correlation. Correlation coefficients greater or equal to 0.8 were plotted.*

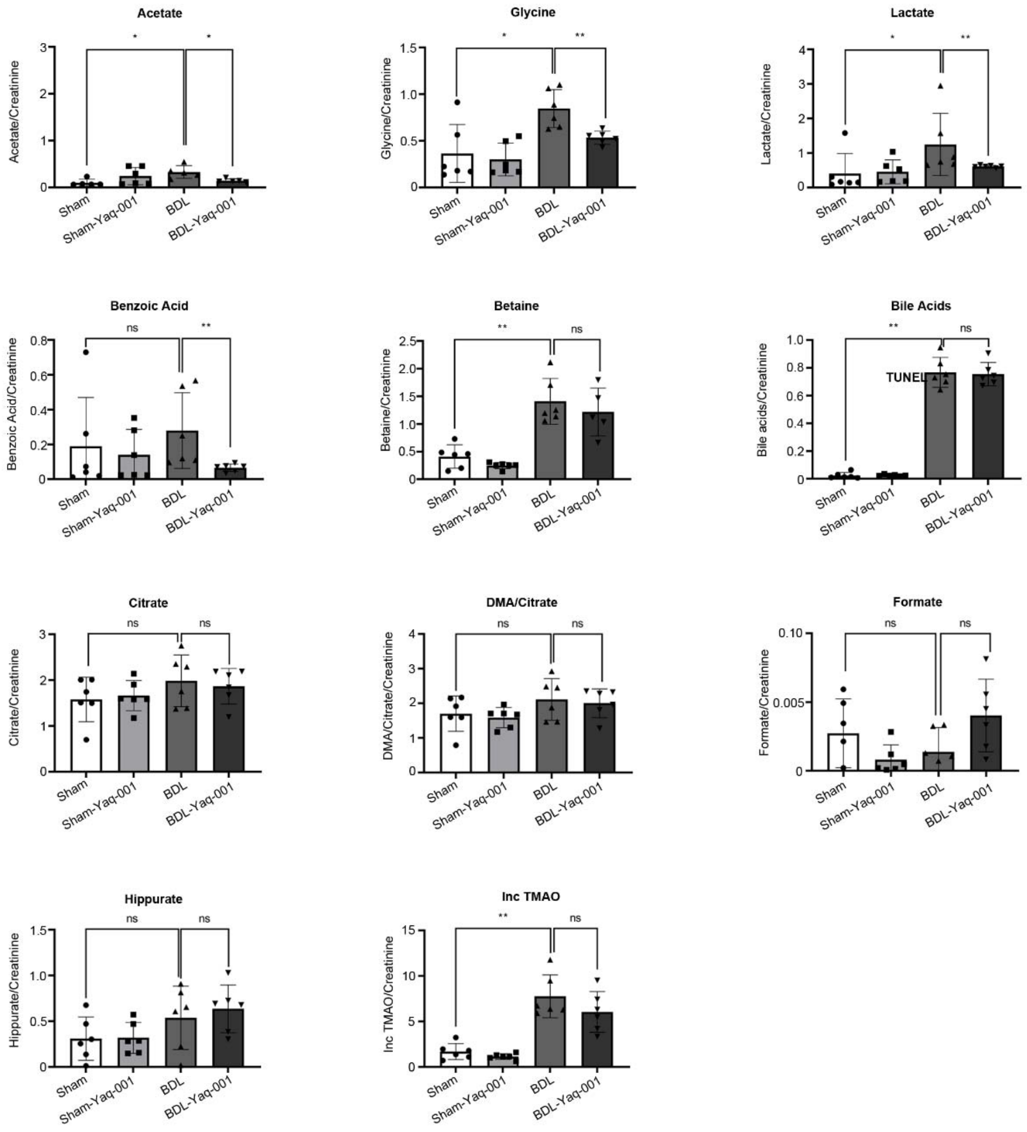


***Fig.S7. Urinary [1]NMR analysis.***

*Relative urinary concentrations of metabolites in Sham, Sham-Yaq-001, BDL and BDL-Yaq-001 (n=6/group). Significant difference of Acetate/Creatinine, Glycine/Creatinine, Lactate/Creatinine, Betaine/Creatinine, Inc TMAO/Creatinine and Bile acids/Creatinine was observed in BDL compared to Sham. Significant difference of Acetate/Creatinine, Glycine/Creatinine and Lactate/Creatinine was also observed in BDL-Yaq-001 compared to BDL.*

**Supplementary Tables**

***Table S1.All raw data for BDL and ACLF models.***

| XR code | Group | ALT | ALP | TBIL | Bile Acid | Alb | Creatinine | Wt(0) | Wt(4) | Urea | Tunel |
|---|---|---|---|---|---|---|---|---|---|---|---|
| 2246 | Sham | 12 | 25 | 0.3 | 9.7 | 15.8739 | 30 | 322 | 472 | 1.7 | 10256 |
| 2247 | Sham | 43 | 157 | 1.3 | 33.5 | 196.302 | 34 | 304 | 443 | 5.8 | 11730 |
| 2252 | Sham | 38 | 110 | 2 | 30.5 | 193.066 | 32 | 317 | 445 | 4.8 | 9420 |
| 2091 | Sham | | | | | | | 329 | 462 | | 12422 |
| 2092 | Sham | | | | | | | 343 | 471 | | 8739 |
| 2093 | Sham | | | | | | | 322 | 452 | | 7924 |
| 1582 | Sham | 46.9 | 116.7 | 0.6 | 35.25 | | 28.1 | 253 | 436 | 4.88 | 8472 |
| 1612 | Sham | 45.3 | 418.0 | < 0.0 | 35.19 | | 21.5 | 386 | 471 | 4.12 | 11343 |
| 1613 | Sham | 41.2 | 94.1 | < 0.0 | 33.04 | | 27.6 | 384 | 503 | 4.59 | 9748 |
| 1614 | Sham | 50.0 | 135.5 | 0.1 | 37.35 | | 27.2 | 385 | 475 | 5.20 | 8922 |
| 1631 | Sham | 42.9 | 114 | 0.8 | 38.12 | | | 354 | 471 | 0.09 | 7923 |
| 1632 | Sham | 50 | 86.7 | 2 | 36.1 | | | 362 | 467 | 3.08 | 13454 |
| 1633 | Sham | 45.3 | 51.7 | 1.3 | 32.93 | | | 351 | 467 | 4.08 | 12124 |
| 1634 | Sham | 59.8 | 89.5 | 2.1 | 33.84 | | | 354 | 473 | 6.23 | 11283 |
| 1714 | Sham | | | | | | | 343 | 484 | | 8924 |
| 1715 | Sham | | | | | | | 342 | 512 | | |

| | | | | | | | | | | |
|---|---|---|---|---|---|---|---|---|---|---|
| 1721 | Sham | | | | | | 22.6 | 339 | 487 | 5.07 |
| 1779 | Sham | | | | | | 32 | 405 | 417 | 5.7 |
| 1780 | Sham | | | | | | 30 | 361 | 456 | 5.5 |
| 1781 | Sham | | | | | | 39 | 390 | 501 | 4.5 |
| 1782 | Sham | | | | | | 27 | 368 | 516 | 4.3 |
| 1783 | Sham | | | | | | 36 | 371 | 480 | 4.4 |
| 2253 | Sham | 45 | 114 | 1.9 | 34.3 | 20.401 | 31 | 360 | 504 | 5.9 |
| 2309 | Sham | 53 | | 2 | 28.7 | | 27 | 359 | 432 | 5.7 |
| 2310 | Sham | 71 | 104 | 1 | 32.1 | | 28 | 378 | 452 | 6.6 |
| 2311 | Sham | 50 | 151 | 1.8 | 30.6 | | 30 | 365 | 485 | 5.5 |
| 2312 | Sham | 49 | 84 | 1.4 | 32.6 | | 30 | 352 | 468 | 7 |
| 2313 | Sham | 55 | 92 | 2 | 31.4 | | 27 | 379 | 499 | 5.2 |
| 2097 | Sham | | | | | | | 330 | 455 | |
| 2098 | Sham | | | | | | | 384 | 537 | |
| 2099 | Sham | | | | | | | 336 | 483 | |
| 3319 | Sham | | | | | | | | | 3.67 |
| 3320 | Sham | | | | | | | | | 4.71 |
| 3334 | Sham | | | | | | | | | 4 |
| 3335 | Sham | | | | | | | | | 3.83 |

| | | | | | | | | | | | |
|---|---|---|---|---|---|---|---|---|---|---|---|
| 3336 | Sham | | | | | | | | | 4.71 | |
| 2254 | Sham-Yaq-001 | 35 | | | | 31.708 | | 310 | 472 | | 13423 |
| 2255 | Sham-Yaq-001 | | | | | 11.9092 | 32 | 315 | 436 | | 12459 |
| 2256 | Sham-Yaq-001 | 28 | 165 | 1.5 | 34.1 | 35.8234 | 30 | 319 | 460 | 4 | 11732 |
| 2257 | Sham-Yaq-001 | 24 | 93 | 1 | 30.3 | 14.3252 | 31 | 315 | 445 | 4.1 | 9789 |
| 2335 | Sham-Yaq-001 | 53 | 113 | 2 | 33.6 | | 25 | 405 | 513 | 6.3 | 10210 |
| 2336 | Sham-Yaq-001 | 41 | 119 | 1 | 31.3 | 4.411 | 24 | 408 | 515 | 5.3 | 14715 |
| 2337 | Sham-Yaq-001 | 31 | 106 | 1 | 26 | 4.1664 | 21 | 392 | 489 | 4.4 | 7543 |
| 2338 | Sham-Yaq-001 | 49 | 59 | 1.2 | 21.9 | 19.9726 | 22 | 385 | 478 | 5.4 | 8973 |
| 2339 | Sham-Yaq-001 | 42 | 127 | 1 | 29.1 | | 28 | 387 | 469 | 5.1 | 9782 |
| 1607 | Sham-Yaq-001 | 33.9 | 114.8 | 0.4 | 35.56 | | 29.4 | 359 | 418 | 4.27 | 10747 |
| 1608 | Sham-Yaq-001 | 37.4 | 76.6 | 0.4 | 34.56 | | 32.0 | 342 | 475 | 5.82 | 11793 |
| 1609 | Sham-Yaq-001 | 33.4 | 130.0 | < 0.0 | 34.00 | | 29.5 | 383 | 445 | 5.34 | 9784 |
| 1610 | Sham-Yaq-001 | 30.5 | 95.1 | 0.4 | 33.30 | | 25.0 | 328 | 453 | 4.62 | 8792 |
| 1643 | Sham-Yaq-001 | | | | | | | 403 | 504 | | 8243 |
| 1644 | Sham-Yaq-001 | | | | | | | 386 | 556 | | 7333 |
| 1645 | Sham-Yaq-001 | | | | | | | 358 | 474 | | |
| 1646 | Sham-Yaq-001 | | | | | | | 368 | 485 | | |
| 1653 | Sham-Yaq-001 | 32.4 | | 1.7 | | | | 372 | 475 | 3.02 | |

| | | | | | | | | | | |
|---|---|---|---|---|---|---|---|---|---|---|
| 1654 | Sham-Yaq-001 | 33.5 | | 1.5 | | | | 370 | 485 | |
| 1794 | Sham-Yaq-001 | | | | | | 37 | 436 | 514 | 5.8 |
| 1795 | Sham-Yaq-001 | | | | | | 35 | 427 | 510 | 4.9 |
| 1796 | Sham-Yaq-001 | | | | | | 34 | 456 | 511 | 4.5 |
| 1797 | Sham-Yaq-001 | | | | | | 41 | 429 | 569 | 5.6 |
| 1798 | Sham-Yaq-001 | | | | | | 38 | 397 | 498 | 5.4 |
| 3322 | Sham-Yaq-001 | | | | | | | | | 3.67 |
| 3323 | Sham-Yaq-001 | | | | | | | | | 4.52 |
| 3324 | Sham-Yaq-001 | | | | | | | | | 0.14 |
| 3328 | Sham-Yaq-001 | | | | | | | | | 3.85 |
| 3329 | Sham-Yaq-001 | | | | | | | | | 4.65 |
| 3330 | Sham-Yaq-001 | | | | | | | | | 3.66 |
| 2248 | BDL | | | | | 38.667 | 39 | 320 | 420 | 6.8 |
| 2249 | BDL | 84 | 212 | 128.6 | 26.7 | 49.518 | 33 | 317 | 416 | 5.9 |
| 2250 | BDL | 66 | 160 | 141.9 | 26.4 | 33.113 | 40 | 322 | 424 | 4.1 |
| 2251 | BDL | 61 | 165 | 173.7 | 27.1 | 38.25 | 37 | 313 | 394 | 6 |
| 2089 | BDL | | | | | | | 351 | 473 | |
| 2090 | BDL | | | | | | | 353 | 428 | |
| 2088 | BDL | | | | | | | 347 | 448 | |

| | | | | | | | | | | | |
|---|---|---|---|---|---|---|---|---|---|---|---|
| 1584 | BDL | 83.3 | 325.9 | 98.2 | 19.90 | | 33.0 | 309 | 396 | 3.72 | 42393 |
| 1586 | BDL | 61.4 | 196.8 | 141.1 | 26.27 | | 32.2 | 281 | 452 | 5.84 | 57892 |
| 1592 | BDL | 82.8 | 399.9 | 93.0 | 13.08 | | 26.3 | 303 | 386 | 4.57 | 32728 |
| 1594 | BDL | 55.8 | 256.9 | 110.7 | 24.86 | | 29.0 | 295 | 395 | 4.59 | 34888 |
| 1619 | BDL | 88 | 288 | 121.3 | 24.4 | | | 339 | 425 | | 60723 |
| 1621 | BDL | 102 | 283 | 79.3 | 16.2 | | | 356 | 429 | | 59872 |
| 1622 | BDL | | 195 | 111.3 | 22.3 | | | 351 | 447 | | 64324 |
| 1625 | BDL | 106 | 254 | 143 | 13.2 | | | 342 | 443 | | 67822 |
| 1626 | BDL | 82 | 147 | 188.7 | 23.9 | | | 340 | 413 | 5 | 49322 |
| 1723 | BDL | | | | | | 23.7 | | | 5 | 52843 |
| 1775 | BDL | 103 | 95 | 178.8 | 32.5 | 84.902 | 32 | 351 | 402 | 5.9 | 37420 |
| 1777 | BDL | 109 | 254 | 139.7 | 27 | 74.784 | 38 | 335 | 379 | 5.7 | 42534 |
| 1778 | BDL | 105 | 232 | 148.9 | 25.7 | | 28 | 333 | 390 | 4.8 | 69423 |
| 1799 | BDL | 69 | 216 | 158.3 | 20 | | 26 | 409 | 475 | 5 | 52432 |
| 1800 | BDL | | | | | | 36 | 423 | 455 | 4.6 | 49320 |
| 1801 | BDL | | | | | | 33 | 391 | 450 | 5.5 | |
| 1802 | BDL | | | | | | 33 | 412 | 422 | 6 | |
| 1803 | BDL | | | | | | 39 | 325 | 375 | 5.8 | |
| 2304 | BDL | | | | | | | | *466* | 6.9 | |

| | | | | | | | | | | |
|---|---|---|---|---|---|---|---|---|---|---|
| 2305 | BDL | | | | | | 33 | *369* | *453* | 6.6 |
| 2306 | BDL | 60 | 342 | 152.2 | 21.5 | | 23 | 375 | 436 | 4.3 |
| 2307 | BDL | | | | | | | | 435 | 6.4 |
| 2308 | BDL | 62 | 250 | 133 | 28.2 | | 32 | 362 | 450 | 6.2 |
| 1774 | BDL | | | | | | | | 479 | 5.1 |
| 1776 | BDL | | | | | | | 270 | 334 | 6.3 |
| 3316 | BDL | | | | | | | | | 3.87 |
| 3317 | BDL | | | | | | | | | 3.23 |
| 3318 | BDL | | | | | | | | | 4.6 |
| 3337 | BDL | | | | | | | | | 4.34 |
| 3339 | BDL | | | | | | | | | 7.72 |
| 2330 | BDL-Yaq-001 | 88 | 245 | 99.2 | 30.8 | | 32 | 332 | 365 | 3.5 |
| 2331 | BDL-Yaq-001 | 80 | 226 | 100.5 | 25 | 34.2716 | 32 | 348 | 407 | 3.5 |
| 2332 | BDL-Yaq-001 | | | | | | | | *436* | 3.4 |
| 2333 | BDL-Yaq-001 | 88 | 240 | 132 | 27.3 | | 30 | 356 | 411 | 6.2 |
| 2334 | BDL-Yaq-001 | 61 | 173 | 128.9 | 26.1 | 30.632 | 28 | 461 | 507 | 4.6 |
| 1602 | BDL-Yaq-001 | | | | | | | 440 | 543 | 5.19 |
| 1642 | BDL-Yaq-001 | | | | | | | | *400* | 5.9 |
| 1657 | BDL-Yaq-001 | | | | | | | | 485 | |

| | | | | | | | | | | |
|---|---|---|---|---|---|---|---|---|---|---|
| 1658 | BDL-Yaq-001 | | | | | | | | 522 | |
| 3325 | BDL-Yaq-001 | | | | | | | | | 4.91 |
| 3326 | BDL-Yaq-001 | | | | | | | | | 4.04 |
| 3327 | BDL-Yaq-001 | | | | | | | | | 4.52 |
| 3331 | BDL-Yaq-001 | | | | | | | | | 4.33 |
| 3332 | BDL-Yaq-001 | | | | | | | | | 4.64 |
| 3333 | BDL-Yaq-001 | | | | | | | | | 5.64 |
| 1599 | BDL-Yaq-001 | 54.0 | 271.0 | 117.0 | 21.83 | | 29.9 | 305 | 448 | 4.87 |
| 1600 | BDL-Yaq-001 | 28.0 | 211.5 | 123.8 | 25.10 | | 29.8 | 295 | 385 | 6.10 |
| 1601 | BDL-Yaq-001 | 54.6 | 225.4 | 99.8 | 24.27 | | 31.8 | 287 | 395 | 5.34 |
| 1615 | BDL-Yaq-001 | 47.7 | 262.5 | 138.4 | 26.94 | | 18.8 | 358 | 398 | 3.47 |
| 1616 | BDL-Yaq-001 | 81.2 | 260.5 | 102.1 | 25.68 | | 24.1 | 372 | 413 | 5.50 |
| 1635 | BDL-Yaq-001 | 48 | 219 | 133.7 | 20.4 | | | 358 | 452 | 3.5 |
| 1636 | BDL-Yaq-001 | 74 | 238 | 141.2 | 24.6 | | | 381 | 482 | 3.9 |
| 1637 | BDL-Yaq-001 | 69 | 197 | 105.5 | 20.8 | | | 353 | 440 | 5.3 |
| 1638 | BDL-Yaq-001 | 56 | 192 | 135.1 | 26.4 | | | 378 | 471 | 5.9 |
| 1655 | BDL-Yaq-001 | 67 | 259 | 130 | 89 | | | 365 | 489 | 0 |
| 1716 | BDL-Yaq-001 | | | | | | | 334 | 415 | |
| 1717 | BDL-Yaq-001 | | | | | | | 357 | 456 | |

| | | | | | | | | | | | |
|---|---|---|---|---|---|---|---|---|---|---|---|
| 1718 | BDL-Yaq-001 | | | | | | | 348 | 443 | | |
| 1784 | BDL-Yaq-001 | | | | | | | 416 | 487 | | |
| 1785 | BDL-Yaq-001 | 67 | 190 | 122.1 | 22.9 | | 27 | 412 | 410 | 4.6 | 37832 |
| 1786 | BDL-Yaq-001 | 99 | 214 | 94.9 | 29 | | | 408 | 461 | 5.8 | 32420 |
| 1787 | BDL-Yaq-001 | 132 | 147 | 169.1 | 26.6 | | 29 | 409 | 440 | 4.3 | 25457 |
| 1788 | BDL-Yaq-001 | 77 | 233 | 152.3 | 32 | | | 448 | 477 | 6 | 27822 |
| 1789 | BDL-Yaq-001 | 50 | 181 | 134.4 | 32.1 | | | 389 | 438 | 5 | 52893 |
| 1790 | BDL-Yaq-001 | 34 | 173 | 172.2 | 29.1 | | 28 | 415 | 510 | 4.9 | 52434 |
| 1791 | BDL-Yaq-001 | 38 | 138 | 125.6 | 27.6 | | 36 | 392 | 431 | 5.2 | 58723 |
| 1793 | BDL-Yaq-001 | 68 | 185 | 123.6 | 26.6 | | | 383 | 438 | 4.9 | 47433 |
| 2242 | BDL-Yaq-001 | 53 | 202 | 132.2 | 26.8 | 35.028 | 29 | 302 | 387 | 4.9 | 38433 |
| 2243 | BDL-Yaq-001 | 33 | 193 | 122.5 | 21.4 | 54.552 | 29 | 339 | 435 | 5.6 | 35303 |
| 2244 | BDL-Yaq-001 | 64 | 292 | 136.8 | 25 | 42.238 | 31 | 315 | 405 | 4.2 | 32777 |
| 2245 | BDL-Yaq-001 | 46 | 249 | 159.8 | 31 | 49.625 | 32 | 304 | 402 | 5.4 | 39433 |
| 2095 | BDL-Yaq-001 | | | | | | | 337 | 438 | | 37823 |
| 2096 | BDL-Yaq-001 | | | | | | | 340 | 413 | | 49843 |
| 2102 | BDL-Yaq-001 | | | | | | | 369 | 441 | | 58787 |
| 1579 | Sham-LPS | | | | | | | 255 | 396 | | |
| 1580 | Sham-LPS | 58.6 | 149.3 | < 0.0 | 35.52 | | 20.7 | 239 | 371 | 3.28 | |

| | | | | | | | | | |
|---|---|---|---|---|---|---|---|---|---|
| 1581 | Sham-LPS | 42.5 | 116.4 | 0.1 | 33.36 | 24.2 | 254 | 436 | 4.13 |
| 1611 | Sham-LPS | 54.2 | 85.1 | < 0.0 | 33.36 | 19.8 | 332 | 397 | 4.57 |
| 1627 | Sham-LPS | 42 | 70.4 | 1.4 | 33.36 | | 359 | 456 | 4.08 |
| 1628 | Sham-LPS | 102.4 | 71 | 1.8 | 33.84 | | 348 | 458 | 6.62 |
| 1629 | Sham-LPS | 116.6 | 91.8 | 1.5 | 33 | | 376 | 528 | 4.71 |
| 1630 | Sham-LPS | 73.3 | 61.8 | 2 | 33 | | 355 | 487 | 5.51 |
| 1722 | Sham-LPS | | | | | 24.7 | 331 | 487 | 5.39 |
| 1603 | Sham-LPS-Yaq-001 | 39.2 | 77.8 | < 0.0 | 30.00 | 25.7 | 336 | 464 | 5.09 |
| 1604 | Sham-LPS-Yaq-001 | 47.9 | | | 30.29 | | 374 | 438 | |
| 1605 | Sham-LPS-Yaq-001 | 50.3 | 101.8 | 3.7 | 29.88 | 44.0 | 358 | 481 | 5.34 |
| 1606 | Sham-LPS-Yaq-001 | 50.3 | 146.7 | 0.6 | 34.01 | 28.9 | 378 | 446 | 6.22 |
| 1647 | Sham-LPS-Yaq-001 | | | | | | 368 | 481 | |
| 1648 | Sham-LPS-Yaq-001 | | | | | | 398 | 521 | |
| 1649 | Sham-LPS-Yaq-001 | | | | | | 358 | 450 | |
| 1650 | Sham-LPS-Yaq-001 | | | | | | 392 | 560 | |
| 1651 | Sham-LPS-Yaq-001 | 43.9 | 69.5 | 0.7 | | | 408 | 549 | 4.14 |
| 1652 | Sham-LPS-Yaq-001 | | | | | | 354 | 481 | |
| 1583 | BDL-LPS | 59.4 | 254.2 | 130.1 | 30.93 | 34.3 | 271 | 412 | 5.52 |
| 1587 | BDL-LPS | 123.8 | 458.4 | 79.0 | 13.99 | 29.7 | 277 | 372 | 5.79 |

| | | | | | | | | | |
|---|---|---|---|---|---|---|---|---|---|
| 1588 | BDL-LPS | | 1051.1 | 151.9 | 17.48 | 59.4 | 284 | 362 | 7.63 |
| 1589 | BDL-LPS | 104.5 | 236.5 | 148.3 | 31.21 | 22.4 | 251 | 359 | 4.01 |
| 1590 | BDL-LPS | 104.8 | 266.9 | 140.9 | 27.69 | 25.4 | 286 | 387 | 5.80 |
| 1591 | BDL-LPS | 109.5 | 191.7 | 123.1 | 27.06 | 42.4 | 276 | 381 | 5.97 |
| 1712 | BDL-LPS | | | | | | 333 | 441 | |
| 1713 | BDL-LPS | | | | | | 345 | 478 | |
| 1724 | BDL-LPS | | | | | 38.6 | 382 | 467 | 9.64 |
| 1725 | BDL-LPS | | | | | | 369 | 464 | |
| 1726 | BDL-LPS | | | | | | 387 | 472 | |
| 2319 | BDL-LPS | 107 | 175 | 123.8 | 24 | 60 | 381 | 439 | 8.1 |
| 2320 | BDL-LPS | 124 | 287 | 136.9 | 25.5 | 59 | 412 | 473 | 7.8 |
| 2321 | BDL-LPS | 94 | 284 | 123.6 | 22.3 | 44 | 392 | 419 | 7.4 |
| 2322 | BDL-LPS | 60 | 234 | 116.9 | 26.6 | 38 | 398 | 444 | 7.8 |
| 2323 | BDL-LPS | 85 | 226 | 132 | 21.5 | 44 | 402 | 462 | 6.1 |
| 1595 | BDL-LPS-Yaq-001 | 68.5 | 231.6 | 140.5 | 30.87 | 26.3 | 302 | 430 | 5.22 |
| 1596 | BDL-LPS-Yaq-001 | 57.3 | 199.1 | 123.2 | 27.06 | 30.9 | 309 | 390 | 4.28 |
| 1597 | BDL-LPS-Yaq-001 | 46.1 | 234.8 | 139.0 | 25.37 | 33.3 | 304 | 385 | 6.47 |
| 1598 | BDL-LPS-Yaq-001 | 50.8 | 263.3 | 152.9 | 29.72 | 25.3 | 322 | 432 | 5.00 |
| 1617 | BDL-LPS-Yaq-001 | 84.9 | 223.2 | 87.9 | 27.71 | 28.3 | 324 | 362 | 6.41 |

| | | | | | | | | | |
|---|---|---|---|---|---|---|---|---|---|
| 1618 | BDL-LPS-Yaq-001 | 66.4 | 204.9 | 90.9 | 25.79 | 27.2 | 334 | 408 | 4.87 |
| 1639 | BDL-LPS-Yaq-001 | | | | | | 365 | 460 | |
| 1640 | BDL-LPS-Yaq-001 | 67 | 200 | 124.4 | 21.3 | | 400 | 486 | 10.8 |
| 1641 | BDL-LPS-Yaq-001 | 59 | 163 | 115.6 | 22.4 | | 368 | 458 | 8.1 |
| 1656 | BDL-LPS-Yaq-001 | 86 | 276 | 14.8 | 32 | | 378 | 491 | |
| 1719 | BDL-LPS-Yaq-001 | | | | | | 352 | 442 | |
| 1720 | BDL-LPS-Yaq-001 | | | | | | 358 | 497 | |

***Raw data for biological analysis in Experiment 1, Experiment 2 and Experiment 3.***

*ALT: Plasma alanine transaminase; ALP: Plasma alkaline phosphatase; TBIL: total bilirubin; Alb: Albumin; Wt (0): the weight in the first day of the experiment; Wt (4): the weight at the end of the experiment; Urea; plasma urea concentrations; Tunel: Terminal deoxynucleotidyl transferase (TdT) dUTP Nick-End Labeling assay*

| XR code | Group | PP | MAP | CPA | BW | NH3A | NH3PV | A-E | PV-E | A-DNA | PV-DNA |
|---|---|---|---|---|---|---|---|---|---|---|---|
| 2246 | Sham | 4.74 | | | | | | | | | |
| 2247 | Sham | 6.507 | | | | | | | | | |
| 2252 | Sham | 2.537 | | | | | | | | | |
| 2091 | Sham | | | | | | | | | | |
| 2092 | Sham | | | | | | | | | | |
| 2093 | Sham | | | | | | | | | | |
| 1582 | Sham | 6.15 | 128.71 | 0.027 | | | | | | | |
| 1612 | Sham | 5.17 | 137.31 | 0.02 | | | | | | | |
| 1613 | Sham | 5.84 | 110.42 | | | | | | | | |
| 1614 | Sham | 5.38 | 115.36 | 0.02 | | | | | | | |
| 1631 | Sham | 4.05 | 121 | | | 36.2 | | | | | |
| 1632 | Sham | 2.37 | 123 | 0.02 | | | | | | | |
| 1633 | Sham | 4.88 | 104 | 0.004 | | | | | | | |
| 1634 | Sham | 6.7 | 105.2 | 0.005 | | | | | | | |
| 1714 | Sham | | | | | | | | | | |
| 1715 | Sham | | | | | | | | | | |

| | | | | | | | | | |
|---|---|---|---|---|---|---|---|---|---|
| 1721 | Sham | | | 54 | 132 | 0.1 | 0.2 | NEG | NEG |
| 1779 | Sham | | | 59.3 | 161 | 0.1 | 0.3 | NEG | NEG |
| 1780 | Sham | | | 94.4 | 143 | 0.2 | 0.3 | NEG | NEG |
| 1781 | Sham | | | 61 | 132 | 0.1 | 0.2 | NEG | NEG |
| 1782 | Sham | | | 55 | 211 | 0.1 | 0.1 | NEG | NEG |
| 1783 | Sham | | | 57 | 111 | 0 | 0 | NEG | NEG |
| 2253 | Sham | 6.687 | | | | | | | |
| 2309 | Sham | 5.77 | 104.4 | | | | | | |
| 2310 | Sham | 8.5 | 108.8 | | | | | | |
| 2311 | Sham | 7.08 | 102 | | | | | | |
| 2312 | Sham | 7.71 | 89.43 | | | | | | |
| 2313 | Sham | 7.2 | 116.1 | | | | | | |
| 2097 | Sham | | | | | | | | |
| 2098 | Sham | | | | | | | | |
| 2099 | Sham | | | | | | | | |
| 3319 | Sham | | | | | | | | |
| 3320 | Sham | | | | | | | | |

| | | | | |
|---|---|---|---|---|
| 3334 | Sham | | | |
| 3335 | Sham | | | |
| 3336 | Sham | | | |
| 2254 | Sham-Yaq-001 | 7.773 | | |
| 2255 | Sham-Yaq-001 | 7.627 | | |
| 2256 | Sham-Yaq-001 | 6.923 | | |
| 2257 | Sham-Yaq-001 | 5.73 | | |
| 2335 | Sham-Yaq-001 | 4.43 | 112.8 | |
| 2336 | Sham-Yaq-001 | 4.77 | 111.4 | |
| 2337 | Sham-Yaq-001 | 7.24 | 113.5 | |
| 2338 | Sham-Yaq-001 | 8.48 | 118.7 | |
| 2339 | Sham-Yaq-001 | 6.37 | 103.9 | |
| 1607 | Sham-Yaq-001 | 5.69 | 121.85 | 0.007 |
| 1608 | Sham-Yaq-001 | 6.13 | 116.66 | 0.013 |
| 1609 | Sham-Yaq-001 | 5.93 | 130 | 0.015 |
| 1610 | Sham-Yaq-001 | 4.38 | 107 | 0.009 |
| 1643 | Sham-Yaq-001 | 7.5 | | 0.011 |

| | | | | | | | | | | |
|---|---|---|---|---|---|---|---|---|---|---|
| 1644 | Sham-Yaq-001 | 4.15 | | 0.008 | | | | | | |
| 1645 | Sham-Yaq-001 | 6.8 | | 0.01 | | | | | | |
| 1646 | Sham-Yaq-001 | 4.5 | | 0.009 | | | | | | |
| 1653 | Sham-Yaq-001 | 7.7 | 103 | 0.01 | | | | | | |
| 1654 | Sham-Yaq-001 | 6.77 | 129 | 0.003 | | | | | | |
| 1794 | Sham-Yaq-001 | | | | 31 | 98 | 0.05 | 0.1 | NEG | NEG |
| 1795 | Sham-Yaq-001 | | | | 51 | 101 | 0 | 0.1 | NEG | NEG |
| 1796 | Sham-Yaq-001 | | | | 45 | 52 | 0 | 0.1 | NEG | NEG |
| 1797 | Sham-Yaq-001 | | | | 54 | 61 | 0.1 | 0.2 | NEG | NEG |
| 1798 | Sham-Yaq-001 | | | | 61 | 73 | 0.1 | 0.18 | NEG | NEG |
| 3322 | Sham-Yaq-001 | | | | | | | | | |
| 3323 | Sham-Yaq-001 | | | | | | | | | |
| 3324 | Sham-Yaq-001 | | | | | | | | | |
| 3328 | Sham-Yaq-001 | | | | | | | | | |
| 3329 | Sham-Yaq-001 | | | | | | | | | |
| 3330 | Sham-Yaq-001 | | | | | | | | | |
| 2248 | BDL | | | | 245.4 | 532.6 | 0.5 | 0.62 | NEG | POS |

| | | | | | | | | | | |
|---|---|---|---|---|---|---|---|---|---|---|
| 2249 | BDL | 9.937 | | | 160.7 | 280.4 | 0.65 | 0.98 | POS | POS |
| 2250 | BDL | 12.09 | | | 214 | 520.5 | 0.61 | 1.3 | POS | POS |
| 2251 | BDL | 10.7 | | | 354 | 441.7 | 0.73 | 1.8 | NEG | NEG |
| 2089 | BDL | | | | 231 | 341 | 1.1 | 1.5 | NEG | NEG |
| 2090 | BDL | | | | 198 | 332 | 0.83 | 1.2 | NEG | POS |
| 2088 | BDL | | | | 211 | 313 | 1 | 1.3 | NEG | NEG |
| 1584 | BDL | 10.83 | 74 | 0.092 | | | | | | |
| 1586 | BDL | 12.75 | 104.5 | 0.262 | | | | | | |
| 1592 | BDL | | 101 | 0.491 | | | | | | |
| 1594 | BDL | | 84.66 | 0.284 | | | | | | |
| 1619 | BDL | 15.2 | 97.1 | 0.344 | 172.8 | | 0.3 | 0.8 | POS | POS |
| 1621 | BDL | 11.3 | 48 | 0.093 | 281.3 | | 0.7 | 1.2 | POS | POS |
| 1622 | BDL | | 89 | 0.118 | 116.4 | | 1.2 | 1.4 | NEG | NEG |
| 1625 | BDL | 9.4 | 55.7 | 0.11 | | | 0.9 | 1.3 | NEG | POS |
| 1626 | BDL | 14.7 | 67.5 | | | | 1.4 | 1.8 | NEG | NEG |
| 1723 | BDL | | | | | | | | | |
| 1775 | BDL | 12.88 | | | 139 | 265.9 | | | | |

| | | | | | |
|---|---|---|---|---|---|
| 1777 | BDL | 13.44 | | 227 | |
| 1778 | BDL | 13.25 | | 164.9 | 204.1 |
| 1799 | BDL | | | 185.8 | 345.6 |
| 1800 | BDL | | | 224.3 | 429.7 |
| 1801 | BDL | | | 148.7 | |
| 1802 | BDL | | | 149.7 | 213.1 |
| 1803 | BDL | | | 170 | 401.8 |
| 2304 | BDL | | | | |
| 2305 | BDL | | | | |
| 2306 | BDL | 12.8 | 78.77 | | |
| 2307 | BDL | | | | |
| 2308 | BDL | 12 | 112.2 | | |
| 1774 | BDL | | | | |
| 1776 | BDL | | | 93 | |
| 3316 | BDL | | | | |
| 3317 | BDL | | | | |
| 3318 | BDL | | | | |

| | | | | |
|---|---|---|---|---|
| 3337 | BDL | | | |
| 3339 | BDL | | | |
| 2330 | BDL-Yaq-001 | 10.8 | 96.03 | |
| 2331 | BDL-Yaq-001 | 9.23 | 94.88 | |
| 2332 | BDL-Yaq-001 | | | |
| 2333 | BDL-Yaq-001 | 11.4 | 93.25 | |
| 2334 | BDL-Yaq-001 | 10.3 | 83.49 | |
| 1602 | BDL-Yaq-001 | | | |
| 1642 | BDL-Yaq-001 | | | 76 |
| 1657 | BDL-Yaq-001 | | | |
| 1658 | BDL-Yaq-001 | | | |
| 3325 | BDL-Yaq-001 | | | |
| 3326 | BDL-Yaq-001 | | | |
| 3327 | BDL-Yaq-001 | | | |
| 3331 | BDL-Yaq-001 | | | |
| 3332 | BDL-Yaq-001 | | | |
| 3333 | BDL-Yaq-001 | | | |

| | | | | | | | | |
|---|---|---|---|---|---|---|---|---|
| 1599 | BDL-Yaq-001 | 10.21 | 99 | 0.05 | | | | |
| 1600 | BDL-Yaq-001 | 11.89 | 77.6 | 0.213 | | | | |
| 1601 | BDL-Yaq-001 | 11.76 | 88.3 | 0.335 | | | | |
| 1615 | BDL-Yaq-001 | 11.69 | 89.14 | 0.025 | | | | |
| 1616 | BDL-Yaq-001 | 12.19 | 105.69 | 0.249 | | | | |
| 1635 | BDL-Yaq-001 | 8.3 | 98.6 | 0.025 | 79.2 | 97 | NEG | POS |
| 1636 | BDL-Yaq-001 | 11.7 | 78.9 | 0.224 | 130.9 | 154 | NEG | NEG |
| 1637 | BDL-Yaq-001 | 11.2 | 73.8 | 0.036 | 147.1 | 176 | POS | NEG |
| 1638 | BDL-Yaq-001 | 13.1 | 95 | 0.086 | 141 | 182 | NEG | NEG |
| 1655 | BDL-Yaq-001 | 9.92 | 107.65 | 0.207 | 132 | 176 | NEG | NEG |
| 1716 | BDL-Yaq-001 | | | | 131 | 229.9 | POS | NEG |
| 1717 | BDL-Yaq-001 | | | | 112 | 150.1 | | |
| 1718 | BDL-Yaq-001 | | | | | | | |
| 1784 | BDL-Yaq-001 | | | | | | | |
| 1785 | BDL-Yaq-001 | 10.69 | | | 170.2 | 315.1 | | |
| 1786 | BDL-Yaq-001 | 10.96 | | | 98.7 | 378.1 | | |
| 1787 | BDL-Yaq-001 | 12.4 | | | | | | |

| | | | | | | | | | | | |
|---|---|---|---|---|---|---|---|---|---|---|---|
| 1788 | BDL-Yaq-001 | 10.33 | | | | 197.1 | | | | | |
| 1789 | BDL-Yaq-001 | 10.28 | | | | 143.9 | | | | | |
| 1790 | BDL-Yaq-001 | 11.06 | | | | 163.2 | 204.8 | | | | |
| 1791 | BDL-Yaq-001 | 11.48 | | | | 165.4 | 281.7 | | | | |
| 1793 | BDL-Yaq-001 | 10.34 | | | | 150.5 | 239.5 | | | | |
| 2242 | BDL-Yaq-001 | 5.62 | | | | | | 0.4 | 0.54 | NEG | NEG |
| 2243 | BDL-Yaq-001 | 11.1 | | | | 204.2 | 352.3 | 0.3 | *0.54* | NEG | NEG |
| 2244 | BDL-Yaq-001 | 11.68 | | | | 187 | 368.4 | 0.34 | 0.4 | NEG | NEG |
| 2245 | BDL-Yaq-001 | 13.48 | | | | 209.9 | 245 | 0.51 | 0.63 | NEG | NEG |
| 2095 | BDL-Yaq-001 | | | | | 145 | 217 | 0.42 | 0.5 | POS | POS |
| 2096 | BDL-Yaq-001 | | | | | 132 | 254 | 0.57 | 0.65 | NEG | NEG |
| 2102 | BDL-Yaq-001 | | | | | 121 | 231 | 0.4 | 0.45 | POS | POS |
| 1579 | Sham-LPS | 5.14 | | 0.006 | 79.55 | | | | | | |
| 1580 | Sham-LPS | 10.83 | 74 | 0.005 | 79.6 | | | | | | |
| 1581 | Sham-LPS | 12.75 | 104.5 | 0.01 | 78.69 | | | | | | |
| 1611 | Sham-LPS | 6.92 | 113.36 | 0.011 | 79.44 | | | | | | |
| 1627 | Sham-LPS | 11.2 | 115.7 | 0.009 | | 123 | 154 | | | | |

| | | | | | | | |
|---|---|---|---|---|---|---|---|
| 1628 | Sham-LPS | 7.3 | 88.5 | 0.005 | | 145 | 156 |
| 1629 | Sham-LPS | 7.3 | 104.5 | 0.013 | | 111 | 132 |
| 1630 | Sham-LPS | 6.4 | 107.1 | 0.002 | | 121 | 162 |
| 1722 | Sham-LPS | | | | | 103 | 154 |
| 1603 | Sham-LPS-Yaq-001 | 9.82 | 112.16 | 0.08 | 79.14 | | |
| 1604 | Sham-LPS-Yaq-001 | 5.78 | 111.27 | 0.008 | 79.16 | | |
| 1605 | Sham-LPS-Yaq-001 | 5.2 | 82.5 | 0.006 | 79.45 | | |
| 1606 | Sham-LPS-Yaq-001 | 7.61 | 112.55 | 0.028 | 79.71 | | |
| 1647 | Sham-LPS-Yaq-001 | 6.23 | | 0.013 | | 98 | 121 |
| 1648 | Sham-LPS-Yaq-001 | 9.69 | | 0.014 | | 76 | 143 |
| 1649 | Sham-LPS-Yaq-001 | 7.37 | | 0.008 | | 56 | 132 |
| 1650 | Sham-LPS-Yaq-001 | 8.1 | | 0.012 | | 104 | 112 |
| 1651 | Sham-LPS-Yaq-001 | 8.94 | 107.3 | 0.004 | | 105 | 167 |
| 1652 | Sham-LPS-Yaq-001 | 7.27 | | 0.015 | | | |
| 1583 | BDL-LPS | 17.22 | 92.5 | 0.11 | 80.47 | | |
| 1587 | BDL-LPS | 20.7 | 110 | 0.381 | 79.13 | | |
| 1588 | BDL-LPS | | | 0.149 | 81.4 | | |

| | | | | | | | |
|---|---|---|---|---|---|---|---|
| 1589 | BDL-LPS | 15.58 | 103 | 0.108 | 79.38 | | |
| 1590 | BDL-LPS | 17.47 | 93 | 0.178 | 80.26 | | |
| 1591 | BDL-LPS | 19.28 | 108.1 | 0.275 | | | |
| 1712 | BDL-LPS | | | | | 321 | 421 |
| 1713 | BDL-LPS | | | | | 333 | 433 |
| 1724 | BDL-LPS | | | | | 354 | 507.7 |
| 1725 | BDL-LPS | | | | | 288 | 369.4 |
| 1726 | BDL-LPS | | | | | 219 | |
| 2319 | BDL-LPS | 12.9 | 131.6 | | | 289 | 389 |
| 2320 | BDL-LPS | 15.2 | 111 | | | 333 | 411 |
| 2321 | BDL-LPS | 10.8 | 101.5 | | 80.1 | | |
| 2322 | BDL-LPS | 13.6 | 61.67 | | | | |
| 2323 | BDL-LPS | | 110.9 | | 80.3 | | |
| 1595 | BDL-LPS-Yaq-001 | 9.6 | 84.9 | 0.116 | 78.87 | 198 | 321 |
| 1596 | BDL-LPS-Yaq-001 | 12.07 | 82.44 | 0.149 | 80.34 | 211 | 333 |
| 1597 | BDL-LPS-Yaq-001 | 11.57 | 89.4 | 0.504 | 79.91 | 232 | 341 |
| 1598 | BDL-LPS-Yaq-001 | 12.86 | 101.42 | 0.158 | 80.02 | 199 | 243 |

| | | | | | | | |
|---|---|---|---|---|---|---|---|
| 1617 | BDL-LPS-Yaq-001 | 14 | 63.87 | 0.24 | 80.37 | 278 | 402 |
| 1618 | BDL-LPS-Yaq-001 | 14.53 | 107.77 | 0.284 | 78.58 | | |
| 1639 | BDL-LPS-Yaq-001 | | | 0.132 | 77.8 | | |
| 1640 | BDL-LPS-Yaq-001 | 4.75 | 85.4 | | 78.2 | 237.2 | |
| 1641 | BDL-LPS-Yaq-001 | 6.5 | 93.5 | 0.208 | 78 | 283.9 | |
| 1656 | BDL-LPS-Yaq-001 | 9.07 | 115.5 | 0.177 | 78 | | |
| 1719 | BDL-LPS-Yaq-001 | | | | 78.7 | | |
| 1720 | BDL-LPS-Yaq-001 | | | | 79.1 | | |

***Raw data for haemodynamic analysis and microbiological analysis in Experiment 1 and Experiment 3.***

*PP: Portal pressure; MAP: Mean arterial pressure; CPA: Collagen proportionate area; BW: Brain water; NH3A: Arterial ammonia; NH3PV, Portal vein ammonia; A-E: Arterial endotoxin; PV-E: Portal vein endotoxin; A-DNA: Arterial bacterial DNA positivity; PV-DNA: Portal vein bacterial DNA positivity.*

| XR code | Group | Survival | PV-L | A-L | A-M | A-N | PV-N | PV-M |
|---|---|---|---|---|---|---|---|---|
| 1582 | Sham | 6 | | | | | | |
| 1612 | Sham | 6 | | | | | | |
| 1613 | Sham | 6 | | | | | | |
| 1614 | Sham | 6 | | | | | | |
| 1631 | Sham | 6 | | | | | | |
| 1632 | Sham | 6 | | | | | | |
| 1633 | Sham | 6 | | | | | | |
| 1634 | Sham | 6 | | | | | | |
| 1714 | Sham | 6 | | | | | | |
| 1715 | Sham | 6 | | | | | | |
| 1721 | Sham | 6 | | | | | | |
| 2309 | Sham | | 0.99 | 0.091696 | 0.2427975 | 0.334845 | 0.170037 | 1.56 |
| 2310 | Sham | | 1.593 | 0.193167 | 0.1802304 | 0.225907 | 0.197483 | 5.68 |
| 2311 | Sham | | 1.21 | 0.126879 | 0.4648476 | 0.581554 | 0.150278 | 0.525 |
| 2312 | Sham | | 0.774 | 0.045501 | 0.1856222 | 0.309259 | 0.112744 | |
| 2313 | Sham | | 1.197 | 0.074451 | 0.1698687 | 0.242267 | 0.111989 | |
| 2254 | Sham-Yaq-001 | 6 | | | | | | |
| 2255 | Sham-Yaq-001 | 6 | | | | | | |

| | | | | | | | | |
|---|---|---|---|---|---|---|---|---|
| 2256 | Sham-Yaq-001 | 6 | | | | | | |
| 2257 | Sham-Yaq-001 | 6 | | | | | | |
| 2335 | Sham-Yaq-001 | 6 | 1.26 | 1.188 | 0.129112 | 0.2396434 | 0.289828 | 0.192746 |
| 2336 | Sham-Yaq-001 | 6 | 1.71 | 1.67 | 0.187608 | 0.4667015 | 0.496707 | 0.196224 |
| 2337 | Sham-Yaq-001 | 6 | 1.638 | 0.855 | 0.072595 | 0.1410271 | 0.273096 | 0.171662 |
| 2338 | Sham-Yaq-001 | 6 | 1.521 | 0.279 | 0.026479 | 0.03133728 | 0.158777 | 0.131567 |
| 2339 | Sham-Yaq-001 | 6 | 1.17 | 0.666 | 0.058615 | 0.1930627 | 0.334601 | 0.139045 |
| 1607 | Sham-Yaq-001 | 6 | | | | | | |
| 1608 | Sham-Yaq-001 | 6 | | | | | | |
| 1609 | Sham-Yaq-001 | 6 | | | | | | |
| 1610 | Sham-Yaq-001 | 6 | | | | | | |
| 1643 | Sham-Yaq-001 | 6 | | | | | | |
| 1644 | Sham-Yaq-001 | 6 | | | | | | |
| 1645 | Sham-Yaq-001 | 6 | | | | | | |
| 3322 | Sham-Yaq-001 | 6 | | | | | | |
| 3323 | Sham-Yaq-001 | 6 | | | | | | |
| 3324 | Sham-Yaq-001 | 6 | | | | | | |
| 3328 | Sham-Yaq-001 | 6 | | | | | | |
| 2248 | BDL | 6 | | | | | | |

| | | | | | | | | |
|---|---|---|---|---|---|---|---|---|
| 2249 | BDL | 6 | | | | | | |
| 2250 | BDL | 6 | | | | | | |
| 2251 | BDL | 6 | | | | | | |
| 2089 | BDL | 6 | | | | | | |
| 2090 | BDL | 6 | | | | | | |
| 2088 | BDL | 6 | | | | | | |
| 1584 | BDL | 6 | | | | | | |
| 1586 | BDL | 6 | | | | | | |
| 1592 | BDL | 6 | | | | | | |
| 1594 | BDL | 6 | | | | | | |
| 1619 | BDL | 6 | | | | | | |
| 1621 | BDL | 6 | | | | | | |
| 1622 | BDL | 6 | | | | | | |
| 1625 | BDL | 6 | | | | | | |
| 2304 | BDL | | 10.854 | 13.725 | 4.987281 | 4.125406 | 3.802135 | 3.1774 |
| 2305 | BDL | | 11.943 | 6.21 | 1.450364 | 2.177176 | 4.408699 | 2.76453 |
| 2306 | BDL | | 14.238 | 10.926 | 5.316985 | 1.917207 | 2.73969 | 7.132725 |
| 2307 | BDL | | 7.43 | 7.63 | 2.165547 | 2.654027 | 2.412603 | 1.186095 |
| 2308 | BDL | | | 5.83 | 0.796145 | 2.189118 | | |

| | | | | | | | | |
|---|---|---|---|---|---|---|---|---|
| 2330 | BDL-Yaq-001 | 6 | 8.28 | 4.32 | 1.167445 | 1.20785 | 1.995944 | 2.297998 |
| 2331 | BDL-Yaq-001 | 6 | 5.76 | 5.382 | 2.200463 | 1.428749 | 1.557838 | 2.384053 |
| 2332 | BDL-Yaq-001 | | 2.277 | 3.906 | 1.095117 | 1.436549 | 0.587384 | 0.759127 |
| 2333 | BDL-Yaq-001 | 6 | 9.756 | 4.374 | 1.308578 | 1.367693 | 2.249129 | 3.536979 |
| 2334 | BDL-Yaq-001 | 6 | | | | | | |
| 1599 | BDL-Yaq-001 | 6 | | | | | | |
| 1600 | BDL-Yaq-001 | 6 | | | | | | |
| 1601 | BDL-Yaq-001 | 6 | | | | | | |
| 1615 | BDL-Yaq-001 | 6 | | | | | | |
| 1616 | BDL-Yaq-001 | 6 | | | | | | |
| 1635 | BDL-Yaq-001 | 6 | | | | | | |
| 1636 | BDL-Yaq-001 | 6 | | | | | | |
| 1637 | BDL-Yaq-001 | 6 | | | | | | |
| 1638 | BDL-Yaq-001 | 6 | | | | | | |
| 2242 | BDL-Yaq-001 | 6 | | | | | | |
| 2243 | BDL-Yaq-001 | 6 | | | | | | |
| 2244 | BDL-Yaq-001 | 6 | | | | | | |
| 2245 | BDL-Yaq-001 | 6 | | | | | | |
| 2095 | BDL-Yaq-001 | 6 | | | | | | |

| | | |
|---|---|---|
| 2096 | BDL-Yaq-001 | 6 |
| 2102 | BDL-Yaq-001 | 6 |
| 1579 | Sham-LPS | 6 |
| 1580 | Sham-LPS | 6 |
| 1581 | Sham-LPS | 4 |
| 1611 | Sham-LPS | 6 |
| 1627 | Sham-LPS | 6 |
| 1628 | Sham-LPS | 6 |
| 1629 | Sham-LPS | 6 |
| 1630 | Sham-LPS | 6 |
| 1722 | Sham-LPS | 3 |
| 1603 | Sham-LPS-Yaq-001 | 6 |
| 1604 | Sham-LPS-Yaq-001 | 6 |
| 1605 | Sham-LPS-Yaq-001 | 6 |
| 1606 | Sham-LPS-Yaq-001 | 6 |
| 1647 | Sham-LPS-Yaq-001 | 6 |
| 1648 | Sham-LPS-Yaq-001 | 6 |
| 1649 | Sham-LPS-Yaq-001 | 6 |
| 1650 | Sham-LPS-Yaq-001 | 6 |

| | | |
|---|---|---|
| 1651 | Sham-LPS-Yaq-001 | 6 |
| 1652 | Sham-LPS-Yaq-001 | 6 |
| 1583 | BDL-LPS | 3 |
| 1587 | BDL-LPS | 2 |
| 1588 | BDL-LPS | 4 |
| 1589 | BDL-LPS | 5 |
| 1590 | BDL-LPS | 4 |
| 1591 | BDL-LPS | 3 |
| 1712 | BDL-LPS | 2 |
| 1713 | BDL-LPS | 1 |
| 1724 | BDL-LPS | 4 |
| 1725 | BDL-LPS | 3 |
| 1726 | BDL-LPS | 6 |
| 2319 | BDL-LPS | 3 |
| 2320 | BDL-LPS | 4 |
| 2321 | BDL-LPS | 2 |
| 2322 | BDL-LPS | 1 |
| 2323 | BDL-LPS | 2 |
| 1595 | BDL-LPS-Yaq-001 | 6 |

| | | |
|---|---|---|
| 1596 | BDL-LPS-Yaq-001 | 4 |
| 1597 | BDL-LPS-Yaq-001 | 5 |
| 1598 | BDL-LPS-Yaq-001 | 6 |
| 1617 | BDL-LPS-Yaq-001 | 3 |
| 1618 | BDL-LPS-Yaq-001 | 6 |
| 1639 | BDL-LPS-Yaq-001 | 6 |
| 1640 | BDL-LPS-Yaq-001 | 6 |
| 1641 | BDL-LPS-Yaq-001 | 6 |
| 1656 | BDL-LPS-Yaq-001 | 4 |
| 1719 | BDL-LPS-Yaq-001 | 6 |
| 1720 | BDL-LPS-Yaq-001 | 6 |

***Raw data for survival and whole blood analysis in Experiment 1, Experiment 2 and Experiment 3.***

*Survival: Time to coma stage after LPS injection or to sacrifice; PV-L: Portal vein leukocyte; A-L: Arterial leukocyte; A-M: Arterial monocyte; A-N: Arterial neutrophil; PV-N: Portal vein neutrophil; PV-M: Portal vein monocyte.*

| XR code | Group | Kupffer | Kupffer LPS | CD43 hi LPS | CD43 lo LPS | CD43 hi | CD43 lo | Neu | Neu LPS |
|---|---|---|---|---|---|---|---|---|---|
| 2246 | Sham | 1.56 | 2.93 | 2.42 | 7.19 | 4.6 | 17.9 | 4.57 | 8.81 |
| 2247 | Sham | 5.68 | 6.53 | 3.31 | 5.84 | 1.83 | 8.57 | 9.02 | 15.9 |
| 2252 | Sham | 0.525 | 1.93 | 2.65 | 6.75 | 1.07 | 5.9 | 5.82 | 6.34 |
| 2091 | Sham | | | 2.61 | 12.4 | 1.64 | 5.15 | 3.46 | 11.3 |
| 2335 | Sham-Yaq-001 | | | 4.6 | 21.7 | 2.17 | 14.4 | 9.49 | 31.8 |
| 2336 | Sham-Yaq-001 | | | 6.84 | 19.3 | 5.23 | 15.5 | 19.1 | 19.9 |
| 2337 | Sham-Yaq-001 | | | 3.83 | 7.87 | 3.4 | 8.33 | 5.44 | 11.4 |
| 2338 | Sham-Yaq-001 | | | | | | | | |
| 2339 | Sham-Yaq-001 | | | | | | | | |
| 2304 | BDL | 7.68 | 10.7 | 27.8 | 43.6 | 6.74 | 15.8 | 9.96 | 58 |
| 2305 | BDL | 7.83 | 9.32 | 35.8 | 67.4 | 6.97 | 24 | 17.8 | 79.5 |
| 2306 | BDL | 6.57 | 7.08 | 27 | 30.6 | 5.97 | 10.8 | 19.1 | 72.4 |
| 2307 | BDL | | | 31.3 | 38.7 | 10.9 | 16.7 | 19.7 | 59.9 |
| 2330 | BDL-Yaq-001 | 3.09 | 3.15 | 14.3 | 33.6 | 4.83 | 12.5 | 17.5 | 66.9 |
| 2331 | BDL-Yaq-001 | 2.86 | 4.2 | 22 | 36.6 | 7.89 | 12.9 | 6.73 | 41.9 |
| 2332 | BDL-Yaq-001 | 9.91 | 6.27 | 14.5 | 15.5 | 2.46 | 6.98 | 14.6 | 53.7 |

| | | | |
|---|---|---|---|
| 2333 | BDL-Yaq-001 | 2.09 | 3.03 |
| 2334 | BDL-Yaq-001 | 2.79 | 1.73 |

***Raw data for ROS generation in cells with or without LPS in Experiment 2.***

*Kupffer: Kupffer cell ROS generation; Kupffer LPS: LPS-induced ROS generation in Kupffer; CD43 hi LPS: LPS-induced ROS generation in CD43 high monocyte; CD43 lo LPS: LPS-induced ROS generation in CD43 low monocytes; CD43 hi: ROS generation in CD43 high monocyte; CD43 lo: ROS generation in CD43 low monocyte; Neu: ROS generation in neutrophil; Neu LPS: LPS-induced ROS generation in neutrophil.*

| XR code | Group | IL-10 | IL-1β | IL-6 | TNF-α |
|---|---|---|---|---|---|
| 1612 | Sham | 0 | 2.5 | 20.479 | 25.151 |
| 1613 | Sham | 0 | 0.68 | 20.479 | 8.091 |
| 1614 | Sham | 0 | 9.28 | 106.8 | 13.429 |
| 1782 | Sham | 0 | 0.68 | 20.479 | 0 |
| 2246 | Sham | 1.46 | 2.5 | 5.851 | 37.866 |
| 2312 | Sham | 0 | 5.96 | 0 | 0 |
| 2247 | Sham | 2.046 | 2.3305 | 28.378 | 44.514 |
| 2252 | Sham | 5.209 | 0.888 | 5.851 | 37.866 |
| 1795 | Sham-Yaq-001 | 0 | 29.76 | 0 | 0 |
| 1796 | Sham-Yaq-001 | 0 | 0.432 | 5.851 | 31.404 |

| | | | | | |
|---|---|---|---|---|---|
| 1797 | Sham-Yaq-001 | 1.46 | 7.62 | 45.74 | 0 |
| 2338 | Sham-Yaq-001 | 0 | 0.68 | 12.922 | 44.514 |
| 2339 | Sham-Yaq-001 | 0 | 5.96 | 0 | 0 |
| 2254 | Sham-Yaq-001 | 2.81 | 0.68 | 20.479 | 31.404 |
| 2255 | Sham-Yaq-001 | 2.81 | 3.317 | 12.922 | 37.866 |
| 2256 | Sham-Yaq-001 | 0 | 0.432 | 5.851 | 31.404 |
| 2336 | Sham-Yaq-001 | 0 | 1.823 | 0 | 31.404 |
| 1584 | BDL | 15.9 | 20.48 | 0 | 0 |
| 1586 | BDL | 6.54 | 12.52 | 0 | 0 |
| 1799 | BDL | 6.54 | 22.04 | 0 | 0 |
| 2249 | BDL | 11.3 | 9.28 | 9.311 | 37.866 |
| 2250 | BDL | 1.46 | 43.4 | 0 | 0 |
| 2308 | BDL | 11.3 | 14.14 | 0 | 0 |
| 1776 | BDL | 0 | 1.344 | 20.479 | 37.866 |
| 1778 | BDL | 1.282 | 43.091 | 36.548 | 37.866 |
| 1799 | BDL | 1.282 | 2.313 | 20.479 | 44.514 |
| 1800 | BDL | 7.678 | 14.235 | 36.548 | 37.866 |
| 2251 | BDL | 0 | 78.935 | 12.922 | 31.404 |
| 3337 | BDL | 0 | 17.691 | 12.922 | 37.866 |

| | | | | | |
|---|---|---|---|---|---|
| 3339 | BDL | 7.678 | 116.844 | 44.946 | 44.514 |
| 1601 | BDL-Yaq-001 | 1.46 | 5.96 | 0 | 0 |
| 1786 | BDL-Yaq-001 | 1.46 | 14.14 | 0 | 31.404 |
| 1790 | BDL-Yaq-001 | 0 | 0.432 | 12.922 | 31.404 |
| 1791 | BDL-Yaq-001 | 30.717 | 15.74 | 12.922 | 31.404 |
| 1793 | BDL-Yaq-001 | 98.46 | 72.9 | 18.14 | 75.26 |
| 2243 | BDL-Yaq-001 | 1.282 | 0.432 | 16.652 | 37.866 |
| 1637 | BDL-Yaq-001 | 1.282 | 6.458 | 44.946 | 37.866 |
| 1785 | BDL-Yaq-001 | 0 | 11.966 | 0 | 25.151 |
| 1787 | BDL-Yaq-001 | 0 | 55.955 | 0 | 25.151 |
| 1788 | BDL-Yaq-001 | 5.205 | 45.59 | 12.922 | 31.404 |
| 1789 | BDL-Yaq-001 | 26.95 | 38.127 | 12.922 | 44.514 |
| 2242 | BDL-Yaq-001 | 4.396 | 24.76 | 0 | 31.404 |
| 2244 | BDL-Yaq-001 | 1.282 | 4.345 | 0 | 31.404 |
| 1627 | Sham-LPS | 70.66 | 223.88 | 974.253 | 201.331 |
| 1628 | Sham-LPS | 86.64 | 123.64 | 907.118 | 192.716 |
| 1629 | Sham-LPS | 98.46 | 255.04 | 2486.3 | 218.749 |
| 1630 | Sham-LPS | 78.68 | 290.02 | 2321.977 | 184.165 |
| 1581 | Sham-LPS | 74.68 | 14.235 | 5.851 | 65.399 |

| | | | | | |
|---|---|---|---|---|---|
| 1611 | Sham-LPS | 23.345 | 1.344 | 0 | 37.866 |
| 1652 | Sham-LPS-Yaq-001 | 7.678 | 286 | 2643.04 | 173.36 |
| 1603 | Sham-LPS-Yaq-001 | 3.597 | 38.127 | 250.651 | 72.634 |
| 1604 | Sham-LPS-Yaq-001 | 25.143 | 44.339 | 1237.501 | 126.298 |
| 1605 | Sham-LPS-Yaq-001 | 32.415 | 263.869 | 152.203 | 245.321 |
| 1647 | Sham-LPS-Yaq-001 | 58.622 | 90.776 | 920.471 | 118.346 |
| 1648 | Sham-LPS-Yaq-001 | 51.035 | 233.047 | 2144.371 | 281.528 |
| 1649 | Sham-LPS-Yaq-001 | 50.092 | 93.425 | 1557.328 | 118.346 |
| 1650 | Sham-LPS-Yaq-001 | 52.925 | 68.526 | 974.253 | 118.346 |
| 1651 | Sham-LPS-Yaq-001 | 37.942 | 171.024 | 3152.308 | 346.838 |
| 1589 | BDL-LPS | 106.28 | 152.34 | 312.82 | 79.989 |
| 1590 | BDL-LPS | 230.56 | 167.06 | 312.82 | 95.032 |
| 2321 | BDL-LPS | 2096.08 | 3293.84 | 56103.06 | 37461.14 |
| 2323 | BDL-LPS | 4961.36 | 2681.26 | 15882.16 | 80881.58 |
| 1583 | BDL-LPS | 41.657 | 18.856 | 62.312 | 44.514 |
| 1587 | BDL-LPS | 93.48 | 64.654 | 167.259 | 65.399 |
| 1588 | BDL-LPS | 3931.847 | 1666.258 | 0 | 20191.978 |
| 1591 | BDL-LPS | 12.773 | 28.359 | 0 | 51.328 |
| 1656 | BDL-LPS-Yaq-001 | 1.46 | 4.24 | 13840.505 | 434.062 |

| | | | | | |
|---|---|---|---|---|---|
| 1595 | BDL-LPS-Yaq-001 | 73.983 | 23.569 | 117.898 | 51.328 |
| 1596 | BDL-LPS-Yaq-001 | 0 | 15.38 | 0 | 37.866 |
| 1597 | BDL-LPS-Yaq-001 | 36.094 | 14.235 | 147.229 | 31.404 |
| 1598 | BDL-LPS-Yaq-001 | 54.82 | 59.52 | 501.164 | 184.165 |
| 1617 | BDL-LPS-Yaq-001 | 1.282 | 56.966 | 20.479 | 44.514 |
| 1618 | BDL-LPS-Yaq-001 | 30.586 | 10.845 | 62.312 | 51.328 |
| 1640 | BDL-LPS-Yaq-001 | 294.831 | 315.393 | 11419.219 | 607.439 |

***Raw data for cytokines.***

***Table S2. Cytokines in BDL and ACLF models.***

| | Median Concentration (IQR) | | | |
|---|---|---|---|---|
| **Cytokine** | **Sham (n=8)** | **Sham-Yaq-001 (n=9)** | **BDL (n=13)** | **BDL-Yaq-001 (n=13)** |
| IL-1β | 2.42 (0.73-5.10) | 1.83 (0.56-6.79) | 17.69 (10.90-43.25) | 14.14 (5.15-41.86) |
| IL-6 | 20.48 (5.81-26.40) | 5.85(0-16.70) | 12.92(0-28.51) | 12.92 (0-14.79) |
| TNF-α | 19.29 (2.02-37.87) | 31.40 (0-34.64) | 37.87 (0-37.87) | 31.40 (28.28-37.87) |
| IL-10 | 0 (0-1.9) | 0 (0-2.14) | 6.54(0.64-9.49) | 1.46 (0.64-16.08) |

| | Median Concentration (IQR) | | | |
|---|---|---|---|---|
| **Cytokine** | **Sham-LPS (n=6)** | **Sham-LPS-Yaq-001 (n=9)** | **BDL-LPS (n=8)** | **BDL-LPS-Yaq-001 (n=8)** |
| IL-1β | 173.8 (11.01-263.8) | 93.43 (56.43-248.5) | 159.7 (37.43-2428) | 19.47 (11.69-58.88)* |
| IL-6 | 940.7 (4.39-2363) | 1238 (585.6-2394) | 240 (15.58-11990) | 132.6 (30.94-8690) |
| TNF-α | 188.4(58.52-205.7) | 126.3 (118.3-263.4) | 87.51(54.85-33144) | 51.33 (39.53-371.6) |
| IL-10 | 76.68 (58.83-89.60) | 37.94 (16.41-51.98) | 168.4 (54.61-3471) | 33.34 (1.33-69.19)* |

*$p < 0.05$ compared with BDL-LPS

***Table S3.All DEGs in different organs.***

***Liver: BDL VS Sham***

| Pathway description | Gene list |
|---|---|
| Cytokine-cytokine receptor interaction | Cxcl16/Csf1/Ifngr1/Ccr5/Tgfb2/Cd4/Ccl20/Cxcl12/Il16/Tgfb1/Ccl24/Cxcl6/Ccl2/Tgfb3/Bmp7/Bmp4/Cxcl11/Ccl21/Cx3cl1/Il18/Ccl7/Il1a/Ccr2/Tnfsf10/Ifnar1/Il1b/Cxcl2/Ccr6/Bmp6/Tnf/Ltb/Il1rn/Pf4/Cntf/Cd40/Cxcr3/Il1r1/Il7/Ccl5 |
| Toll-like receptor signaling pathway | Tlr1/Tlr8/Tlr7/Tlr6/Ly96/Cd86/Nfkb1/Tlr9/Cd14/Nfkbia/Cxcl11/Tlr4/Tlr2/Tlr3/Ifnar1/Il1b/Tnf/Tlr5/Irak1/Cd40/Fos/Ccl5 |
| NF-kappa B signaling pathway | Bcl2/Vcam1/Ly96/Cxcl12/Nfkb1/Cd14/Nfkbia/Tlr4/Ccl21/Ptgs2/Il1b/Cxcl2/Icam1/Tnf/Ltb/Ddx58/Irak1/Cd40/Il1r1 |
| Chemokine signaling pathway | Cxcl16/Ccr5/Ccl20/Cxcl12/Ccl24/Nfkb1/Cxcl6/Ccl2/Nfkbia/Jak2/Cxcl11/Ccl21/Cx3cl1/Ccl7/Ccr2/Cxcl2/Ccr6/Pf4/Cxcr3/Ccl5 |
| TNF signaling pathway | Csf1/Vcam1/Ccl20/Nfkb1/Cxcl6/Ccl2/Nfkbia/Cx3cl1/Ptgs2/Il1b/Cxcl2/Icam1/Tnf/Nod2/Fos/Ccl5 |
| NOD-like receptor signaling pathway | Casp1/Bcl2/Nfkb1/Ccl2/Nfkbia/Tlr4/Nlrp3/Il18/Ifnar1/Il1b/Cxcl2/Tnf/Nod2/Ccl5 |
| IL-17 signaling pathway | Ccl20/Nfkb1/Cxcl6/Ccl2/Nfkbia/Ptgs2/Ccl7/Il1b/Cxcl2/Tnf/Fos |
| Necroptosis | Ifngr1/Casp1/Bcl2/Jak2/Tlr4/Nlrp3/Il1a/Tlr3/Tnfsf10/Ifnar1/Il1b/Tnf/Stat6 |
| Th17 cell differentiation | Ifngr1/Cd4/Tgfb1/Nfkb1/Nfkbia/Jak2/Rorc/Il1b/Stat6/Fos/Il1r1 |

| | |
|---|---|
| MAPK signaling pathway | Csf1/Tgfb2/Tgfb1/Nfkb1/Tgfb3/Cd14/Il1a/Pdgfra/Il1b/Tnf/Vegfa/Irak1/Fos/Il1r1 |
| Cytosolic DNA-sensing pathway | Casp1/Nfkb1/Nfkbia/Il18/Il1b/Ddx58/Ccl5 |
| Th1 and Th2 cell differentiation | Ifngr1/Cd4/Nfkb1/Nfkbia/Jak2/Stat6/Fos |
| TGF-beta signaling pathway | Tgfb2/Tgfb1/Tgfb3/Bmp7/Bmp4/Bmp6/Tnf |
| C-type lectin receptor signaling pathway | Casp1/Nfkb1/Nfkbia/Nlrp3/Ptgs2/Il1b/Tnf |
| JAK-STAT signaling pathway | Ifngr1/Bcl2/Jak2/Pdgfra/Ifnar1/Cntf/Stat6/Il7 |
| PI3K-Akt signaling pathway | Csf1/Bcl2/Nfkb1/Jak2/Tlr4/Tlr2/Pdgfra/Ifnar1/Vegfa/Il7 |
| Neutrophil extracellular trap formation | Casp1/Tlr8/Tlr7/Nfkb1/Tlr4/Tlr2/Itgam |
| Apoptosis | Bcl2/Nfkb1/Nfkbia/Tnfsf10/Tnf/Fos |
| Natural killer cell mediated cytotoxicity | Ifngr1/Tnfsf10/Ifnar1/Icam1/Tnf |
| T cell receptor signaling pathway | Cd4/Nfkb1/Nfkbia/Tnf/Fos |
| HIF-1 signaling pathway | Ifngr1/Bcl2/Nfkb1/Tlr4/Vegfa |
| Cell adhesion molecules | Vcam1/Cd4/Cd86/Icam1/Itgam/Cd40 |
| Leukocyte transendothelial migration | Vcam1/Cxcl12/Icam1/Itgam |
| Cellular senescence | Tgfb2/Tgfb1/Nfkb1/Tgfb3/Il1a |
| B cell receptor signaling pathway | Nfkb1/Nfkbia/Fos |

***Liver: BDL-Yaq-001 VS BDL***

| **Pathway description** | **Gene list** |
|---|---|
| Cytokine-cytokine receptor interaction | Ccl24/Il16/Tgfb2/Bmp4/Cxcl16/Ccl4/Ccl17/Ccl20/Bmp6/Ccl3 |
| Toll-like receptor signaling pathway | Tlr3/Tlr1/Irf3/Ccl4/Tlr4/Tlr7/Ccl3 |
| Chemokine signaling pathway | Ccl24/Cxcl16/Ccl4/Ccl17/Ccl20/Ccl3 |
| Cytosolic DNA-sensing pathway | Casp1/Irf3/Ccl4 |
| Necroptosis | Tlr3/Casp1/Tyk2/Tlr4 |
| NOD-like receptor signaling pathway | Casp1/Irf3/Tyk2/Tlr4 |
| TGF-beta signaling pathway | Tgfb2/Bmp4/Bmp6 |
| NF-kappa B signaling pathway | Vcam1/Ccl4/Tlr4 |
| Neutrophil extracellular trap formation | Casp1/Tlr4/Tlr7 |
| IL-17 signaling pathway | Ccl17/Ccl20 |
| TNF signaling pathway | Vcam1/Ccl20 |
| C-type lectin receptor signaling pathway | Casp1/Ccl17 |

***Liver: Sham-Yaq-001 VS Sham***

| Pathway description | Gene list |
|---|---|
| Toll-like receptor signaling pathway | Fos/Tlr7/Cd14/Ccl5 |
| Cytosolic DNA-sensing pathway | Casp1/Ccl5 |
| MAPK signaling pathway | Fos/Cd14/Tgfb1 |
| Th17 cell differentiation | Fos/Tgfb1 |
| TNF signaling pathway | Fos/Ccl5 |
| NOD-like receptor signaling pathway | Casp1/Ccl5 |
| Neutrophil extracellular trap formation | Casp1/Tlr7 |
| Cytokine-cytokine receptor interaction | Ccl5/Tgfb1 |

***Colon: BDL VS Sham***

| Pathway description | Gene list |
|---|---|
| Cytokine-cytokine receptor interaction | Cxcl1/Tnfsf10/Il15/Cxcl10/Ccl7/Ifngr1/Bmp2/Cxcl16/Ccl20/Il18/Tgfb1 |
| Toll-like receptor signaling pathway | Fos/Irf7/Cxcl10/Tlr4/Ly96/Stat1/Rela |
| TNF signaling pathway | Cxcl1/Fos/Il15/Cxcl10/Ccl20/Rela/Icam1 |
| IL-17 signaling pathway | Cxcl1/Fos/Cxcl10/Ccl7/Ccl20/Rela |

| | |
|---|---|
| Chemokine signaling pathway | Cxcl1/Cxcl10/Ccl7/Cxcl16/Ccl20/Stat1/Rela |
| NOD-like receptor signaling pathway | Cxcl1/Irf7/Tlr4/Stat1/Rela/Il18 |
| NF-kappa B signaling pathway | Cxcl1/Tlr4/Ly96/Rela/Icam1 |
| Th17 cell differentiation | Fos/Ifngr1/Stat1/Rela/Tgfb1 |
| Cytosolic DNA-sensing pathway | Irf7/Cxcl10/Rela/Il18 |
| Th1 and Th2 cell differentiation | Fos/Ifngr1/Stat1/Rela |
| HIF-1 signaling pathway | Tlr4/Vegfa/Ifngr1/Rela |
| Necroptosis | Tnfsf10/Tlr4/Ifngr1/Stat1 |
| Natural killer cell mediated cytotoxicity | Tnfsf10/Ifngr1/Icam1 |
| Apoptosis | Tnfsf10/Fos/Rela |
| MAPK signaling pathway | Fos/Vegfa/Rela/Tgfb1 |
| JAK-STAT signaling pathway | Il15/Ifngr1/Stat1 |
| B cell receptor signaling pathway | Fos/Rela |
| TGF-beta signaling pathway | Bmp2/Tgfb1 |
| T cell receptor signaling pathway | Fos/Rela |

***Colon: BDL-Yaq-001 VS BDL***

| **Pathway description** | **Gene list** |
|---|---|
| Cytokine-cytokine receptor interaction | Cx3cl1/Cxcl10/Il15/Bmp2/Cxcl16/Tnfrsf11b/Bmp6/Cntf/Il1r1/Csf1/Ifnar1/Il7/Bmp4/Il18/Cxcl2 |

| | |
|---|---|
| TNF signaling pathway | Mapk1/Cx3cl1/Cxcl10/Il15/Rela/Ptgs2/Icam1/Fos/Csf1/Cxcl2 |
| IL-17 signaling pathway | Mapk1/Cxcl10/Rela/Ptgs2/Fos/Cxcl2 |
| Toll-like receptor signaling pathway | Tlr4/Mapk1/Cxcl10/Rela/Fos/Ifnar1 |
| NF-kappa B signaling pathway | Tlr4/Rela/Ptgs2/Icam1/Il1r1/Cxcl2 |
| NOD-like receptor signaling pathway | Tlr4/Mapk1/Rela/Ifnar1/Il18/Cxcl2 |
| Chemokine signaling pathway | Mapk1/Cx3cl1/Cxcl10/Rela/Cxcl16/Cxcl2 |
| Th17 cell differentiation | Mapk1/Stat6/Rela/Il1r1/Fos |
| JAK-STAT signaling pathway | Stat6/Il15/Cntf/Ifnar1/Il7 |
| Th1 and Th2 cell differentiation | Mapk1/Stat6/Rela/Fos |
| MAPK signaling pathway | Vegfa/Mapk1/Rela/Il1r1/Fos/Csf1 |
| TGF-beta signaling pathway | Mapk1/Bmp2/Bmp6/Bmp4 |
| HIF-1 signaling pathway | Vegfa/Tlr4/Mapk1/Rela |
| Cytosolic DNA-sensing pathway | Cxcl10/Rela/Il18 |
| B cell receptor signaling pathway | Mapk1/Rela/Fos |
| Natural killer cell mediated cytotoxicity | Mapk1/Icam1/Ifnar1 |
| T cell receptor signaling pathway | Mapk1/Rela/Fos |
| Apoptosis | Mapk1/Rela/Fos |
| Necroptosis | Tlr4/Stat6/Ifnar1 |

***Colon: Sham-Yaq-001 VS Sham***

| Pathway description | Gene list |
|---|---|
| Toll-like receptor signaling pathway | Tlr5/Tlr2/Fos/Rela/Mapk8/Ifnar1/Mapk1 |
| Cytokine-cytokine receptor interaction | Ccl20/Il16/Ccr6/Cxcl13/Cxcl14/Bmp7/Cntf/Tgfb1/Ifnar1 |
| IL-17 signaling pathway | Ccl20/Fos/Rela/Mapk8/Mapk1 |
| Chemokine signaling pathway | Ccl20/Ccr6/Cxcl13/Cxcl14/Rela/Mapk1 |
| Th17 cell differentiation | Fos/Rela/Tgfb1/Mapk8/Mapk1 |
| TNF signaling pathway | Ccl20/Fos/Rela/Mapk8/Mapk1 |
| Apoptosis | Gzmb/Fos/Rela/Mapk8/Mapk1 |
| Th1 and Th2 cell differentiation | Fos/Rela/Mapk8/Mapk1 |
| T cell receptor signaling pathway | Fos/Rela/Mapk8/Mapk1 |
| MAPK signaling pathway | Fos/Rela/Tgfb1/Mapk8/Mapk1 |
| NOD-like receptor signaling pathway | Rela/Mapk8/Ifnar1/Mapk1 |
| B cell receptor signaling pathway | Fos/Rela/Mapk1 |
| TGF-beta signaling pathway | Bmp7/Tgfb1/Mapk1 |
| Natural killer cell mediated cytotoxicity | Gzmb/Ifnar1/Mapk1 |
| C-type lectin receptor signaling pathway | Rela/Mapk8/Mapk1 |
| PI3K-Akt signaling pathway | Tlr2/Rela/Ifnar1/Mapk1 |
| Cellular senescence | Rela/Tgfb1/Mapk1 |

| | |
|---|---|
| Neutrophil extracellular trap formation | Tlr2/Rela/Mapk1 |
| HIF-1 signaling pathway | Rela/Mapk1 |

***Brain: BDL VS Sham***

| **Pathway description** | **Gene list** |
|---|---|
| Cytokine-cytokine receptor interaction | Tgfb2/Il18/Bmp4/Ccr5/Tnfrsf11b/Il23a/Tgfb1 |
| Cytosolic DNA-sensing pathway | Il18/Irf7/Ddx58 |
| Toll-like receptor signaling pathway | Irak1/Irf7/Tlr2 |
| TGF-beta signaling pathway | Tgfb2/Bmp4/Tgfb1 |
| MAPK signaling pathway | Tgfb2/Pdgfra/Irak1/Tgfb1 |
| NOD-like receptor signaling pathway | Il18/Irf7/Nod1 |
| NF-kappa B signaling pathway | Irak1/Ddx58 |
| Th17 cell differentiation | Il23a/Tgfb1 |
| Cell cycle | Tgfb2/Tgfb1 |
| JAK-STAT signaling pathway | Pdgfra/Il23a |

***Brain: BDL-Yaq-001 VS BDL***

| Pathway description | Gene list |
|---|---|
| Cytokine-cytokine receptor interaction | Il18/Tgfb2/Il23a/Ccr5 |
| Cytosolic DNA-sensing pathway | Il18/Irf7 |
| Toll-like receptor signaling pathway | Irf7/Tlr7 |
| NOD-like receptor signaling pathway | Il18/Irf7 |
| Neutrophil extracellular trap formation | Tlr7/Itgam |

***Brain:Sham-Yaq-001 VS Sham***

| Pathway description | Gene list |
|---|---|
| TGF-beta signaling pathway | Bmp4 |
| Cytokine-cytokine receptor interaction | Bmp4 |

***Kidney: BDL VS Sham***

| Pathway description | Gene list |
|---|---|
| Cytokine-cytokine receptor interaction | Tnfrsf11b/Cd4/Bmp7/Cxcl11/Ccl24/Tnfsf10/Cxcl16/Pf4 |
| Toll-like receptor signaling pathway | Irak1/Tlr8/Cxcl11/Tlr7/Fos |

| | |
|---|---|
| Chemokine signaling pathway | Cxcl11/Ccl24/Cxcl16/Pf4 |
| NF-kappa B signaling pathway | Icam1/Irak1/Ptgs2 |
| Th17 cell differentiation | Cd4/Rorc/Fos |
| TNF signaling pathway | Icam1/Fos/Ptgs2 |
| Cell adhesion molecules | Itgam/Icam1/Cd4 |
| Neutrophil extracellular trap formation | Itgam/Tlr8/Tlr7 |

***Kidney:BDL-Yaq-001 VS BDL***

| Pathway description | Gene list |
|---|---|
| Chemokine signaling pathway | Ccl21/Ccl24/Pf4/Stat1/Cxcl11 |
| Cytokine-cytokine receptor interaction | Ccl21/Ccl24/Pf4/Cntf/Cxcl11 |
| Toll-like receptor signaling pathway | Tlr3/Stat1/Cxcl11 |

***Kidney:Sham-Yaq-001 VS Sham***

| Pathway description | Gene list |
|---|---|
| Toll-like receptor signaling pathway | Ticam1/Nfkbia/Tlr7/Rela |

| | |
|---|---|
| Th1 and Th2 cell differentiation | Nfkbia/Ifngr1/Rela |
| NF-kappa B signaling pathway | Ticam1/Nfkbia/Rela |
| Th17 cell differentiation | Nfkbia/Ifngr1/Rela |
| NOD-like receptor signaling pathway | Ticam1/Nfkbia/Rela |
| Cytosolic DNA-sensing pathway | Nfkbia/Rela |
| B cell receptor signaling pathway | Nfkbia/Rela |
| IL-17 signaling pathway | Nfkbia/Rela |
| T cell receptor signaling pathway | Nfkbia/Rela |
| TNF signaling pathway | Nfkbia/Rela |
| C-type lectin receptor signaling pathway | Nfkbia/Rela |
| HIF-1 signaling pathway | Ifngr1/Rela |
| Apoptosis | Nfkbia/Rela |
| JAK-STAT signaling pathway | Ifngr1/Cntf |
| Necroptosis | Ticam1/Ifngr1 |
| Chemokine signaling pathway | Nfkbia/Rela |
| Neutrophil extracellular trap formation | Tlr7/Rela |
| Cytokine-cytokine receptor interaction | Ifngr1/Cntf |

***Table S4.Top 20 or significant DEGs in four organs.***

***Liver: BDL VS Sham***

| Upregulated genes | Downregulated genes |
| --- | --- |
| Cxcl16(C-X-C Motif Chemokine Ligand 16) | Apcs(Amyloid P Component, Serum) |
| Csf1(Colony Stimulating Factor 1) | Mif(Macrophage Migration Inhibitory Factor) |
| Casp1(Caspase 1) | Mbl2( Mannose Binding Lectin 2) |
| Ifngr1( Interferon Gamma Receptor 1) | |
| Ccr5(C-C Motif Chemokine Receptor 5) | |
| Cd68( CD68 Molecule) | |
| Tgfb2( Transforming Growth Factor Beta 2) | |
| Tlr2( Toll Like Receptor 2) | |
| Bcl2(BCL2 Apoptosis Regulator) | |
| Tlr1(Toll Like Receptor 1) | |
| Vcam1(Vascular Cell Adhesion Molecule 1) | |
| Tlr8(Toll Like Receptor 8) | |
| Tlr6( Toll Like Receptor 6) | |
| Ly96(Lymphocyte Antigen 96) | |

| |
|---|
| Tlr7( Toll Like Receptor 7) |
| Cd4(CD4 Molecule) |
| Cd86(CD86 Molecule) |
| Ccl20( C-C Motif Chemokine Ligand 20) |

***Liver: BDL-Yaq-001 VS BDL***

| Upregulated genes | Downregulated genes |
|---|---|
| Ccl24(C-C Motif Chemokine Ligand 24) | Tlr3(Toll Like Receptor 3) |
| Bmp1(Bone Morphogenetic Protein 1) | IL16( Interleukin 16) |
| Irf3( Interferon Regulatory Factor 3) | Casp1(Caspase 1) |
| Tyk2( Tyrosine Kinase 2) | Tgfb2( Transforming Growth Factor Beta 2) |
| Ccl4(C-C Motif Chemokine Ligand 4) | Bmp4(Bone Morphogenetic Protein 4) |
| Ccl17(C-C Motif Chemokine Ligand 17) | Cxcl16(C-X-C Motif Chemokine Ligand 16) |
| Ccl3( C-C Motif Chemokine Ligand 3) | Vcam1(Vascular Cell Adhesion Molecule 1) |
| | Tlr1(Toll Like Receptor 1) |
| | Tlr4( Toll Like Receptor 4) |
| | Tlr7(Toll Like Receptor 7) |

| | |
|---|---|
| | Ccl20(C-C Motif Chemokine Ligand 20) |
| | Bmp6( Bone Morphogenetic Protein 6) |

***Colon: BDL VS Sham***

| Upregulated genes | Downregulated genes |
|---|---|
| Cxcl1( C-X-C Motif Chemokine Ligand 1) | Tnfsf10(TNF Superfamily Member 10) |
| Fos( Fos Proto-Oncogene, AP-1 Transcription Factor Subunit) | Irf7(Interferon Regulatory Factor 7) |
| Ccl7(C-C Motif Chemokine Ligand 7) | IL15(Interleukin 15) |
| Rela(RELA Proto-Oncogene, NF-KB Subunit) | Vegfa(Vascular Endothelial Growth Factor A) |
| Icam1( Intercellular Adhesion Molecule 1) | Cxcl10(C-X-C Motif Chemokine Ligand 10) |
| | Ifngr1( Interferon Gamma Receptor 1) |
| | Bmp2(Bone Morphogenetic Protein 2) |
| | Tlr4(Toll Like Receptor 4) |
| | Gpi(Glucose-6-Phosphate Isomerase) |
| | Cxcl16(C-X-C Motif Chemokine Ligand 16) |
| | Ly96(Lymphocyte Antigen 96) |
| | Ccl20( C-C Motif Chemokine Ligand 20) |

| | |
|---|---|
| | IL18(Interleukin 18) |
| | Tgfb1(Transforming Growth Factor Beta 1) |
| | Stat1(Signal Transducer And Activator Of Transcription 1) |

***Colon:BDL-Yaq-001 VS BDL***

| Upregulated genes | Downregulated genes |
|---|---|
| Vegfa(Vascular Endothelial Growth Factor A) | Mapk1(Mitogen-Activated Protein Kinase 1) |
| Tlr4(Toll Like Receptor 4) | Stat6(Signal Transducer And Activator Of Transcription 6) |
| Cx3cl1(C-X3-C Motif Chemokine Ligand 1) | Rela(RELA Proto-Oncogene, NF-KB Subunit) |
| Cxcl10(C-X-C Motif Chemokine Ligand 10) | Ptgs2(Prostaglandin-Endoperoxide Synthase 2) |
| IL15(Interleukin 15) | Icam1(Intercellular Adhesion Molecule 1) |
| Bmp2(Bone Morphogenetic Protein 2) | Cntf(Ciliary Neurotrophic Factor) |
| Cxcl16(C-X-C Motif Chemokine Ligand 16) | Tnfrsf11b(TNF Receptor Superfamily Member 11b) |
| Bmp6(Bone Morphogenetic Protein 6) | Csf1(Colony Stimulating Factor 1) |
| IL7( Interleukin 7) | IL1r1(Interleukin 1 Receptor Type 1) |
| IL18( Interleukin 18) | Ifnar1(Interferon Alpha And Beta Receptor Subunit 1) |
| | Fos(Fos Proto-Oncogene, AP-1 Transcription Factor Subunit) |

| | |
|---|---|
| | Bmp4(Bone Morphogenetic Protein 4) |
| | Cxcl2(C-X-C Motif Chemokine Ligand 2) |

***Brain: BDL VS Sham***

| Upregulated genes | Downregulated genes |
|---|---|
| Bmp1(Bone Morphogenetic Protein 1) | Tgfb2( Transforming Growth Factor Beta 2) |
| IL23a(Interleukin 23 Subunit Alpha) | IL18(Interleukin 18) |
| | Bmp4(Bone Morphogenetic Protein 4) |
| | Ccr5(C-C Motif Chemokine Receptor 5) |
| | Pdgfra(Platelet Derived Growth Factor Receptor Alpha) |
| | Irak1( Interleukin 1 Receptor Associated Kinase 1) |
| | Tnfrsf11b(TNF Receptor Superfamily Member 11b) |
| | Irf7( Interferon Regulatory Factor 7) |
| | Tlr2(Toll Like Receptor 2) |
| | F3(Coagulation Factor III, Tissue Factor) |
| | Nod1(Nucleotide Binding Oligomerization Domain Containing 1) |

| | |
|---|---|
| | Ddx58(RNA Sensor RIG-I ) |
| | Tgfb1(Transforming Growth Factor Beta 1) |

***Brain: BDL-Yaq-001 VS BDL***

| Upregulated genes | Downregulated genes |
|---|---|
| IL18(Interleukin 18) | Irf7( Interferon Regulatory Factor 7) |
| Tgfb2(Transforming Growth Factor Beta 2) | IL23a(Interleukin 23 Subunit Alpha) |
| Tlr7(Toll Like Receptor 7) | |
| Itgam(Integrin Subunit Alpha M) | |
| Ccr5(C-C Motif Chemokine Receptor 5) | |

***Kidney: BDL VS Sham***

| Upregulated genes | Downregulated genes |
|---|---|
| Tlr7(Toll Like Receptor 7) | Tnfrsf11b(TNF Receptor Superfamily Member 11b) |

| | |
|---|---|
| Itgam(Integrin Subunit Alpha M) | Irak1(Interleukin 1 Receptor Associated Kinase 1) |
| Tlr8(Toll Like Receptor 8) | Bmp7(Bone Morphogenetic Protein 7) |
| Icam1( Intercellular Adhesion Molecule 1) | Rorc(RAR Related Orphan Receptor C) |
| Cd4(CD4 Molecule) | Cxcl11(C-X-C Motif Chemokine Ligand 11) |
| | Ccl24(C-C Motif Chemokine Ligand 24) |
| | Tnfsf10(TNF Superfamily Member 10) |
| | Mx1(MX Dynamin Like GTPase 1) |
| | Cxcl 16(C-X-C Motif Chemokine Ligand 16) |
| | Mif(Macrophage Migration Inhibitory Factor) |

***Kidney: BDL-Yaq-001 VS BDL***

| **Upregulated genes** | **Downregulated genes** |
|---|---|
| Ccl21(C-C Motif Chemokine Ligand 21) | Tlr3( Toll Like Receptor 3) |
| Adipoq(Adiponectin, C1Q And Collagen Domain Containing) | Stat1(Signal Transducer And Activator Of Transcription 1) |
| Ccl24(C-C Motif Chemokine Ligand 24) | Cntf(Ciliary Neurotrophic Factor) |
| Pf4(Platelet Factor 4) | Cxcl 11(C-X-C Motif Chemokine Ligand 11) |

### Itgam(Integrin Subunit Alpha M)

*Table S5. P value for microbiome at genus and family level.*

| Genus | BDL *vs* Sham | BDL-Yaq-001 *vs* Sham | BDL-Yaq-001 *vs* BDL | Sham-Yaq-001 *vs* Sham |
|---|---|---|---|---|
| Acetanaerobacterium | 0.01532119 | 0.00736385 | 0.30668507 | 0.40053126 |
| Acetitomaculum | 0.05247016 | 0.08215265 | 0.89832679 | 0.47117 |
| Acetivibrio | 0.90958328 | 0.35453948 | 0.39136594 | 0.54029137 |
| Acidaminobacter | 0.4404007 | NA | 0.39136594 | NA |
| Aciditerrimonas | NA | 0.4404007 | 0.39136594 | NA |
| Acinetobacter | 0.35453948 | 1 | 0.39136594 | 0.54029137 |
| Adlercreutzia | 0.4404007 | NA | 0.39136594 | NA |
| Aerococcus | NA | 0.4404007 | 0.39136594 | NA |
| Agromonas | 0.35453948 | 0.35453948 | NA | 0.54029137 |
| Akkermansia | 1 | 0.5621673 | 0.47697354 | 0.54029137 |
| Alistipes | 0.15256272 | 0.61707508 | 0.30668507 | 0.58948512 |
| Alkaliflexus | 0.4404007 | NA | 0.39136594 | NA |
| Allobaculum | 0.49892153 | 1 | 0.58520976 | 0.912421 |
| Anaerobacter | NA | 0.4404007 | 0.39136594 | NA |
| Anaerophaga | 0.42948846 | 0.5203168 | 0.94849128 | 0.06656797 |

| | | | | |
|---|---|---|---|---|
| Anaeroplasma | NA | 0.4404007 | 0.39136594 | NA |
| Anaerorhabdus | 0.35453948 | 0.5621673 | 0.17295492 | 0.54029137 |
| Anaerosphaera | 0.4404007 | NA | 0.39136594 | NA |
| Anaerosporobacter | 0.5621673 | 0.5621673 | 1 | 0.54029137 |
| Anaerostipes | 0.07374822 | 0.03805333 | 0.30668507 | 0.91406196 |
| Anaerotruncus | 0.28397677 | 1 | 0.37109337 | 0.91510603 |
| Anaerovorax | 0.65213965 | 0.8157978 | 0.89146737 | 0.63095404 |
| Anoxynatronum | 0.4404007 | NA | 0.39136594 | NA |
| Aquisalibacillus | 0.35453948 | 0.35453948 | NA | 0.54029137 |
| Asaccharobacter | 0.90958328 | 0.35453948 | 0.39136594 | 0.54029137 |
| Bacteroides | 0.07414553 | 0.05378409 | 0.44328852 | 0.16580656 |
| Barnesiella | 0.00340524 | 0.61707508 | 0.02984206 | 0.59403234 |
| Bilophila | NA | 0.4404007 | 0.39136594 | NA |
| Blautia | 0.00562291 | 0.02432984 | 0.39136594 | 0.59403234 |
| Brevibacillus | 0.4404007 | NA | 0.39136594 | NA |
| Brevundimonas | NA | NA | NA | 0.30743417 |
| Butyricicoccus | 0.17473582 | 0.4744429 | 1 | 0.24095467 |

| | | | | |
|---|---|---|---|---|
| Butyricimonas | 0.82553866 | 0.1662058 | 0.23353681 | 0.30610347 |
| Butyrivibrio | 0.28397677 | 0.28397677 | 0.89832679 | 0.33735565 |
| Catenibacterium | 0.4404007 | 0.04479485 | 0.07394617 | NA |
| Catonella | 0.2597512 | 0.01683902 | 0.09633096 | 0.89565209 |
| Cellulosilyticum | NA | 0.4404007 | 0.39136594 | 0.30743417 |
| Clostridium IV | 0.72098486 | 0.03831876 | 0.05528499 | 0.91510603 |
| Clostridium sensu stricto | 0.16270325 | 0.02046163 | 0.30454972 | 0.47117 |
| Clostridium XI | 0.00145409 | 0.00145409 | NA | 0.45554509 |
| Clostridium XlVa | 0.5203168 | 0.5203168 | 1 | 0.59403234 |
| Clostridium XlVb | 0.76377457 | 0.33948494 | 0.79381541 | 0.63095404 |
| Clostridium XVIII | 0.05334908 | 0.05334908 | NA | 0.63095404 |
| Coprobacillus | 0.90958328 | 0.35453948 | 0.39136594 | 0.54029137 |
| Coprococcus | 0.5203168 | 0.43203489 | 0.89832679 | 0.45554509 |
| Dehalobacter | 0.35453948 | 0.35453948 | NA | 0.54029137 |
| Desmospora | 0.35453948 | 1 | 0.39136594 | 0.54029137 |
| Desulfonispora | 1 | 0.35453948 | 0.39136594 | 0.23784041 |

| | | | | |
|---|---|---|---|---|
| Dolosigranulum | 0.35453948 | 0.35453948 | NA | 0.54029137 |
| Dorea | 1 | 0.31664484 | 0.48172068 | 0.0477332 |
| Enterococcus | 0.4404007 | NA | 0.39136594 | NA |
| Enterorhabdus | 1 | 0.35453948 | 0.39136594 | 0.54029137 |
| Escherichia/Shigella | 1 | 0.35453948 | 0.39136594 | 0.54029137 |
| Ethanoligenens | 0.4988949 | 0.4988949 | 1 | 0.28686865 |
| Eubacterium | 0.45246035 | 0.93810564 | 0.33951 | 0.18689377 |
| Facklamia | 0.4404007 | NA | 0.39136594 | NA |
| Faecalibacterium | 0.2115897 | 0.2115897 | 0.93621767 | NA |
| Finegoldia | 0.35453948 | 1 | 0.39136594 | 0.54029137 |
| Flavonifractor | 0.61707508 | 0.5203168 | 0.89832679 | 0.91510603 |
| Fusibacter | 0.35453948 | 0.35453948 | NA | 0.76090673 |
| Garciella | 0.4404007 | NA | 0.39136594 | NA |
| ge | 0.35453948 | 0.33722298 | 0.07541262 | 0.76090673 |
| Gemmiger | 0.69918794 | 0.43234696 | 0.65506528 | 0.76090673 |
| Geopsychrobacter | 0.20151637 | 0.01683902 | 0.44278682 | 0.40053126 |
| Gordonibacter | 0.35453948 | 1 | 0.39136594 | 0.54029137 |

| | | | | |
|---|---|---|---|---|
| Guggenheimella | 0.35453948 | 0.35453948 | NA | 0.54029137 |
| Hallella | 0.51800823 | 0.35045046 | 1 | 0.06656797 |
| Haloplasma | 0.4404007 | 0.4404007 | 1 | NA |
| Holdemania | 0.2115897 | 0.2115897 | 0.93621767 | NA |
| Howardella | 0.5621673 | 0.5621673 | 1 | 0.76090673 |
| Hydrogenoanaerobacterium | 0.48463991 | 0.48463991 | 0.64159157 | 0.71863151 |
| Lachnobacterium | 0.56717624 | 0.01841616 | 0.70147811 | 0.19945761 |
| Lachnospiracea_incertae_sedis | 0.94305667 | 0.83032426 | 0.79829785 | 1 |
| Lactobacillus | 0.72098486 | 0.35311123 | 0.17923391 | 0.74911913 |
| Lactococcus | NA | 0.4404007 | 0.39136594 | NA |
| Lactonifactor | 0.19065754 | 0.19065754 | 1 | 0.47117 |
| Mahella | NA | NA | NA | 0.30743417 |
| Marvinbryantia | 0.02680913 | 0.01241933 | 0.30668507 | 1 |
| Meniscus | 0.35311123 | 0.61707508 | 0.09669757 | 0.59403234 |
| Methylobacillus | NA | NA | NA | 0.30743417 |

| | | | | |
|---|---|---|---|---|
| Methylobacterium | 0.4404007 | NA | 0.39136594 | NA |
| Mitsuokella | 0.90958328 | 0.35453948 | 0.39136594 | 0.54029137 |
| Morganella | 0.4404007 | 0.4404007 | 1 | NA |
| Moryella | 0.2115897 | 0.4404007 | 0.72214814 | NA |
| Mucispirillum | 0.19255509 | 0.01063445 | 0.17489828 | 0.23784041 |
| Odoribacter | 0.35453948 | 0.35453948 | NA | 0.54029137 |
| Oribacterium | 0.5621673 | 0.5621673 | 0.93621767 | 0.54029137 |
| Oscillibacter | 0.28397677 | 0.35311123 | 0.44328852 | 0.74911913 |
| Paludibacter | 0.28397677 | 0.72098486 | 0.07363827 | 0.74911913 |
| Papillibacter | 0.05247016 | 0.24650234 | 0.1598642 | 0.90946395 |
| Parabacteroides | 0.10041249 | 0.00340524 | 0.44328852 | 0.04282561 |
| Paraprevotella | 0.56229003 | 0.05076856 | 0.28856602 | 0.37892286 |
| Parasporobacterium | 0.5621673 | 0.5621673 | 0.81027298 | 0.76090673 |
| Parasutterella | 0.06104832 | 0.14011314 | 0.72214814 | 0.56964559 |
| Parvimonas | 0.1033507 | 0.25673098 | 0.89146737 | 0.05466394 |
| Pasteurella | 1 | 0.35453948 | 0.39136594 | 0.23784041 |
| Peptococcus | 0.66268282 | 0.68385706 | 0.37160385 | 0.03664283 |

| | | | | |
|---|---|---|---|---|
| Peptostreptococcaceae_incertae_sedis | 0.13985412 | 0.13985412 | NA | 0.89565209 |
| Phocaeicola | 0.5621673 | 0.35453948 | 0.17295492 | 0.54029137 |
| Porphyrobacter | 0.4404007 | NA | 0.39136594 | NA |
| Prevotella | 0.03831876 | 0.00822097 | 0.12520103 | 0.59403234 |
| Proteiniclasticum | 0.35453948 | 0.35453948 | NA | 0.54029137 |
| Proteiniphilum | 0.35453948 | 0.35453948 | NA | 0.54029137 |
| Pseudobutyrivibrio | 0.35453948 | 0.35453948 | NA | 0.54029137 |
| Pseudoflavonifractor | 0.35311123 | 0.05378409 | 0.25015301 | 1 |
| Pseudomonas | 0.4404007 | 0.10053369 | 0.20040659 | 0.30743417 |
| Rikenella | 1 | 0.35453948 | 0.39136594 | 0.54029137 |
| Robinsoniella | 0.00822097 | 0.00534099 | 0.44328852 | 0.19945761 |
| Roseburia | 1 | 0.72098486 | 0.20133649 | 1 |
| Roseomonas | 0.4404007 | NA | 0.39136594 | 0.30743417 |
| Ruminococcus | 0.05378409 | 0.35311123 | 0.12520103 | 0.91510603 |
| Saccharofermentans | NA | 0.4404007 | 0.39136594 | 0.30743417 |
| Salinihabitans | 0.01818685 | 0.10053369 | 0.18354212 | NA |

| | | | | |
|---|---|---|---|---|
| Salirhabdus | NA | 0.4404007 | 0.39136594 | NA |
| Sarcina | 0.4404007 | NA | 0.39136594 | NA |
| Serratia | 0.4404007 | 0.4404007 | 1 | 0.09420989 |
| Shuttleworthia | NA | 0.2115897 | 0.17295492 | NA |
| Sphingomonas | 0.35453948 | 0.90958328 | 0.39136594 | 0.54029137 |
| Sporobacter | 0.41895696 | 0.07136199 | 0.5514099 | 0.1236104 |
| Sporobacterium | 0.07410347 | 0.25673098 | 0.79025036 | 0.54029137 |
| Staphylococcus | 0.90958328 | 1 | 1 | 0.54029137 |
| Stenotrophomonas | NA | 0.4404007 | 0.39136594 | NA |
| Streptococcus | 0.35453948 | 0.35453948 | NA | 0.76090673 |
| Streptohalobacillus | 0.69918794 | 0.33722298 | 0.45653718 | 0.54029137 |
| Subdoligranulum | 0.90958328 | 1 | 1 | 0.76090673 |
| Sutterella | 0.13985412 | 0.13985412 | NA | 0.28686865 |
| Syntrophococcus | 0.17473582 | 0.61707508 | 0.30668507 | 1 |
| Tannerella | 0.94305667 | 0.17473582 | 0.09669757 | 0.45554509 |
| Telmatospirillum | 0.35453948 | 0.35453948 | NA | 0.54029137 |
| Terasakiella | 0.62908074 | 0.13985412 | 0.39136594 | 0.40053126 |

| | | | | |
|---|---|---|---|---|
| Thermotalea | 0.84679572 | 0.35453948 | 0.17295492 | 0.54029137 |
| TM7_genera_incerta | | | | |
| e_sedis | 0.00259104 | 0.00543786 | 0.72214814 | 0.91510603 |
| Turicibacter | 0.88624771 | 0.56717624 | 0.40571736 | 1 |
| Veillonella | 0.4404007 | 0.4404007 | 1 | NA |
| Xylanibacter | 0.35453948 | 0.35453948 | NA | 0.54029137 |
| Zhangella | 0.28397677 | 0.5203168 | 0.44328852 | 0.91510603 |

| **Family** | **BDL *vs* Sham** | **BDL-Yaq-001*vs* Sham** | **BDL-Yaq-001 *vs* BDL** | **Sham-Yaq-001 *vs* Sham** |
|---|---|---|---|---|
| Acetobacteraceae | 0.4404007 | NA | 0.39136594 | 0.30743417 |
| Aerococcaceae | NA | 0.4404007 | 0.39136594 | NA |
| Bacillaceae 2 | 0.4404007 | 0.2115897 | 0.53288403 | NA |
| Bacteroidaceae | 0.07335102 | 0.05345511 | 0.4051996 | 0.16452182 |
| Bacteroidales_incertae_sedis | 0.20999967 | NA | 0.17168169 | NA |
| Bradyrhizobiaceae | 0.35453948 | 0.35453948 | NA | 0.54029137 |
| Carnobacteriaceae | NA | 0.4404007 | 0.39136594 | 0.30743417 |
| Clostridiaceae 1 | 0.12043817 | 0.01540553 | 0.3961912 | 1 |

| | | | | |
|---|---|---|---|---|
| Clostridiales_Incertae Sedis XI | 0.11843302 | 0.29415002 | 0.94526394 | 0.0694531 |
| Clostridiales_Incertae Sedis XII | NA | NA | NA | 0.30743417 |
| Clostridiales_Incertae Sedis XIII | 1 | 0.40690323 | 0.45314609 | 0.46421431 |
| Coriobacteriaceae | 0.4404007 | NA | 0.39136594 | NA |
| Cytophagaceae | 0.16816441 | 1 | 0.12270598 | 0.65997959 |
| Deferribacteraceae | 0.20999967 | 0.09967861 | 0.54534967 | 0.09420989 |
| Desulfovibrionaceae | NA | 0.4404007 | 0.39136594 | NA |
| Enterobacteriaceae | NA | 0.4404007 | 0.39136594 | 0.30743417 |
| Erysipelotrichaceae | 0.88367297 | 0.71293781 | 0.79335055 | 0.74223373 |
| Eubacteriaceae | 0.29415002 | 0.69627034 | 0.4538293 | 0.0876688 |
| Geobacteraceae | 0.29315467 | 0.02117369 | 0.45126989 | 0.23784041 |
| Hyphomicrobiaceae | 0.56229003 | 0.7164708 | 0.74246327 | 1 |
| Incertae Sedis XI | 0.4404007 | NA | 0.39136594 | NA |
| Lachnospiraceae | 0.22463864 | 0.0318884 | 0.22428921 | 0.91510603 |
| Lactobacillaceae | 0.66206869 | 0.25047813 | 0.17586617 | 0.73299509 |
| Marinilabiaceae | 0.3591194 | 0.3591194 | 1 | 0.20462644 |
| Methylobacteriaceae | 0.4404007 | NA | 0.39136594 | NA |

| | | | | |
|---|---|---|---|---|
| Methylocystaceae | NA | NA | NA | 0.09217699 |
| Methylophilaceae | NA | NA | NA | 0.30743417 |
| Moraxellaceae | 0.35453948 | 1 | 0.39136594 | 0.54029137 |
| Pasteurellaceae | NA | NA | NA | 0.09217699 |
| Peptococcaceae 1 | 0.4404007 | 0.10053369 | 0.22919141 | 0.02509438 |
| Peptostreptococcaceae | 0.00545366 | 0.00545366 | NA | 0.51860502 |
| Porphyromonadaceae | 0.00804566 | 0.51974274 | 0.02966226 | 0.334409 |
| Prevotellaceae | 0.07374822 | 0.02135907 | 0.36844896 | 0.45554509 |
| Pseudomonadaceae | NA | 0.4404007 | 0.39136594 | 0.30743417 |
| Rhodobacteraceae | 0.20999967 | 0.4404007 | 0.47603349 | NA |
| Rikenellaceae | 0.34503338 | 0.88283181 | 0.36251635 | 0.57741616 |
| Ruminococcaceae | 0.09994538 | 0.08561168 | 0.79786271 | 0.59403234 |
| Staphylococcaceae | NA | 0.4404007 | 0.39136594 | NA |
| Sutterellaceae | 0.01825586 | 0.04980852 | 0.39136594 | 0.16941504 |
| TM7 | 0.01754448 | 0.01754448 | NA | 1 |
| Verrucomicrobiaceae | 0.4404007 | 0.20999967 | 0.59058933 | NA |
| Xanthomonadaceae | NA | 0.4404007 | 0.39136594 | NA |
